# Rewiring native post-transcriptional carbon regulators to build multi-layered genetic circuits and optimize engineered microbes for bioproduction

**DOI:** 10.1101/2023.10.04.560922

**Authors:** Trevor R. Simmons, Gina Partipilo, Anna C. Stankes, Rashmi Srivastava, Ryan Buchser, Darian Chiu, Benjamin K. Keitz, Lydia M. Contreras

## Abstract

As the bioprocessing and synthetic biology spaces rapidly expand, post-transcriptional regulation is emerging as driver for maximizing signal response rate and for minimizing cost per protein within cells. Designing robust post-transcriptional control systems that have precise tunability and can achieve diverse regulatory outcomes are paramount to advance the field. Herein, we develop a new approach for engineered post-transcriptional control in bacteria by rewiring a native regulatory system in *Escherichia coli*, the Carbon Storage Regulatory (Csr) Network, to create tunable, complex genetic circuits. First, by co-opting native components of the Csr Network to regulate translation of a target mRNA transcript, we establish a Csr-regulated Buffer Gate. Next, by rationally engineering the interactions between our synthetic construct and the native components of the Csr Network, we expand our original design into a genetic toolbox of 12 Buffer Gates that achieve precise tunability across a 10-fold range of target gene expression. Subsequently, to further regulatory capabilities using this approach, we develop a Csr-regulated NOT Gate through engineering a Csr-activated sequence into our synthetic constructs. We then build upon the Csr Buffer and Not Gates to create post-transcriptional dual input Boolean OR, NOR, AND and NAND Logic Gates, as well as a genetic pulse circuit. As a third step, we demonstrate portability of our Csr-regulated Buffer Gates into three industrially relevant bacteria by recapitulating Buffer Gate activity simply by leveraging the conserved homologous Csr Network in each species. Lastly, as a demonstration of downstream application, we apply our system to a proof-of-concept synthetic mevalonate pathway. Using our engineered constructs, we optimize mevalonate production in *E.* coli resulting in a three-fold increase in production relative to a transcriptionally controlled mevalonate pathway. As a whole, we establish a novel approach to rewire post-transcriptional regulatory networks for complex bacterial computation that can be utilized for efficient bioproduction in engineered microbes.

## Main

The majority of metabolic engineering approaches rely on overexpressing genetic components that operate independently from their host organism. This dysregulation between engineered and native components imposes metabolic burden and ultimately limits the performance of an engineered system (Glick et al. 1995 *Biotechnol. Adv.*, Kurland et al. 1996 *Mol. Microbiol.*). To address the gap of coordinating gene expression control of both native and non-native genes, the field of dynamic metabolic control (DMC) has proposed the integration of native regulatory networks and synthetic systems such that organisms can holistically tune their own activity (Liu et al. 2018 *JIMB*). There are many examples of successfully leveraging DMC to improve system performance (Farmer et al. 2000 *Nat. Biotechnol.*, Zhang et al. 2012 *Nat. Biotechnol.*, Dahl et al. 2013 *Nat Biotechnol.*, Stevens et al. 2015 *ACS Synth. Biol.*, Dinh et al. 2019 *PNAS*, Gao et al. 2019 *Nat. Comm.*, Zhou et al. 2021 *Metab. Eng.*) to highlight a few. While each of these approaches significantly improved production of a target metabolite, each system was optimized for a singular metabolite or pathway. In accordance with this, the Engineering Biology Research Consortium (EBRC) identified the need to establish genetic toolkits that allows researchers to simply “drag and drop” their target pathway of interest and have that heterologous pathway be synchronized with host regulatory networks is critical to advance the future of synthetic biology (Engineering Biology Roadmap (2019)). Moreover, these systems must be tunable and be able to integrate with complex genetic circuitry. Current systems traditionally lack the ability to perform both.

One technique for improving coordination between the host organism and a heterologous system is to replace transcriptionally regulated elements with post-transcriptional substitutes. Implementing post-transcriptionally regulated synthetic systems reduces the cost-per protein for the cell (Frumkin et al. 2017 *Mol. Cell*, Westbrook et al. 2019 *Biotechnol. Bioeng*., Takahashi et al. 2014 *ACS Syn. Biol.*, Ceroni et al. 2018 *Nat. Methods*). Examples of this include RNA-RNA- based regulation; first used as an engineering handle through the development of Riboregulators (Isaacs et al. 2004), which co-opted specific trans-acting noncoding RNAs to thermodynamically release a 5’ UTR hairpin in the 5’ Untranslated Region (5’ UTR) of a target gene that contained the ribosome binding site (RBS). This original system has been optimized and applied in a variety of engineering contexts. (Green et al. 2014 *Cell*, Pardee et al. 2016 *Cell*, Kim et al. (2019) *Nat. Chem. Biol,* Lahiry et al. 2017 *ACS Syn. Bio,* Takahashi et al. 2013 *NAR,* Rodrigo et al. 2012 *PNAS,* Sowa and Vazquez-Anderson et al. 2015 *NAR*, Mihailovic et al. 2018 *Nat. Comm.*). In parallel, RNA-Protein interactions were identified as a potential regulatory node first by the development of engineered intergenic regions in operons to provide targeted control of RNAse E degradation (Smolke et al. 2000 *AEM,* Pfleger et al. 2006 *Nat. Biotechnol*). Additional RNA- Protein systems have been developed primarily by leveraging native RNA-Protein interactions to achieve novel regulatory outcomes in both prokaryotes (Na et al. 2013 *Nat. Biotechnol.*) and eukaryotes (Saito et al. 2007 *Nat Chem. Biol.,* Kennedy et al. 2014 *NAR*). Most recently, both CRISPR-Cas13a and CRISPR-Csm proteins were used to regulate mRNA targets in prokaryotes and eukaryotes through direct mRNA target cleavage or mRNA target binding, respectively (Abudayyeh et al. 2017 *Nature*, Colognori et al. 2022 *Nat. Biotechnol*).

Engineered post-transcriptional regulation continues to remain an area of opportunity for synthetic gene regulation. Developing engineered systems that leverage native post- transcriptional regulatory networks that have well-characterized interactions, are conserved across multiple species, and yield diverse regulatory outcomes provide attractive engineering targets (Pfleger et al. 2006 *Nat. Biotechnol.*, Na et al. 2013 *Nat. Biotechnol.,* Cho et al. 2023 *Nat. Comm.*). An underexplored system in the context of engineered networks is the Carbon Storage Regulatory system (Csr system), also known as the Regulator of Secondary Metabolites (Rsm) system (Romeo and Babitzke 2018 *Microbiol. Spectr.*) as it natively interacts with central carbon metabolism (Morin et al. 2016 *Mol. Microbiol.*), making it an ideal system for metabolic engineering applications.

The Csr Network is a post-transcriptional regulatory cascade conserved throughout bacteria (Vakulskas et al. 2015 *Microbiol. Mol. Biol. Rev.*). The main actuator of the Network is the Carbon Storage Regulatory Protein A (CsrA), a global RNA-Binding Protein (RBP) implicated in regulating dozens of mRNA transcripts, as well as hypothesized to affect hundreds more, either by direct or indirect interactions (Sowa et al. 2017 *NAR*, Leistra et al. 2018 *Sci. Rep.,* Potts et al. 2017 *Nat. Comm.*, Rojano-Nisimura and Simmons et al. 2023 *BioRxiv*). Canonically, CsrA regulates mRNA targets through binding an A(N)GGA motif (“GGA Motif”) within a stem loop of an RNA hairpin in the 5’ untranslated region (5’ UTR) of an mRNA transcript (Dubey et al. 2005 *RNA*); in this scheme the RBS is occluded by CsrA upon binding and prevents translation initiation (Baker et al. 2002 *Mol. Microbiol.).* Recently, CsrA has also been shown to activate translation by destabilizing Shine Dalgaro-sequestering hairpins or blocking RNAse E degradation sites (Renda et al. 2020 *mBio,* Yakhnin et al. 2013 *Mol. Microbiol.*). CsrA is regulated by two sRNAs, CsrB and CsrC, which contain multiple copies of the GGA motif and can bind up to 9 and 5 copies of CsrA, respectively (Weilbacher et al. 2003 *Mol. Microbiol*.) These sRNAs are regulated by a second protein CsrD, which facilitates degradation of the two sRNAs via RNAse E (Leng et al. 2015 *Mol. Microbiol.*). Importantly, the overall topology of this network is well-defined and conserved throughout most *Gammaproteobacteria* as well as other classes of bacteria, therefore the potential regulatory schemes developed in one species could be applied to a larger set of organisms.

Herein, we rewire native regulatory interactions of the Csr Network to build multi-layered, modular genetic circuits and highlight their ability to achieve complex computation across multiple bacteria. Ultimately, we demonstrate improved metabolite production, as exemplified by mevalonate, an essential precursor in the production of terpenoids and other high-value compounds. As a first step, we relied on natively expressed CsrA protein to repress target genes tagged with an engineered cognate 5’ UTR to construct a Buffer Gate in *E. coli.* To activate expression of our synthetic target genes, we overexpress the *csrB* sRNA, which sequesters CsrA disrupting CsrA regulation of our target genes. We rationally engineer a suite of both the 5’ UTRs and the *csrB* sequences to achieve over a 10-fold range of regulation. As a second step, we expand regulatory capabilities of this system by developing a semi-synthetic NOT Gate through engineering a different 5’ UTR that activates upon CsrA interaction. Developing both the Buffer and NOT Gates allows for higher order computation via two-input OR, NOR, AND, and NAND gates. Additionally, we demonstrate complex applications of the full Csr cascade by creating a Csr-regulated genetic pulse. As a third step, we show portability of the Buffer Gates into other bacteria containing a homologous Csr Network to achieve regulation of target genes. Lastly, we apply these tools to optimize mevalonate production using a Csr-regulated construct and benchmark it against an established transcriptionally regulated design. Our Csr-regulated design yields mevalonate titers 3-fold greater than those produced using our in-house transcriptionally regulated system. These results demonstrate the potential of rewiring native posttranscriptional carbon regulation to synthetize effective complex gene circuits for bioproduction in a way that establishes a potential foundational approach for synchronizing native bacterial systems with engineered bioproduction pathways.

## Results

### Developing a Buffer Gate via native CsrA Control

The CsrA protein is expressed throughout the lifetime of a bacterial cell and is known to directly control the translation of dozens of mRNA transcripts (Sowa et al. 2017 *NAR*), as such we chose a design where natively expressed CsrA regulates heterologous target genes which were fused to an engineered 5’ UTR sequence. This enables CsrA to post-transcriptionally repress any target gene by occluding the RBS (Figure 1a – top panel) because the engineered 5’ UTR sequence is based on native 5’ UTR sequence previously confirmed to directly bind CsrA and block ribosome binding. To activate translation of the target gene, the *csrB* sRNA is overexpressed from a plasmid and sequesters CsrA away from the engineered 5’ UTR sequence. Removing CsrA allows ribosome access to the mRNA transcript, resulting in translation initiation (Figure 1a – Bottom Panel). We define this system as a Csr-regulated Buffer Gate (Csr Buffer Gate). To create our synthetic construct, we placed *csrB* under the isopropyl β-D-1- thiogalactopyranoside (IPTG)-inducible P_LlacO_ promoter and our engineered 5’ UTR-target gene fusion under constitutive expression of the P_Con12_ promoter (Adamson et al. 2016 *PNAS*) upstream from the inducible *csrB* to provide sufficient insulation (Fig. 1b). Our engineered 5’ UTR sequence was based upon the native *glgC* 5’ UTR transcript, as its CsrA binding sites are well-documented, the transcript is only bound by CsrA, and it has the greatest *in vivo* fold-repression (10-20-fold) by CsrA (Liu et al. 1995 *J. Bacteriol.*, Baker et al 2002 *Mol Microbiol.*, Sowa et al. 2017, Leistra et al. 2018 *Sci. Rep.*). CsrA preferentially binds the *glgC* 5’ UTR at two primary GGA-motif sites; one in the stem loop, and one in the native ribosome binding site. There is also a GGA-motif directly upstream of the hairpin, as well as an AGAGA motif directly downstream of the hairpin, which act as secondary binding sites for CsrA (Baker et al. 2002 *Mol Microbiol.*, Leistra et al. 2018 *Sci. Rep.*). We selected the *glgC* -61 to -1 sequence relative to the native translation start site for the engineered 5’ UTR scaffold, as it is the minimal sequence containing the hairpin and all CsrA binding sites (Figure 1b). We appended a “TTGGT” five nucleotide (5-nt) spacer to the 3’ end of the *glgC* sequence to create a node for tuning the RBS without having to alter the fourth CsrA binding site that contains the native RBS (Figure 1b). Five nucleotides were the maximum base pair length that did not disrupt the predicted secondary structure, such that the GGA motif remained in the stem loop of the predicted 5’ UTR hairpin structure (Supp Fig. S1).

**Figure 1.**
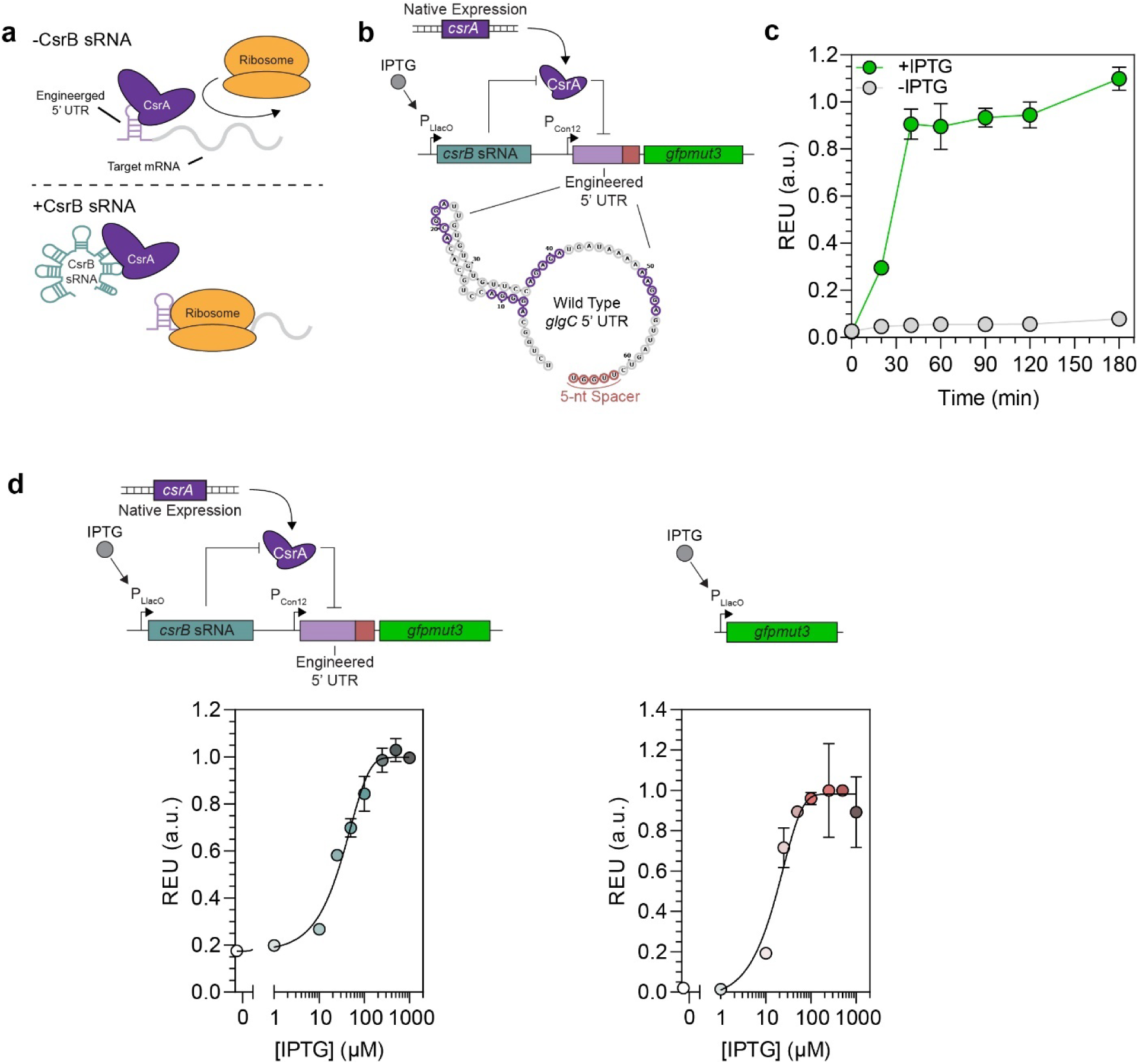
Developing and Benchmarking a Proof-of-concept Csr-Regulated Buffer Gate Circuit using natively available CsrA and plasmid-based CsrB expression: a) Visual representation of the native CsrA and CsrB regulatory roles. In the absence of CsrB, the CsrA protein binds to the engineered 5’ UTR of the target mRNA and blocks translation via occlusion of the ribosome binding site (RBS). Upon induction of CsrB transcription, the sRNA sequesters the CsrA proteins, allowing for translation of our target mRNA. b) Genetic diagram of the Csr-regulated Buffer Gate and predicted secondary structure of the engineered 5’ UTR sequence. We utilized the native *glgC* 5’ UTR with known CsrA binding sites (purple circles) and the 5-nt sequence appended to the 3’ end (red circles). c) Time course of the induced and uninduced Csr- regulated Buffer Gate measured by Relative Expression Units (REU) of GFPmut3. d) Relative fluorescence of GFP expression from the Csr-regulated Buffer Gate (left) and P_LlacO_-controlled Buffer Gate (right) at multiple IPTG induction concentrations.

We tested our flagship design using *gfpmut3* for our gene of interest as it could easily be measured via fluorescence (Fig. 1b). We expected that upon induction of the *csrB* sRNA, it will sequester CsrA away from binding the *glgC* 5’ UTR*-gfp* fusion transcript, allowing it to be translated, and yield an increase in GFP fluorescence. This Csr Buffer Gate, upon *csrB* induction, yielded a rapid increase in signal measured by relative expression units (REU) and reached full activation 40-60 minutes post-induction (Fig. 1c). REU is defined as the OD_600_-normalized fluorescence of a sample containing the engineered 5’ UTR-fluorescent protein fusion transcript divided by the OD_600_-normalized fluorescence of the same construct expressed in a CsrA deletion strain — a constitutive positive control (Moon et al. 2012 *Nature*). The fold-change in fluorescence between the induced and uninduced samples is consistent with native CsrA-*glgC* repression previously measured *in vivo* (Sowa et al. 2017 *NAR*). These results support that the additional 5- nts appended to the *glgC* 5’ UTR do not alter the native interactions between CsrA and the modified *glgC* 5’ UTR sequence. Additionally, growth rates for the induced and uninduced samples were similar (Supp. Fig. S2), demonstrating that, at these levels of *csrB* expression, the CsrA levels required for basal cell viability and growth was not affected; it is worth noting that at higher levels of *csrB* expression, higher levels of CsrA sequestration can induce flocculation or growth defects (Romeo 1998 *Mol. Microbiol.*). We also tested the tunability of the system by titrating *csrB* expression and compared the response to that of *gfpmut3* directly regulated by a P_LlacO_ promoter. Both the Csr-regulated (with scheme shown in Fig. 1b) and the transcriptionally regulated (via P_LlacO_ promoter) Buffer Gates possess similar response functions, demonstrating that regulating heterologous gene expression through native levels of CsrA expression provides similarly tunability to established transcriptional Buffer Gates (Fig. 1d). This confers that native CsrA expression levels are sufficient to maintain essential cellular activity and serve as a regulatory hub for building complex regulatory circuits. Lastly, to confirm that CsrA is responsible for repressing the target gene, we ran the Csr-regulated Buffer Gate in a *csrA* deletion strain and found that there was minimal activation of the Buffer Gate upon *csrB* induction (Supp. Fig. S3)

### Tuning Translation Initiation Rate and CsrA-*csrB* interactions to expand the range of target gene expression

After validating the Csr Buffer Gate, we sought to expand its tunability. In a two-pronged approach, we replaced the wild type *csrB* sRNA sequence with previously engineered mutant sequences (Leistra et al. 2017 *ACS Synth. Biol.*) that varied the *in vivo* affinity between CsrA and the *csrB* sRNA. We also identified additional 5-nt spacer sequence to tune RBS strength of our target genes (Fig. 2a). First, we replaced the WT *csrB* sRNA sequence in the Csr Buffer Gate with the mutant *csrB* sRNAs to evaluate how *csrB*-CsrA affinity affected Buffer Gate response. We selected five *csrB* sequences that previously demonstrated a range of 0.7 to 3.5-fold change *csrB*-CsrA affinity relative to the wild-type *csrB*-CsrA interaction. Of the five *csrB* sRNAs tested, we chose to use the *csrB* L2 and H11 (Leistra et al. 2017 *ACS Synth. Biol.*) as they achieved a 0.5-fold and 1.4-fold change in REU, respectively. The remaining *csrB* mutant sRNAs tested either demonstrated minimal difference in REU relative to the wild type *csrB* sRNA or generated significant growth defects (Supp. Fig. S4). Modulating the affinity between *csrB* and CsrA yielded two additional Csr Buffer Gates with expanded tunability of this system.

**Figure 2.**
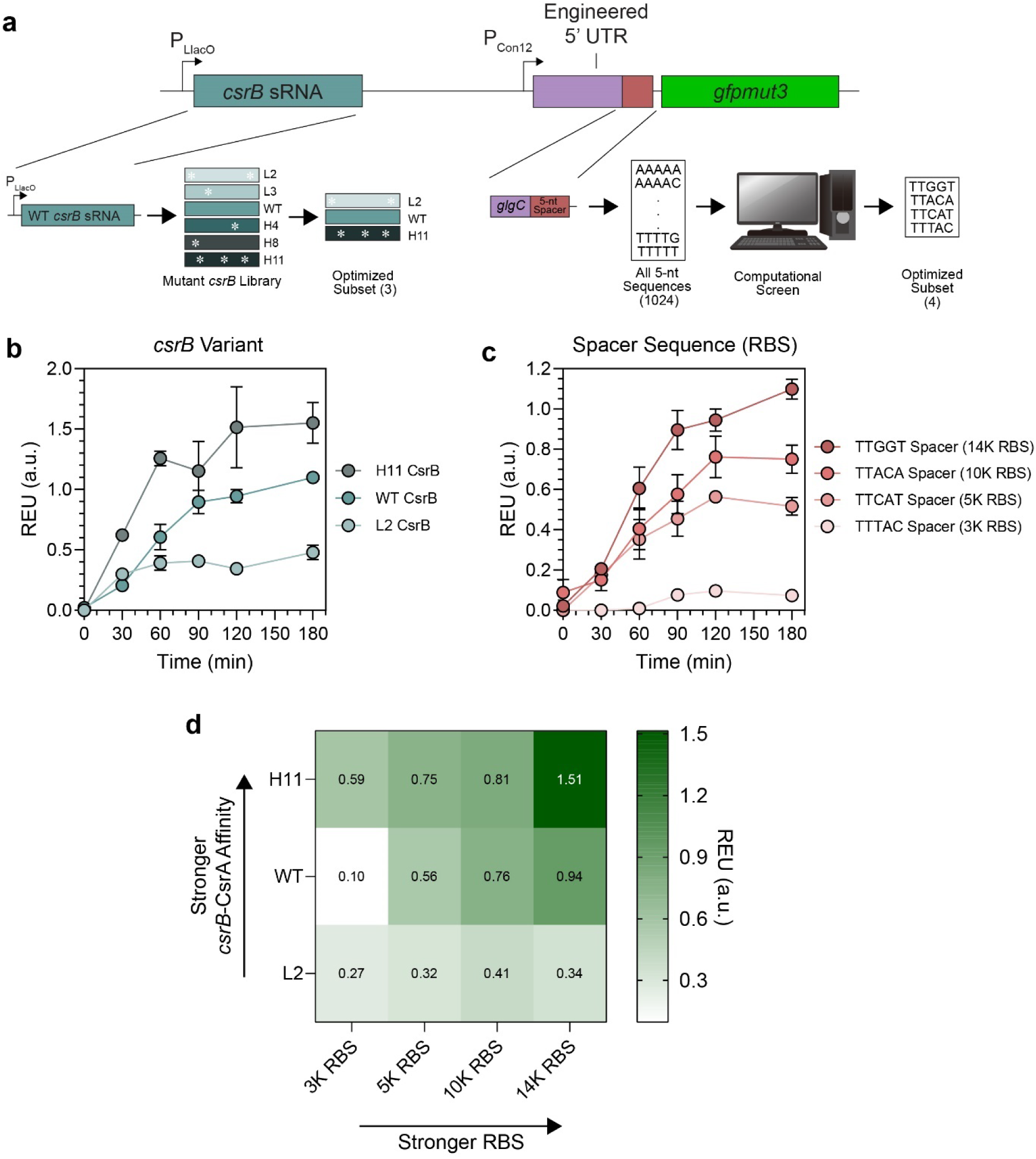
Expanding the dynamic range of gene expression by tuning RBS strength and CsrB-CsrA interaction affinity. a) Workflow for tuning the REU of the Csr-regulated buffer gate, either by tuning the RBS or replacing the WT CsrB sRNA with mutant CsrB. b) REU time course of Csr-regulated Buffer Gates with tunable RBS strengths derived from the variable 5-nt sequence within the engineered 5’ UTR. c) REU time of Csr-regulated Buffer Gates with variable CsrB sRNAs; WT CsrB (Green), High-affinity CsrB (Dark Green), and Low-affinity mutant CsrB L2 (Light Green). d) Heat map of REU for all Csr-regulated Buffer Gate combinations two hours post- induction

Next, to expand target gene translation tunability we sought to identify new 5-nt sequences to vary the RBS. As such, we identified 1023 additional candidate sequences from each possible combination of five nucleotide (i.e., AAAAA to TTTTT or 4^5^ combinations) appended on the 3’ end of the *glgC* 5’ UTR sequence, to replace the “TTGGT” sequence. We removed any transcripts that introduced a new potentially competing GGA binding motif and then screened the remaining pool for their predicted RBS strength via the Salis RBS Calculator 2.0 (Salis 2011 *Meth. Enzymol.*). We selected only the sequences that generated non-zero predicted Translation Initiation Rates (TIRs). From this subset, we evaluated predicted secondary structure of each construct using RNAfold from ViennaRNA (http://rna.tbi.univie.ac.at/cgi-bin/RNAWebSuite/RNAfold.cgi) and selected only the candidates that did not change in secondary structure compared to the wild type *glgC* 5’ UTR sequence. From this final group of 65 sequences, we screened four sequences that maximized the range of predicted RBS strengths (Fig. 2c). Three of the four constructs tested generated measurable GFP expression when used in the Csr-regulated Buffer Gate that correlated with predicted RBS strength (Fig. 2c). One of the four 5-nt spacers did not generate detectable GFP signal. Importantly, each working constructed demonstrated the same 10-fold signal activation upon induction as observed in the original Csr-regulated Buffer Gate, reinforcing that the new spacers in the transcript do not alter the CsrA-5’ UTR interactions (Fig 2c, Supp. Fig. S5).

Finally, we combined the mutant *csrB* sRNAs and the variable RBS strength 5’UTR sequences to create a library of 12 Csr-regulated Buffer Gate constructs that achieve a 10-fold range in target gene expression (Fig. 2d, Supp. Fig. S6, S7). Inducer concentration, RBS strength, and CsrA-*csrB* affinity can each be used to tune the Csr-regulated Buffer Gates. Most traditional transcriptional Buffer Gates only offers the first two nodes for system tunability.

### Evaluating multi-gene regulation via Csr Buffer Gate

Next, we investigated the capabilities of the Csr Buffer Gate for multi-gene regulation. To start, we replaced *gfpmut3* as our gene of interest in the Csr buffer gate (Fig. 1), with *mcherry* and *eyfp* and observed consistent activation upon induction of *csrB* as measured by REU for each of these new gene targets. Activation was consistent for all three genes of interest upon addition of IPTG (Fig. 3a). Next, we evaluated if the Csr-regulated Buffer Gate could control all three genes simultaneously. Each 5’ UTR-fluorescent protein fusion plasmid was transformed into the same strain and the Csr Buffer gate was able to effectively regulate GFP, mCherry, and EYFP simultaneously without changing REU signal for each protein relative to that of the individually regulated protein (Fig. 3b). These results demonstrate that the native CsrA expressed is sufficient to regulate at least three synthetic targets simultaneously, and still perform required cellular functions. We then sought to establish if the system could regulate all three genes in a single synthetic operon. To test this, we constructed a single operon consisting of *gfp*, *mcherry*, and *eyfp* each fused to the original engineered 5’ UTR sequence (Fig. 3c) and expected that CsrA would post-transcriptionally regulate each target gene at each of the 5’ UTR sequences and prevent translation of each protein. To this extent, we observed complete signal activation for each gene in the operon only when the *csrB* sRNA is induced (Fig. 3c). Our results demonstrate that native amounts of CsrA can post-transcriptionally regulate at least three genes by interacting at each engineer 5; UTR sequence simultaneously using this approach.

**Figure 3.**
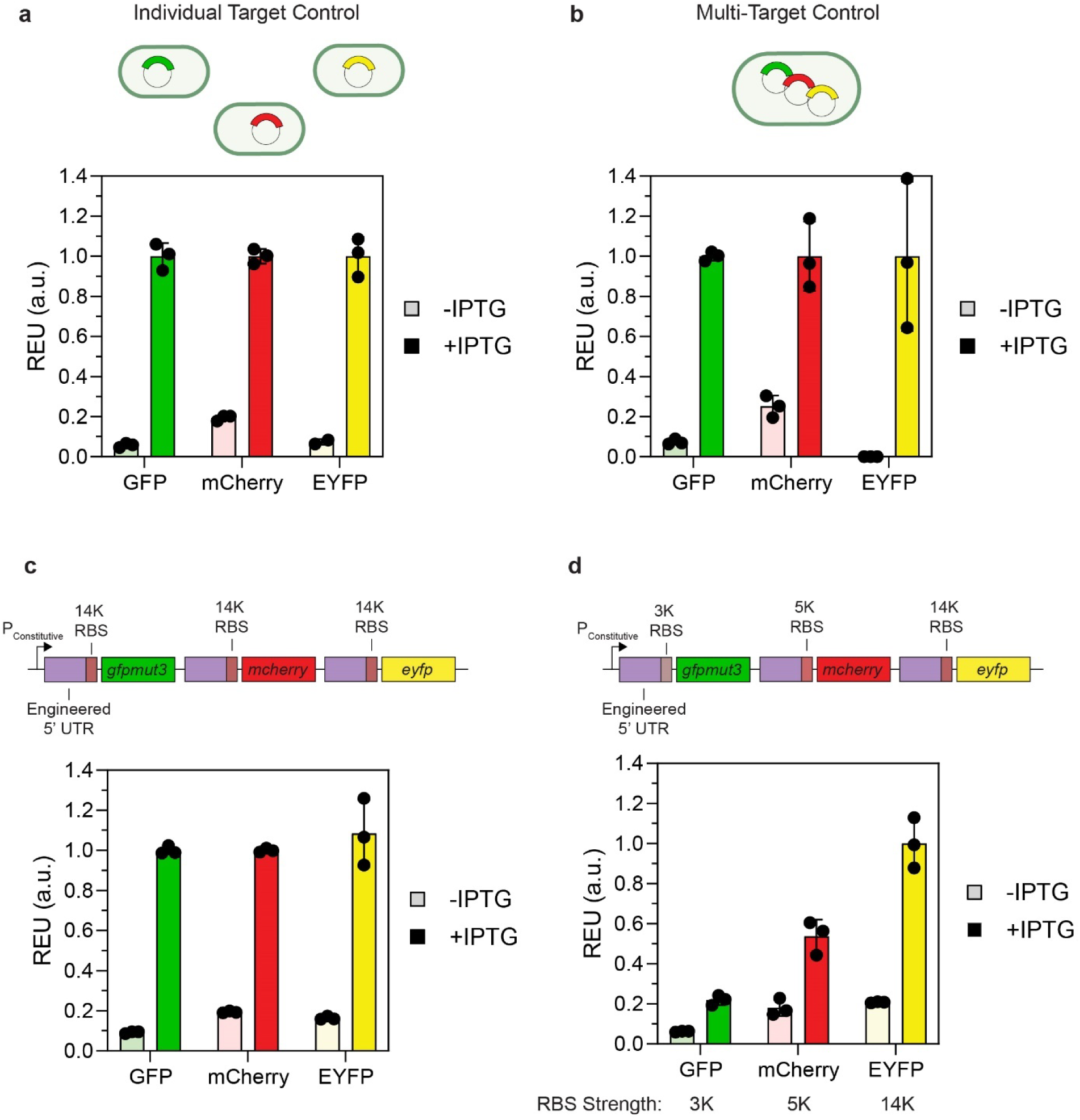
Evaluating regulatory capabilities of the semi-synthetic circuit across multiple genes of interest. A) REU of GFP, mCherry, and EYFP individual Csr-regulated buffer gates two hours post-induction. B) REU of GFP, mCherry, and EYFP Csr-regulated buffer gates in one strain two hours post-induction. C) Diagram for synthetic GFP-mCherry-EYFP operon in the Csr-regulated Buffer Gate. D) REU of GFP, mCherry, and EYFP two hours post-induction regulated simultaneously via the synthetic operon with each gene using the strongest 5-nt spacer. E) REU of GFP, mCherry, and EYFP two hours post-induction. Each gene in the operon had a different RBS strength based on the engineered 5’ UTR selected upstream of each gene.

We then evaluated the tunability of each individual gene within the operon by the engineered 5’ UTRs with different RBS strengths. To test this, we replaced the original engineered 5’ UTR (14K RBS) upstream of *gfpmut3* with the 5’ UTR containing the weakest strength RBS (3K RBS). Concurrently, we replaced the 5’ UTR upstream of *mcherry* with the engineered 5’ UTR of intermediate strength (10K RBS). As in the single transcript case, the REUs observed for GFP and mCherry decreased in proportion to the 5’ UTR RBS strength selected to regulate each respective gene (Fig 3d). This greatly expands the versatility and design space of post-transcriptional operon control, as individual gene expression possesses insulation from the rest of the operon and can be tuned in a predictive fashion.

### Integrating non-canonical CsrA-regulated 5’ UTR creates a Csr-regulated NOT Gate and higher order multi-input Logic Gates

With the engineered 5’ UTR derived from the native *glgC* 5’ UTR, the logic of any gate controlled by Csr system strictly follows that the target gene is repressed by the CsrA protein when the *csrB* sRNA is absent and that the target gene is activated via sequestering the CsrA protein upon *csrB* induction. However, this limits further potential applications to only Buffer Gate-derived systems (e.g., OR, AND, etc.). The inverse logic behavior, where expression of a target gene is activated when the *csrB sRNA* is not induced, would expand the synthetic logical functions available to Csr-regulated circuits. Towards this goal, we identified the *ymdA* 5’ UTR transcript which natively exhibits CsrA-dependent activation upon binding and contains two GGA motifs within its hairpin (Renda et al. 2020 *mBio*). The RBS sequence is sequestered in the stem of the hairpin and inaccessible to the ribosome. When CsrA binds to the *ymdA* hairpin, the hairpin structurally rearranges and releases the RBS, allowing translation initiation (Fig. 4a); that is, in this case, availability of (free) CsrA *activates* expression of the target gene. We evaluated this regulation to reverse the logic of the Csr Buffer Gate by replacing the *glgC* sequence in the engineered 5’ UTR with the -58 to -1 *ymdA* 5’ UTR sequence. When *csrB* is induced and sequesters CsrA from the *ymdA* 5’ UTR, we expect to see a decrease in target protein signal, in this case GFP (Fig. 4b). In our new design, we observed a reduction in GFP signal upon induction of CsrB and release of the hairpin blocking the RBS. The system achieved approximately a 2-fold reduction in REU 60 minutes post-induction (Fig. 4c), establishing a Csr-regulated NOT Gate (Csr NOT Gate).

**Figure 4.**
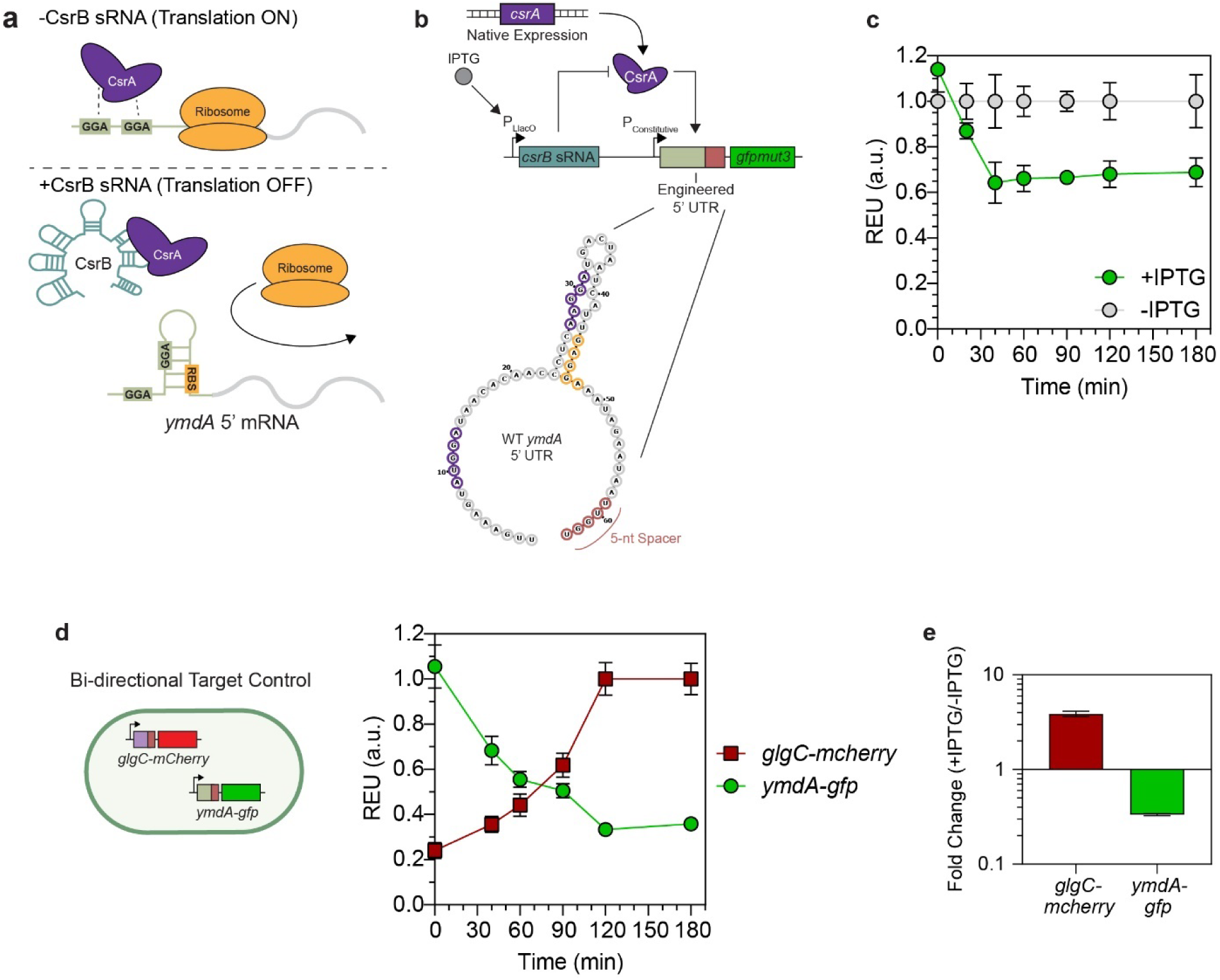
Constructing a Csr-regulated NOT Gate by implementing a new 5’ UTR of inverse regulatory interactions with CsrA and a constructing bi-directional circuit to achieve diverging regulatory outcomes. a) Diagram of the native interaction between the *ymdA* 5’ UTR and CsrA in the absence and presence of CsrB, and the engineered Csr-regulated NOT Gate. b) REU time-course of uninduced and induced Csr-regulated NOT Buffer Gate. c) Diagram of the bi-directional switch. The NOT Gate regulating GFP and the Buffer Gate regulating mCherry are transformed into a single strain. Time course of fold-change in fluorescence between the induced and uninduced system. d) Time course of bi-directional circuit. e) Fold change in regulation for the bi-directional switch two hours post-induction.

Next, we investigated if the Csr NOT Gate (*ymdA* 5’ UTR-regulated) could be used in parallel with the Csr Buffer Gate (*glgC* 5’ UTR-regulated) to achieve bi-directional regulatory outcomes through a single post-transcriptional regulatory hub. To do this, we used the Csr- regulated NOT Gate to control GFP expression and the Buffer Gate to control mCherry expression (Fig. 4d). These constructs were tested multiple Csr deletion strains and we found that the Δ*csrB*Δ*csrC*Δ*csrD (*Δ*BCD) E. coli* strain generated the best response with minimal leakiness (Supp. Fig. S8). In our new testing strain, we observed 4-fold activation of the mCherry signal and 3-fold repression of GFP signal after CsrB induction (Fig. 4d, and Fig. 4e), showing differential regulatory outcomes can be achieved simultaneously based on the CsrA-5’ UTR interaction.

With both the Buffer and NOT Gate established, we expanded to multi-input logic gates by placing *csrB* under the control of multiple inputs to develop Csr-based circuits that can respond to and actuate bacterial computation based on multiple environment signals. First, we constructed the OR logic gate by placing the *csrB* sRNA under both TetR and LacI regulation. We define the Csr-regulated OR Gate as observing signal activation via *csrB* sequestering CsrA away from the engineered *glgC* 5’ UTR (Buffer gate-derived) in the presence of either anhydrotetracycline (aTc) or IPTG, or both compounds simultaneously (Fig 5a). These expected results were established (Fig. 5a) and the OR Gate achieves approximately an 8-fold activation in REU when induced with IPTG, aTc, or both compounds relative to the uninduced condition. We also constructed the NOR logic gate by using the *ymdA* 5’ UTR (NOT gate-derived) while controlling *csrB* expression through the TetR and LacI regulators. The NOR Gate is defined as loss of signal in the presence of either aTc or IPTG or both compounds simultaneously. Again, our expected results are demonstrated *in vivo*, as the Csr-regulated NOR Gate achieves a 3-fold reduction in REU for each active condition (Fig. 5b). We then designed Csr-regulated AND and NAND Gates by placing *csrB* expression under Ara and Tet regulators. We constructed our gates such that *csrB* is downstream of the P_araBAD_ and P_LTetO_ promoters in series, expecting that *csrB* could be expressed only when both L-arabinose (L-ara) and aTc are present. In the absence of one or the other, RNA Polymerase would be sterically hindered from transcription start site directly downstream of the P_LTetO_ promoter. The AND Gate is defined as signal activation only when L-ara and aTc are both present in the samples. The Csr-regulated AND Gate achieves an 8-fold activation in REU when L-ara and aTc are present (Fig. 5c). We note in the aTc only condition there is some activation in signal, which may be due to the fact the araC regulator being unable to fully occlude RNA Polymerase from binding the available P_LTetO_ promoter. For the NAND Gate, we define the logic as loss of signal only in the presence of L-ara and aTc. Using this design, we observe 2-fold reduction in REU only when both L-ara and aTc are present (Fig. 5d). These results demonstrate that Csr-regulated control can be composed to respond to multiple inputs and achieve higher order bacterial computation.

**Figure 5.**
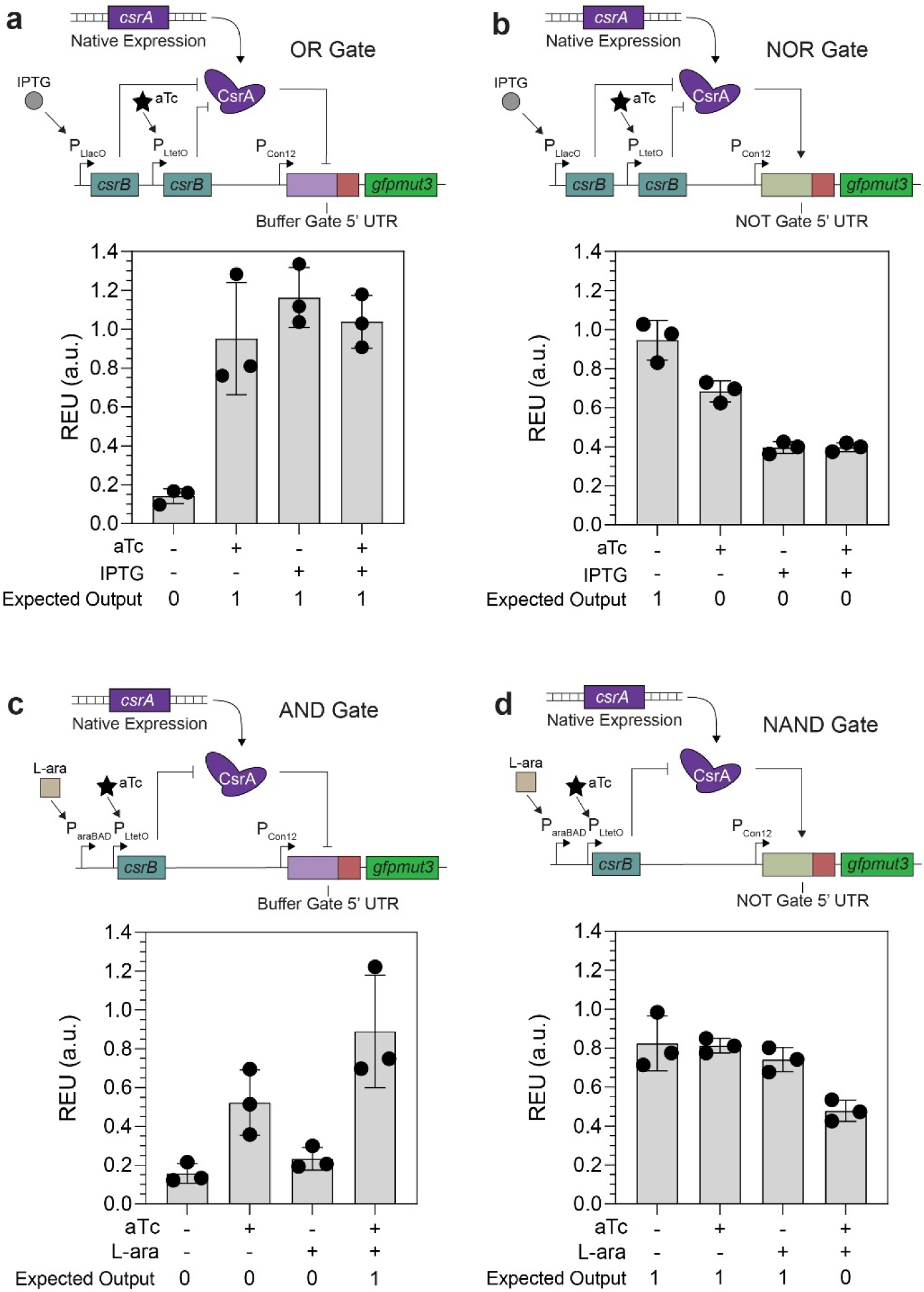
Achieving higher order Csr-regulated Logic Gates by leveraging the Csr- regulated Buffer and NOT Gates. a) Diagram of the Csr-regulated OR Logic Gate under IPTG and aTc control and REU of each condition two hours after induction. The OR Gate uses the Csr- regulated Buffer Gate 5’ UTR to activate *gfpmut3* translation upon IPTG or aTc induction. b) Diagram of the Csr-regulated NOR Logic Gate under IPTG and aTc control and REU of each condition two hours after induction. The NOR Gate uses the Csr-regulated NOT Gate 5’ UTR to activate *gfpmut3* translation upon IPTG or aTc induction. c) Diagram and REU of the Csr- regulated AND Gate under aTc and L-arabinose (L-ara) control two hours post-induction. d) Diagram and REU of the Csr-regulated NAND Gate under aTc and L-ara control two hours post- induction. The expected regulatory outcomes defined for each Logic Gate are either 1, corresponding to *gfpmut3* expression, or 0 corresponding to no *gfpmut3* expression.

Lastly, we created a genetic pulse construct using the Csr-based regulation. To do so, we leveraged the full Csr cascade by using the Csr-regulated Buffer Gate in tandem with a vanillic acid-inducible (Meyer et al. 2019 *Nat. Chem Biol.*) CsrD construct (Extended Fig. 1a and 1b). In this circuit, *csrB* sequesters CsrA away until sufficient CsrD is produced to drive degradation of the *csrB* sRNA and allows for CsrA to re-establish repression of the *gfpmut3* transcript. By inducing the *csrB* sRNA and CsrD protein simultaneously, we demonstrate a Csr-regulated genetic pulse, in which GFP signal was rapidly activated then eliminated without affecting overall cell health (Extended Fig. 1c and 1d). These multi-input gates lay the foundation for constructing additional, more complex logic gates using this Csr-regulated scheme.

### Transporting the Csr Buffer Gate into bacteria containing homologous Csr Network

The Csr Network is conserved across *Gammaproteobacteria* and many other classes of bacteria, particularly through the conservation of homologous CsrA/RsmA proteins (Vakulskas et al. 2015 *Microbiol. Mol. Biol. Rev.*). We therefore wanted to evaluate the modularity of the system and its transferability into organisms beyond *E. coli MG1655.* We also sought to re-establish similar regulatory capabilities by leveraging the conserved CsrA homologs. We first selected *Shewanella oneidensis MR-1* as the bacteria possess unique bioprocessing capabilities of industrial relevance (Partipilo et al. 2022 *ACS Cent. Sci.*, Gao et al. 2023 *bioRxiv*, Zhao et al. 2022 *ACS Synth. Biol.*) and contains a CsrA homolog. While some interactions of the homologous Csr network in *S. oneidensis* have been identified, elucidating complete biochemical mechanisms in this species is an ongoing study in the field (Binnenkade et al. 2011 *PLoS One*). Therefore, if we could recapitulate Csr Buffer Gate activity in MR-1, we suspect that similar patterns would be observed in species with more well-established Csr Networks. To ensure maximum compatibility, we replaced the original P_Con12_ constitutive promoter with ones from the Anderson Library Promoters due to their wide species versatility (Anderson et al. 2010 *J. Biol. Eng.*). As to not alter the stoichiometry between intracellular CsrA and the target mRNA transcript level, we screened several promoters and found that J23107 established the most similar response (Supp. Fig. S9). To test the capabilities of our system, we transformed our new Csr Buffer Gate-containing plasmid into MR-1 and demonstrated GFP tunability by titrating IPTG induction conditions in MR-1 (Fig. 6a), establishing that this regulatory approach is feasible across multiple organisms. This Csr Buffer Gate in MR-1 had a similar response function to a comparable P_LlacO_-regulated *gfpmut3* Buffer Gate and did not exhibit any *csrB-*induction dependent growth defects (Supp. Fig. S10). We then validated that the Csr Buffer Gate in MR-1 was in fact CsrA-regulation dependent. To do so, we sequentially mutated the confirmed GGA motif CsrA binding sites in the Buffer Gate 5’ UTR sequence (Fig. 6b) and screened the response of each mutant. We observed a reduction in regulatory effect with each consecutive binding site mutated. When all sites were mutated, we did not observe any change in GFP fluorescence between the induced and uninduced conditions, reinforcing the homologous CsrA protein regulatory mechanisms are conserved between *E. coli* and MR-1, and the CsrA from *S. oneidensis* can recognize non-native mRNA targets in a predictable fashion (Fig. 6c).

**Figure 6.**
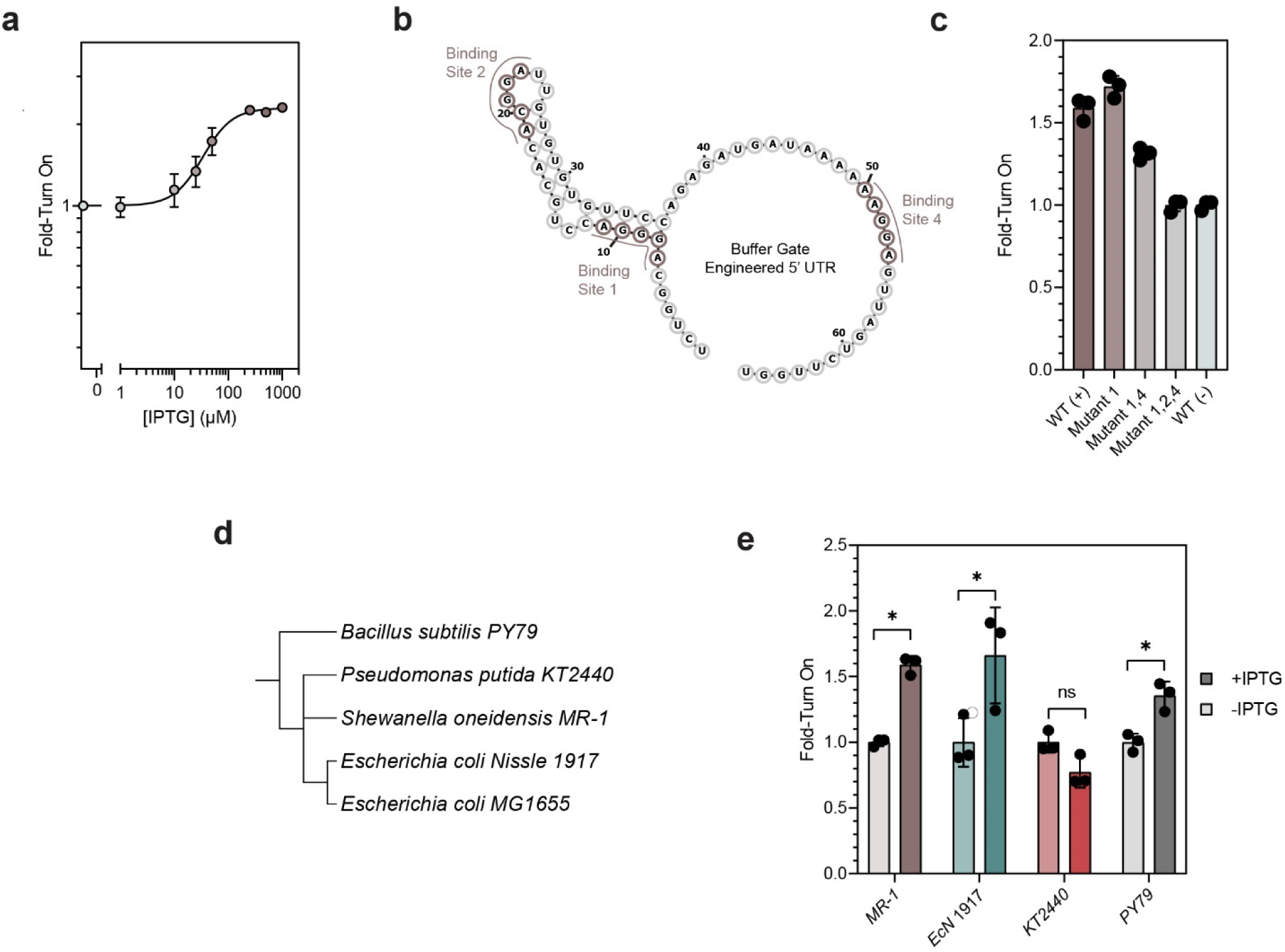
The semi-synthetic Csr Buffer Gate can be ported into multiple bacteria containing homologous bacteria and achieve similar regulation. a) Response function of Csr Buffer Gate in MR-1 at different IPTG induction concentrations. b) Secondary structure of the 5’ UTR used in the Buffer Gate with each GGA binding site identified in tan. c) Fold-Turn on of the Csr-regulated Buffer Gates using the original Buffer Gate 5’ UTR sequence, as well as 5’ UTRs with sequentially mutated CsrA binding sites. d) Phylogenic tree of bacteria tested that contain homologous Csr/Rsm post-transcriptional networks. e) Fold-turn of Csr-Controlled Buffer Gate in each organism. Statistical significance was determined using heteroscedastic t-tests between the induced and uninduced samples, in which the p-value < 0.05.

With the Csr Buffer Gate established in *S. oneidensis* MR-1, we tested the Csr Buffer Gate in *Escherichia coli Nissle 1917* (EcN), and *Pseudomonas putida* KT2440 (KT2440) as these are two bacteria of industrial relevance that contain CsrA homologs (Altegoer et al. 2016 *PNAS*, Praveschotinunt et al. 2019 *Nat. Comm.,* Rottinghaus et al. 2022 *Nat. Comm.*,). We also tested the Csr Buffer Gate in *Bacillus subtilis* PY79 (PY79) given that it is a genetically tractable gram-positive bacterium with a CsrA homologue characterized to regulate mRNA transcripts in a similar fashion (Stork et al. 2021 *Nat. Comm.,* Mukherjee et al. 2011 *Mol. Microbiol.*). Of the three organisms, we observed significant activation upon *csrB* induction in EcN, and PY79, but not in KT2440 (Fig. 6e). Moreover, we measured comparable fold-activation to that of equivalent *lacI- gfpmut3* transcriptionally regulated Buffer Gates in EcN and PY79 (Supp. Fig. S11), demonstrating this post-transcriptional regulatory approach can provide similar regulatory capabilities as previously established tools with the potential similar added advantage of being a basic block for building more complex regulatory logics. Interestingly, we did not see activation of our Csr Buffer Gate in KT2440 despite its similarity to *E. coli*, while we saw activation in PY79, which is more distant genetically (Fig. 6d). We suspect that similar patterns will be observed in additional organisms, as the GGA binding motif has been established for other *E. coli*, *Pseudomonas*, and *Bacillus* species (Sun et al. 2022 *Virulence*, Pourciau et al. 2020 *Front. Microbiol.*). Overall, these results establish that synthetic Csr-regulated systems established in *E. coli* may be able to be ported into bacteria that contain homologous Csr systems and reconstituted by leveraging the conserved functions of the native CsrA proteins.

### Optimizing mevalonate production in *E. coli* via Csr Buffer Gates

We applied our Csr Buffer Gate towards mevalonate production in *E. coli* as a downstream example of a potential bioprocessing application. Given that the Csr network natively drives flux through glycolysis and that expression of the *csrB* sRNA increases accumulation of acetyl-CoA (Morin et al. 2016 *Mol. Microbiol.,* McKee et al. 2012 *Microbial Cell Factories*), we sought to evaluate the ability of our circuits to coordinate synthetic post-transcriptional regulation with native bacterial metabolism; an ultimately success metric was to improve metabolite production relative to other well-established gene regulation systems (e.g. promoter-based LacI-regulation). Mevalonate production in *E. coli* has been previously benchmarked for multiple transcriptional and post-transcriptional synthetic regulatory systems (Martin et al. 2003 *Nat. Biotechnol.*, Pfleger et al. 2006 *Nat. Biotechnol.*, Pitera et al. 2007 *Metab. Eng.*, Shin et al. 2022 *ACS Omega*). Mevalonate is a key precursor in industrial isoprenoid and terpenoid production (Leavell et al. 2016 *Curr. Opin. Biotechnol.*, Kant et al. 2023 *Process Biochem.*) We adapted the previously characterized (Martin et al. 2003 *Nat. Biotechnol.*, Pfleger et al. 2006 *Nat. Biotechnol.*) mevalonate producing synthetic operon (Fig. 7a) and integrated it into the Csr Buffer Gate, such that each gene in the synthetic operon was fused to the engineered *glgC*-derived 5’ UTR (Fig. 7b – pCsr-Mev). To benchmark the efficacy of our system, we selected the synthetic operon from Martin et al. 2003 that is transcriptionally controlled by the LacI regulator (pLac-Mev; Martin et al. 2003 *Nat. Biotechnol.*). As a note, we want to highlight that there have been many significant works towards optimizing mevalonate production in *E. coli* (Ma et al. 2011 *Metab. Eng.*, Wang et al. 2016 *AEM*, Xu et al. 2016 *Bioengineered*). However, most work has primarily focused on strain, carbon source, or protein engineering. Our efforts focus on how co-opting native regulatory components towards a synthetic system affects metabolite production, as such we found that directly comparing mevalonate production to an established synthetic regulatory system would be the most direct.

**Figure 7.**
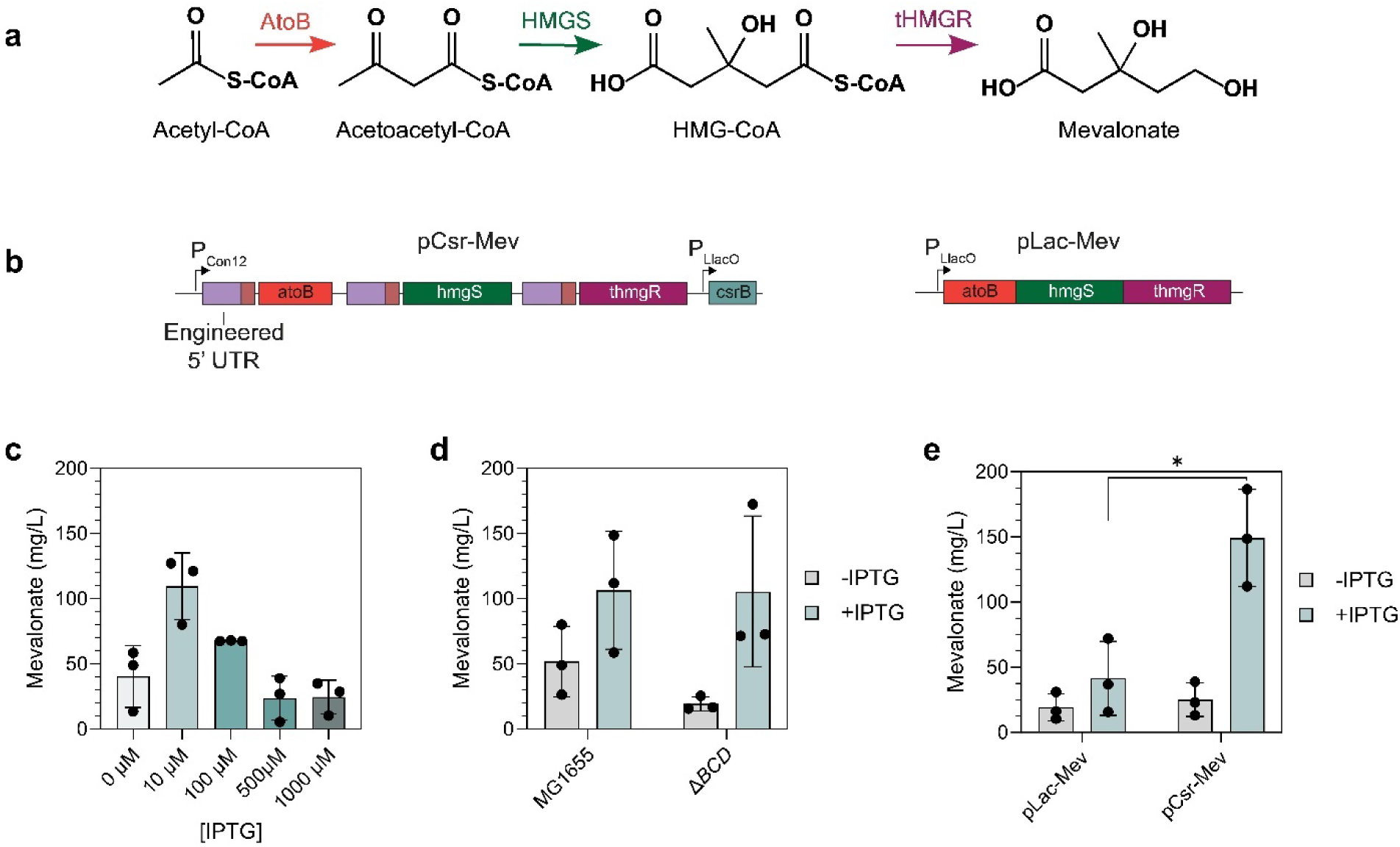
Utilizing the Buffer Gate for mevalonate production outperforms traditional transcriptional-based components in *E. coli*. a) Metabolic pathway for producing mevalonate in *E. coli* (developed in Martin et al. 2003 *Nat. Biotechnol.*). b) Plasmid designs to regulate the synthetic mevalonate operon using a Csr-based regulation (pCsr-Mev) and the original transcriptionally regulated operon (pLac-Mev). c) Final mevalonate titer produced by the pCsr- Mev construct at different IPTG induction concentrations. d) Final mevalonate titers produced by the pCsr-Mev construct in MG1655 and ΔcsrBCD strains of *E. coli.* e) Final mevalonate titers produced by the transcriptionally regulated operon (pLac-Mev) and Csr-regulated operon (pCsr- Mev) after optimization production for both systems. Statistical significance was determined using heteroscedastic t-tests between the induced and uninduced samples, in which the p-value < 0.05.

Initially, we titrated *csrB* induction to find optimal expression conditions and observed that inducing with 10 µM IPTG yielded the greatest mevalonate titer of 110.4 mg/L after 24 hours of culturing (Fig 7c.) Moreover, we observed growth defects in the cells induced with 500 µM IPTG previously established for other Csr-regulated Buffer Gate systems (Supp. Fig. S12a) relative to the cultures induced with 10 µM IPTG. Next, we sought out to reduce the leakiness of the uninduced Csr-regulated mevalonate operon. As we previously observed, utilizing a Δ*csrB*Δ*csrC*Δ*csrD* strain reduced leaky expression in the uninduced logic gates. Porting the mevalonate operon into the Δ*BCD* strain reduced leaky mevalonate production in the uninduced state by approximately 2-fold, while not compromising production for the induced system (Fig. 7d). We also titrated the pLac-Mev system (Martin et al. 2003 *Nat. Biotechnol.*, Pfleger et al. 2006 *Nat. Biotechnol.*, Pitera et al. 2007 *Metab. Eng.*), and found that 100 µM IPTG yielded the maximum mevalonate titer of 56.5 mg/L (Supp. Fig. S13) to ensure each system was fully optimized. We then compared both systems against each other in a single experiment and found that the Csr-regulated system yielded a mevalonate titer of 152.8 mg/L, while the transcriptionally (P_LlacO_-directly) regulated system produced 56.6 mg/L of mevalonate (Fig 7e). To ensure that the difference in titer was not due to increased mRNA expression from our constitutive promoter compared to the inducible P_LlacO_ promoter, we ran a qPCR to quantify the operon mRNA levels in each system. We found that there was no statistical difference in mevalonate operon mRNA levels between the Csr- and LacI-regulated systems (Supp. Fig. S14). With these data in mind, our Csr-regulated system produced approximately a 3-fold greater mevalonate titer, relative to the transcriptionally regulated system.

We postulated that the Csr-regulated system outperformed the transcriptionally regulated circuit because native sequestration of CsrA increases carbon flux through glycolysis and yields a greater accumulation of acetyl-CoA (McKee et al. 2012 *Microbial Cell Factories*). Thus, we thought that due to our semi-synthetic system hinging upon the sequestration of CsrA, we may be favoring glycolysis and acetyl-CoA generation. We then tested mevalonate production in a *csrA::kan* strain, which reduced CsrA activity to 12% of its WT activity (Yakhnin et al. 2013 *Mol. Microbiol.*). Using this strain, we validated that simply deleting the CsrA protein (in either a Csr-regulated system vs. LacI regulated system) results in higher mevalonate yields (Supp. Fig. S15); however, the growth and viability of CsrA mutants are compromised and not viable for scalable bioprocessing (Supp. Fig. S12b and S12c), reinforcing the need to carefully coordinate CsrA sequestration with expression of an engineered metabolic pathways. Overall, this design allows for CsrA-controllable regulation and that post-transcriptional regulation allows for greater titers to be achieved. These results demonstrate that Csr-regulated logic gates could serve as an alternative to traditionally established systems in bioprocessing applications that offers coordination between the synthetically regulated pathway and the native bacterial metabolism.

## Discussion

In this work, we establish a genetic toolbox to leverage the Csr Network from *E. coli* as an engineerable post-transcriptional regulatory hub. As we show in our work, we establish Csr-regulated tunable single input gates, specifically the Buffer (Fig. 1-3) and NOT Gates (Fig 4a-c). We engineered these bi-directional regulatory systems by leveraging two natively occurring (repression and activation) mechanisms that only required CsrA as a single regulator (Fig. 4d and 4e). Using these gates as building blocks we construct higher order bacterial computation systems. Specifically, we established a Csr-regulated NOT Gate by replacing the CsrA-repressed engineered 5’ UTR sequence with a CsrA-activated 5’ UTR sequence derived from the native *ymdA* 5’ UTR in *E. coli* From the Csr Buffer and NOT Gates, we next constructed the OR, NOR, AND, and NAND two-input logic gates (Fig. 5). We then demonstrate the portability of this regulatory approach into other bacteria by recapitulating Csr Buffer Gate activity in *S oneidensis* MR-1, *E. coli Nissle* 1917, and *B. subtilis* PY79 (Fig. 6). Lastly, we demonstrate these circuits can be leveraged to improve target metabolite production by synchronizing control of a synthetic mevalonate operon with the native *E. coli* carbon metabolism (Fig. 7). In this context, we demonstrated that our Csr-regulated Buffer Gate can be applied towards metabolic engineering outcomes and improve mevalonate titer using a synthetic operon relative to that of a transcriptionally regulated counterpart.

Csr-regulated Buffer Gates provide precise control over target gene expression by offering tunability through three primary nodes; inducer concentration, predicted RBS strength, and *csrB-* CsrA affinity. As such this circuit represent additional layer of tunability compared to traditional transcriptionally controlled Buffer Gates. Beyond precise tunability, the Csr-Buffer Gates confer additional benefits relative to previously established Buffer Gate systems. Primarily, we leverage the fact that CsrA is continually expressed throughout the lifetime of the cell (Sowa et al. 2017 *NAR*), thus we rewire the native protein to achieve targeted engineered regulation. Moreover, native levels of CsrA can repress multiple targets simultaneously, while traditional Buffer Gates typically require significantly greater concentrations of regulatory proteins to achieve multi-target repression (e.g., the *lacI^Q^* system (Calos 1978 *Nature*)). Second, target genes in the Csr-Buffer Gate are constitutively transcribed from a plasmid and regulated post-transcriptionally by CsrA. By placing the single regulatory *csrB* sRNA element under the control of a single promoter and utilizing a combinatorial library of 5’ UTRs to control bacterial computation, we endow this system with two advantages: access to a wider number of constitutive promoter libraries, and higher order computation with fewer regulatory elements. Even when three genes were simultaneously expressed (Fig. 3b-d), we did not see increased leaky expression when placing all three genes of interest onto the same plasmid. We postulate that this is due to the single regulatory node controlling multiple genes of interest where the final expression is controlled through the 5’ UTR and RBS as opposed to promoter identity and strength. This additionally allows for rational design of complex regulation without requiring multiple nested promoter and repressor pairs many of which may require expression and maturation of regulatory proteins — causing greater metabolic burden on the cells. Our system controls regulation through one central Lac expression regulating the sRNA *csrB* which can in turn act as a post-transcriptional regulator on multiple, simultaneous target genes. Lastly, we can predictively tune the expression of our target proteins by swapping out the regulatory 5’ UTR element; this can be difficult to achieve using a transcriptionally regulated synthetic operon as varying the RBS of one protein in an operon can affect the translation of the previous protein in the operon. Adding these engineered 5’ UTRs upstream of each target gene provides internal secondary structure within the operon, which has been shown to reduce degradation of synthetic operons in bacteria (Cetnar et al. 2021 *ACS Syn. Biol.*).

The Csr NOT Gate has two main advantages over previously established NOT Gate counterparts. First, most transcriptional logic gates require two nested repressor elements, such as those established in Wang et al. 2011 *Nat. Commun.*, in which the lac system regulates the lambda cI repressor that represses GFP under the P_R_ promoter. The Csr-regulated NOT Gate is directly activated by CsrA target directly activated by CsrA and is repressed by titration of CsrA from the system, no inverter constructs are required. Only recently have transcriptional NOT Gates been developed that do not require inverters (Groseclose et al. 2018 *Nat. Commun.*). Second, post-transcriptional NOT Gates, particularly those derived from toehold switches developed by Green et al. 2014 and Pardee et al. 2014, are greatly limited in sequence tunability as the toehold secondary structure cannot be disrupted for the system to function properly. In the context of the Csr-regulated NOT Gate, the RBS can be directly tuned by varying the 5-nt sequence without disrupting the interaction between the 5’ UTR and CsrA.

The biggest benefit of the Csr-regulated two-input gates is that inverting the gate logic (e.g., OR to NOR, AND to NAND) simply relies on swapping out the engineered 5’ UTR sequence. This means the circuit architecture is identical between each respective gate pair. This has not been yet established in the field of engineered bacteria logic gates, even for the most widely established gates in the synthetic biology community (Nielsen et al. 2016 *Science)*. From a total genetic parts component, the Csr-regulated gates reduce the number of total genetic parts required to achieve similar outcomes for each logic gate, primarily by eliminating the need for nested repressors. Optimized transcriptional two-input logic gates of similar genetic part complexity were recently developed by using several rounds of iterative repressor and anti-repressor engineering in *Escherichia coli* and *Bacteroides* species (Groseclose et al. 2018 *Nat. Commun.*, Huang et al. 2022 *Nat. Commun.*). Csr-regulated logic gates offer a regulatory approach in which many applications can be achieved through a single minimally designed system.

As the Csr Network is conserved across most *Gammaproteobacteria* as well as other classes of bacteria, there are many potential organisms of industrial relevance that could utilize this regulatory scheme (Romeo and Babitzke et al. 2018 *Microbiol. Sprectr.,* Altegeor et al. 2016 *PNAS*). In our work, we demonstrated the Csr-regulated Buffer Gate can be recapitulated in *S. oneidensis* MR-1, *E. coli Nissle 1917* (EcN), and *B. subtilis* (Fig. 6e) and achieve similar levels of fold-regulation to their transcriptional Buffer Gate counterparts (Supp. Fig. S10 and S11). Importantly, we utilized the engineered 5’ UTR sequence as well as the *csrB* sRNA sequenced from *E. coli* and still were able to establish regulation, which alludes to the fact that the natively expressed CsrA homolog in each organism can interact with the non-native transcripts. This is particularly interesting for *B. subtilis*, as no *csrB* sRNA homolog has been experimentally validated. Additionally, through our mutational analysis (Fig. 6c) we demonstrated that the CsrA homolog in *S. oneidensis* appears to follow the same regulatory mechanism of the engineered 5’ UTR to that of the CsrA native to *E.* coli. This highlights that our system offers a predictable regulatory outcome across species. This opens a much larger design space throughout and across bacterial species. This may become relevant when moving into organisms where few promoters are characterized as only one inducible promoter needs to be used to actuate complex logic. Additionally, as these systems rely on native components from a well-conserved network, they may be less likely to fail when ported to another organism, a major concern in *de novo* engineered bacterial circuits (Cardinale et al. 2012 *Biotechnol. J.*, Kittleson et al. 2012 *Curr. Op. in Chem. Biol.*). Conservation of engineered regulatory outcomes has been established using transcriptional regulation through engineered transcription factors (Castillo-Hair et al. 2019 *ACS Synth. Biol.*, Kushwaha et al. 2016 *Nat. Comm.*), as well as CRISPR-Cas-based systems (Gordon et al. 2016 *Metab. Eng.,* Ho et al. 2020 *Mol. Syst. Biol.*, Zhang et al. 2018 *NAR*). Recently, targeted post-transcriptional knockdown of native mRNA targets was established across 12 species of bacteria relying on a synthetic gRNA and the match-making function of the native Hfq protein conserved in each organism (Cho and Yang et al. 2023 *Nat. Comm.*). To our knowledge, our work is the first instance of recapitulating post-transcriptional Buffer Gates on heterologous targets across multiple industrially relevant species.

The fact that these Csr circuits achieve three-fold greater mevalonate titers with the same levels of heterologous mRNAs (Supp. Fig. S14) relative to the P_LlacO_ promoter-controlled version of this engineered pathway indicates the potential of this system to be more efficient from a resource perspective. We hypothesize that using Csr-regulated Buffer Gates for metabolite production natively favors carbon flux through glycolysis towards accumulation of acetyl-CoA pools as when *csrB* is overexpressed, sugars are naturally processed through central carbon metabolism to generate Acetyl-CoA (Morin et al. 2016 *Mol. Microbiol.,* McKee et al. 2012 *Microbial Cell Factories*). During bioprocessing, a greater pool of acetyl-CoA favors improved metabolite titers (Shi et al. 2015 *Curr. Opin. Cell. Biol.*), thus when the Csr-regulated gates are activated more carbon can be shuttled towards acetyl-CoA. This is reflected in the fact that the CsrA deletion strain (*csrA::kan*) containing the Csr-regulated mevalonate operon plasmid produced similar mevalonate titers to that of the wild type and ΔBCD *E. coli* strains with the same plasmid (Supp. Fig. S15). Importantly, we suspect that the synthetic mevalonate operon under transcriptional control does not offer the same effect, as the transcriptional regulator does not interface with any global regulatory proteins.

Overall, we establish a new avenue for engineered post-transcriptional regulation across multiple bacteria. Our system is one of the first semi-synthetic regulatory approaches that offers such a wide range of tunability over single gene expression as well as genes expressed in operon post-transcriptionally. By leveraging the native repression capabilities of CsrA, we reduce the synthetic components required to achieve our desired regulation in our genetic circuits. Moreover, by utilizing both CsrA-repressed and CsrA-activated 5’ UTRs for our engineered target sequences, we can ensure the architecture of each logic gate remains identical regardless of the regulatory outcome or level of complexity. Our demonstration of the improved metabolic yields of desirable compounds upon using native post-transcriptional networks for coupling engineered bioproduction with efficient carbon regulatory processes is also promising. Altogether, this design lays the foundation for rewiring native global RNA-protein regulators, like the Csr Network, as a hub for engineered post-transcriptional regulation that is titratable, portable, and aligned with native bacterial metabolism for efficient bioproduction.

## Materials and Methods

### Bacterial strains and growth conditions

All strains of *E. coli* used in the single- and multi-input gate as well as the mevalonate production experiments are derived from K-12 MG1655. The DH5α strain was used for all plasmid cloning. *S. oneidensis* MR-1, *P. putida* KT2440, *E. coli Nissle* 1917, and *B. subtilis* PY79 were used for the experiments that ported the Csr-regulated Buffer Gate into other bacterial species. The full genotype of each strain used in this work can be found in Table S1. *E. coli* strains used in the single- and multi-input gate were cultured in LB Medium (Miller) (BD Biosciences) supplemented with 100 μg/mL Carbenicillin, and/or 50 μg/mL Kanamycin, and or/ 34 μg/mL Chloramphenicol, as necessary. Seed cultures were grown overnight at 37 °C in an orbital shaking incubator (New Brunswick Scientific *I26*). Cultures were prepared by inoculating 5 mL samples of LB plus the appropriate antibiotics with a single colony of the respective *E. coli* strain grown on LB Agar (Fisher Bioreagents) + appropriate antibiotics. Single- and multi-input gate experiments, overnight cultures were diluted 1:100 into 30 mL LB + antibiotics and grown in the 37 °C shaking incubator. Full protocol for the gate testing is explained below. Experiments testing mevalonate production were prepared by taking an overnight culture and diluted 1:100 into 50 mL LB + 0.2% glycerol (Fisher Bioreagents) and appropriate antibiotics. Samples were also grown in the 37 °C shaking incubator. Full protocol on mevalonate production experiments including extraction is detailed below. All samples were induced with variable concentrations of isopropyl β-D-1- thiogalactopyranoside (IPTG), anhydrotetracycline (aTc), or L-arabinose (L-ara). Stocks of IPTG and were prepared in UltraPure^TM^ Distilled Water (invitrogen), and stocks of aTc were prepared in a 1:1 mixture of UltraPure^TM^ Distilled Water and OmniPur 200 Proof Ethyl Alcohol (Calbiochem, Sigma-Aldrich). Information growth conditions for *S. oneidensis* MR-1, *P. putida* KT2440, *E. coli Nissle* 1917, and *B. subtilis* PY79 can be found in the supplemental materials and methods.

### Plasmid Construction

All plasmids used in this study are provided in Supplementary Table S1. All plasmids were constructed using Gibson Assembly. Oligonucleotide primers and gBlocks were purchased from Integrated DNA Technologies (IDT) and are provided in Supplementary Table S2. Gibson assembly primers were designed as follows: annealing region was designed to include 15-25 nucleotides (nt) of homology to the parent plasmid. The Gibson overhang region was at least 16 nucleotides in length. Primers were designed to minimize GC content, and the Homo-dimer ΔG energy was greater than -18 kcal/mol, as estimated by the IDT OligoAnalyzer (https://www.idtdna.com/calc/analyzer). Plasmids were verified for sequencing by Sanger Sequencing (University of Texas GSAF core) and Plasmidsaurus (https://www.plasmidsaurus.com/). Plasmids derived from previous work was sequence confirmed and analyzed by both Plasmidsaurus and Plannotate (McGuffie,M.J. and Barrick,J.E. 2021 *NAR* http://plannotate.barricklab.org/). It is important to note that the pMP11 plasmid was a generous gift from the lab of Professor Brian Pfleger at the University of Wisconsin-Madison and was used for CRISPR-Cas9-based genome editing of *E. coli* following previously established protocols (Hernández Lozada et al., 2018, Mehrer et al. 2018). Additionally, the pCG004 plasmid was a generous gift from Professor Aditya Kunjapur from the University of Delaware and was a parent plasmid for cloning the Csr-regulated Buffer Gate into a *B. subtilis*-compatible vector.

### Strain Construction

The MG1655 *ΔcsrBΔcsrCΔcsrD E. coli* strain was constructed following the CRISPR-Cas9 protocol previously developed (Mehrer et al. 2018 *Met. Eng.*, and Hernandez Lozada et al. 2018 *ACS Synth. Biol.*). To summarize, the method uses a 2-plasmid system; the first plasmid (sgRNA- *csrD*) contains a constitutively expressed sgRNA that targets the genomic region of *csrD*, the second (pMP11) contains a constitutively expressed *S. pyogenes* Cas9 protein, an L-arabinose-inducible lambda-red recombinase system, an aTc-inducible sgRNA that targets the first plasmid, and a temperature sensitive origin of replication. The csrD gRNA sequence was selected using the CRISPR gRNA Design webtool from Atum (atum.bio). The knockout was made as follows: pMP11 was electroporated (Supplemental Materials and Methods) into an MG1655 *ΔcsrBΔcsrC* strain of *E. coli* from Sowa et al. 2017 and electrocompetent cells were prepared as follows: an overnight starter culture was diluted 1:100 into 50 mL of LB + carbenicillin and grown at 30°C in the shaking incubator. After 1 hour, 2.5 mL of 20% L-arabinose was added to the culture, for a final concentration of 1% L-arabinose, to induce the lambda-red genes. Once the cells reached an OD_600_ of 0.5, they were put on ice, and spun down at 4000 rpm for 10 minutes at 4°C and the supernatant was discarded. The cells were washed with 45 mL of 10% glycerol solution and spun down again at the same conditions. The wash steps were repeated with 25 mL and 10 mL of the 10% glycerol solution. The final pellet was resuspended in 500 μL of 10% glycerol and aliquoted into 50 μL samples. An aliquot of competent cells was mixed with 100 ng of purified pgRNA plasmid and 1 μL of a 60 nt oligo that contains 30 nt upstream of the *csrD* gene and 30 nt downstream of *csrD*. Plasmid and oligo were electroporated and cells were recovered in 900 μL of SOB (Supplemental Materials and Methods) for 3 hours at 30°C and plated on LB Agar plates with appropriate antibiotics. Single colonies were selected the next day and analyzed via cPCR for successful gene knockout. Successful knockouts were grown in an overnight starter culture at 30°C with aTc to cure the pgRNA-*csrD* plasmid. The following day, the cells were grown on LB Agar + antibiotics plate and screened for successful pgRNA curing. Once the pgRNA was cured, single colonies were struck on LB Agar plates and grown overnight at 42°C to cure out pMP11. Individual colonies were screened for curing of pMP11 by streaking colonies on LB Agar and LB Agar + Carbenicillin plates and grown at 30°C overnight.

### Single- and Multi-Input Gate Fluorescence Assays

The response of the Csr-regulated single- and multi-input gates was tested in biological triplicate and each gate was analyzed using cultures grown in 250 mL flasks. To prepare samples, overnight cultures single colonies of the *E. coli* strain containing the Csr-regulated Logic Gate plasmid were inoculated into a 5 mL of LB plus the appropriate antibiotic and grown overnight at 37°C in a shaking incubator. The next day, the overnight culture was diluted 1:100 in 30 mL of LB plus the respective antibiotic. Samples were grown in a shaking incubator at 37°C. After two hours of growth, which corresponded to an OD_600_ between 0.18-0.35, samples were induced with IPTG, aTc, or L-arabinose. For the single-input logic gate experiments, samples were induced with 30 μL of 500 mM IPTG for a final concentration of 500 μM IPTG. For multi-input gate experiments, samples were induced with 30 μL of 500 mM IPTG and 30 μL of 100 μg/mL aTc or 30 μL of 100 μg/mL aTc and 30 μL of 20% L-arabinose. After induction fluorescence and OD_600_ were tracked over time using a BioTek Cytation3 plate reader. At each timepoint, 50 μL of cell culture was diluted into 150 μL of 1x PBS, then fluorescence and OD_600_ were measured. 50 μL of LB was used as a blank. To measure *gfpmut3* fluorescence, excitation and emission wavelengths of 488 nm and 515 nm with a gain of 80. *Mcherry* fluorescence was measured using excitation and emission wavelengths of 585 nm and 613 nm with a gain of 85. *eypf* fluorescence was measured using an excitation and emission wavelengths of 500 nm and 530 nm with a gain of 70. OD_600_ was measuring absorbance using a wavelength of 600 nm.

### Calculating Relative Expression Units (REU)

Relative Expression Units (REU) was calculated by dividing the Fluorescence normalized by OD_600_ (Fl/OD) measured from the samples grown in the testing strain (either MG1655 or MG1655 *ΔcsrBΔcsrCΔcsrD*) by that of the Fl/OD of the samples grown in a *csrA* knockdown strain (*csrA::kan)*. As a note, *csrA* cannot be fully knocked out from the genome, but inserting the *kan* cassette reduced CsrA activity by 85%. The full equation to calculate REU is as follows:

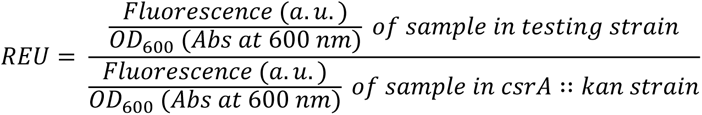

### Mevalonate Production

All mevalonate production experiments were conducted in biological triplicate and cultures were grown in 250 mL flasks. To prepare samples, overnight cultures single colonies of the *E. coli* strain containing the appropriate plasmids were inoculated into a 5 mL of LB plus the appropriate antibiotic and grown overnight at 37°C in a shaking incubator. The next day, the overnight culture was diluted 1:100 in 50 mL of LB supplemented with 0.2% glycerol and the respective antibiotics. Samples were grown in a shaking incubator at 37°C. After two hours of growth, samples were induced with IPTG. 24 hours post-induction, 200 μL of cell culture was sampled for mevalonate quantification. Cell growth was also measured intermittently by tracking OD_600_ over time.

### Mevalonate Extraction and Quantification

Mevalonate extraction was based on the method developed by Rodriguez et al. 2014 *Nat. Protocols.* To summarize, 200 μL of cell culture was sampled from each flask and mixed with 50 μL of 2M HCl (Sigma-Aldrich). Samples were vortexed (Vortex Genie 2) for 15 minutes. 250 μL of 50 μL/mL (-)-trans-caryophyllene (Sigma-Aldric) in ethyl acetate was added to each sample. Samples were vortexed for an additional 5 minutes. Samples were then centrifuged at 3000xg for 7 minutes at room temperature. From each sample, 150 μL of the top ethyl acetate phase was aspirated into a clean GC vial containing a glass insert adapter. Samples were run on an Agilent 5977E Series GC/MSD System. The samples were separated using an Agilent J&W DB- 1MS UI (Durabond Dimethylsiloxane) narrow bore column; 0.25 mm internal diameter; 60-meter length; 0.25-micron film coating. The GC was run as follows: each sample was run through the column at flow rate of 1 mL/minute, initially the column was held 40 °C for 1 minute, and then increased to 320 °C with a ramp rate of 20 °C per minute, finally the column was held at 320 °C for 6 minutes. To quantify mevalonate concentration, an external standard curve was generated by creating a calibration curve of mevalonate dissolved in ethyl acetate at concentrations of 5 mg/L, 10 mg/L, 25 mg/L, 50 mg/L, 100 mg/L, 250 mg/L, 500 mg/L, 1000 mg/L, 1500 mg/L, and 2000 mg/L of ethyl acetate.

## Acknowledgements

This work was financially supported by the National Science Foundation [grant numbers MCB- 1932780 to L.M.C., A.S., and R.B., and CHE-1852015 to R.S.] the National Institutes of Health [grant number R35GM133640 to B.K.K.], the Welch Foundation [grant number F-1929 to B.K.K.], and an NSF CAREER Award [1944334 to B.K.K.]. T.R.S. and G.P. were supported through National Science foundation Graduate Research Fellowships [program award number DGE-1610403]. A.S. was also financially supported through an Undergraduate Research Fellowship from the Office of Undergraduate Research at the University of Texas at Austin. We would like to thank Michaela A. Jones and Dr. Aditya M. Kunjapur from the University of Delaware for generously providing the pCG004 construct and valuable insight on methods to optimize transformation efficiency in *B. subtilis*. We would also like to thank the lab of Dr. Brian Pfleger from the University of Wisconsin-Madison for providing the pMP11 and pgRNA constructs and helpful protocols on CRISPR-Cas9-based recombineering in *E. coli.* We would also like to thank the members of the Contreras Lab for their helpful discussions during the development of this work. Lastly, we would like to thank the Mass Spectrometry Facility at the University of Texas at Austin for their assistance on developing a protocol to quantify mevalonate via GC-MS.

---

**Extended Figure 1.**
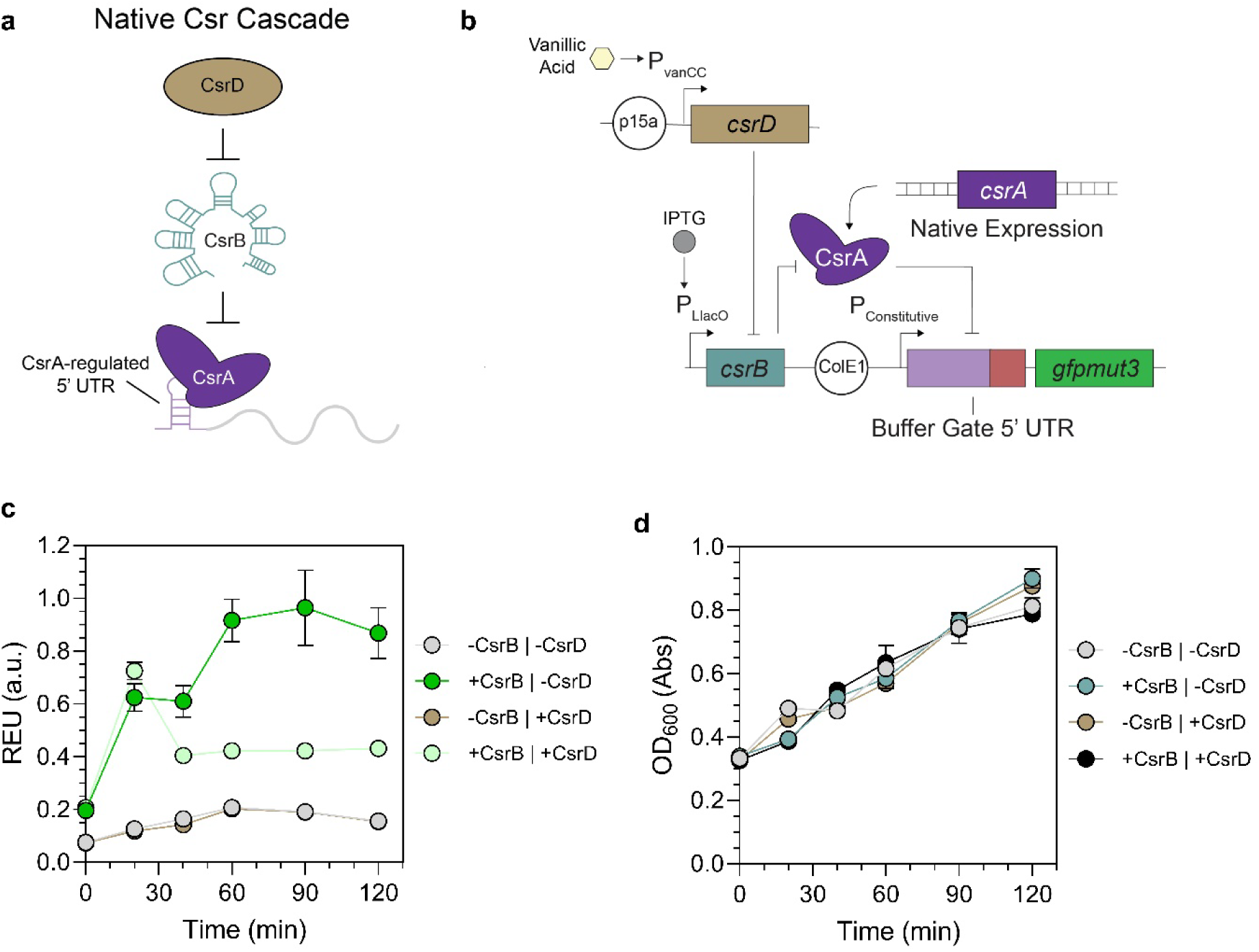
Engineering the Entire Csr Cascade to Achieve a Pulse Response. a) Native regulatory components of the full Csr Network. b) Diagram of the engineered Csr Cascade. The original Csr-regulated Buffer Gate construct is used in parallel with a plasmid containing vanillic acid-inducible CsrD. c) Pulse response time by inducing the *csrB* sRNA and CsrD protein simultaneously (light green line) compared to only inducing the *csrB* sRNA of the Csr-regulated Buffer Gate (dark green line). d) Growth curves for each induction to assess cellular health effects from managing the synthetic pulse system.

## References

1. Glick B. R. (1995). Metabolic load and heterologous gene expression. Biotechnology advances, 13(2), 247–261. 10.1016/0734-9750(95)00004-a

2. Kurland, C. G., & Dong, H. (1996). Bacterial growth inhibition by overproduction of protein. Molecular microbiology, 21(1), 1–4. 10.1046/j.1365-2958.1996.5901313.x

3. Liu, D., Mannan, A. A., Han, Y., Oyarzún, D. A., & Zhang, F. (2018). Dynamic metabolic control: towards precision engineering of metabolism. Journal of industrial microbiology & biotechnology, 45(7), 535–543. 10.1007/s10295-018-2013-9

4. Farmer, W. R., & Liao, J. C. (2000). Improving lycopene production in Escherichia coli by engineering metabolic control. Nature biotechnology, 18(5), 533–537. 10.1038/75398

5. Zhang, F., Carothers, J. M., & Keasling, J. D. (2012). Design of a dynamic sensor-regulator system for production of chemicals and fuels derived from fatty acids. Nature biotechnology, 30(4), 354–359. 10.1038/nbt.2149

6. Dahl, R. H., Zhang, F., Alonso-Gutierrez, J., Baidoo, E., Batth, T. S., Redding-Johanson, A. M., Petzold, C. J., Mukhopadhyay, A., Lee, T. S., Adams, P. D., & Keasling, J. D. (2013). Engineering dynamic pathway regulation using stress-response promoters. Nature biotechnology, 31(11), 1039–1046. 10.1038/nbt.2689

7. Stevens, J. T., & Carothers, J. M. (2015). Designing RNA-based genetic control systems for efficient production from engineered metabolic pathways. ACS synthetic biology, 4(2), 107–115. 10.1021/sb400201u

8. Dinh, C. V., & Prather, K. L. J. (2019). Development of an autonomous and bifunctional quorum-sensing circuit for metabolic flux control in engineered *Escherichia coli*. Proceedings of the National Academy of Sciences of the United States of America, 116(51), 25562–25568. 10.1073/pnas.1911144116

9. Gao, C., Hou, J., Xu, P., Guo, L., Chen, X., Hu, G., Ye, C., Edwards, H., Chen, J., Chen, W., & Liu, L. (2019). Programmable biomolecular switches for rewiring flux in Escherichia coli. Nature communications, 10(1), 3751. 10.1038/s41467-019-11793-7

10. Zhou, S., Yuan, S. F., Nair, P. H., Alper, H. S., Deng, Y., & Zhou, J. (2021). Development of a growth coupled and multi-layered dynamic regulation network balancing malonyl-CoA node to enhance (2S)-naringenin biosynthesis in Escherichia coli. Metabolic engineering, 67, 41–52. 10.1016/j.ymben.2021.05.007

11. Engineering Biology Research Consortium (2019). Engineering Biology: A Research Roadmap for the Next-Generation Bioeconomy. Retrieved from https://roadmap.ebrc.org. DOI: 10.25498/E4159B.

12. Frumkin, I., Schirman, D., Rotman, A., Li, F., Zahavi, L., Mordret, E., Asraf, O., Wu, S., Levy, S. F., & Pilpel, Y. (2017). Gene Architectures that Minimize Cost of Gene Expression. Molecular cell, 65(1), 142–153. 10.1016/j.molcel.2016.11.007

13. Westbrook, A., Tang, X., Marshall, R., Maxwell, C. S., Chappell, J., Agrawal, D. K., Dunlop, M. J., Noireaux, V., Beisel, C. L., Lucks, J., & Franco, E. (2019). Distinct timescales of RNA regulators enable the construction of a genetic pulse generator. Biotechnology and bioengineering, 116(5), 1139–1151. 10.1002/bit.26918

14. Takahashi, M. K., Chappell, J., Hayes, C. A., Sun, Z. Z., Kim, J., Singhal, V., Spring, K. J., Al-Khabouri, S., Fall, C. P., Noireaux, V., Murray, R. M., & Lucks, J. B. (2015). Rapidly characterizing the fast dynamics of RNA genetic circuitry with cell-free transcription-translation (TX-TL) systems. ACS synthetic biology, 4(5), 503–515. 10.1021/sb400206c

15. Ceroni, F., Boo, A., Furini, S., Gorochowski, T. E., Borkowski, O., Ladak, Y. N., Awan, A. R., Gilbert, C., Stan, G. B., & Ellis, T. (2018). Burden-driven feedback control of gene expression. Nature methods, 15(5), 387–393. 10.1038/nmeth.4635

16. Isaacs, F. J., Dwyer, D. J., Ding, C., Pervouchine, D. D., Cantor, C. R., & Collins, J. J. (2004). Engineered riboregulators enable post-transcriptional control of gene expression. Nature biotechnology, 22(7), 841–847. 10.1038/nbt986

17. Green, A. A., Silver, P. A., Collins, J. J., & Yin, P. (2014). Toehold switches: de-novo-designed regulators of gene expression. Cell, 159(4), 925–939. 10.1016/j.cell.2014.10.002

18. Pardee, K., Green, A. A., Takahashi, M. K., Braff, D., Lambert, G., Lee, J. W., Ferrante, T., Ma, D., Donghia, N., Fan, M., Daringer, N. M., Bosch, I., Dudley, D. M., O’Connor, D. H., Gehrke, L., & Collins, J. J. (2016). Rapid, Low-Cost Detection of Zika Virus Using Programmable Biomolecular Components. Cell, 165(5), 1255–1266. 10.1016/j.cell.2016.04.059

19. Kim, J., Zhou, Y., Carlson, P. D., Teichmann, M., Chaudhary, S., Simmel, F. C., Silver, P. A., Collins, J. J., Lucks, J. B., Yin, P., & Green, A. A. (2019). De novo-designed translation-repressing riboregulators for multi-input cellular logic. Nature chemical biology, 15(12), 1173–1182. 10.1038/s41589-019-0388-1

20. Lahiry, A., Stimple, S. D., Wood, D. W., & Lease, R. A. (2017). Retargeting a Dual-Acting sRNA for Multiple mRNA Transcript Regulation. ACS synthetic biology, 6(4), 648– 658. 10.1021/acssynbio.6b00261

21. Takahashi, M. K., & Lucks, J. B. (2013). A modular strategy for engineering orthogonal chimeric RNA transcription regulators. Nucleic acids research, 41(15), 7577–7588. 10.1093/nar/gkt452

22. Rodrigo, G., Landrain, T. E., & Jaramillo, A. (2012). De novo automated design of small RNA circuits for engineering synthetic riboregulation in living cells. Proceedings of the National Academy of Sciences of the United States of America, 109(38), 15271–15276. 10.1073/pnas.1203831109

23. Sowa, S. W., Vazquez-Anderson, J., Clark, C. A., De La Peña, R., Dunn, K., Fung, E. K., Khoury, M. J., & Contreras, L. M. (2015). Exploiting post-transcriptional regulation to probe RNA structures in vivo via fluorescence. Nucleic acids research, 43(2), e13. 10.1093/nar/gku1191

24. Mihailovic, M. K., Vazquez-Anderson, J., Li, Y., Fry, V., Vimalathas, P., Herrera, D., Lease, R. A., Powell, W. B., & Contreras, L. M. (2018). High-throughput in vivo mapping of RNA accessible interfaces to identify functional sRNA binding sites. Nature communications, 9(1), 4084. 10.1038/s41467-018-06207-z

25. Smolke, C. D., Carrier, T. A., & Keasling, J. D. (2000). Coordinated, differential expression of two genes through directed mRNA cleavage and stabilization by secondary structures. Applied and environmental microbiology, 66(12), 5399–5405. 10.1128/AEM.66.12.5399-5405.2000

26. Pfleger, B. F., Pitera, D. J., Smolke, C. D., & Keasling, J. D. (2006). Combinatorial engineering of intergenic regions in operons tunes expression of multiple genes. Nature biotechnology, 24(8), 1027–1032. 10.1038/nbt1226

27. Na, D., Yoo, S. M., Chung, H., Park, H., Park, J. H., & Lee, S. Y. (2013). Metabolic engineering of Escherichia coli using synthetic small regulatory RNAs. Nature biotechnology, 31(2), 170–174. 10.1038/nbt.2461

28. Saito, H., Kobayashi, T., Hara, T., Fujita, Y., Hayashi, K., Furushima, R., & Inoue, T. (2010). Synthetic translational regulation by an L7Ae-kink-turn RNP switch. Nature chemical biology, 6(1), 71–78. 10.1038/nchembio.273

29. Kennedy, A. B., Vowles, J. V., d’Espaux, L., & Smolke, C. D. (2014). Protein-responsive ribozyme switches in eukaryotic cells. Nucleic acids research, 42(19), 12306–12321. 10.1093/nar/gku875

30. Abudayyeh, O. O., Gootenberg, J. S., Essletzbichler, P., Han, S., Joung, J., Belanto, J. J., Verdine, V., Cox, D. B. T., Kellner, M. J., Regev, A., Lander, E. S., Voytas, D. F., Ting, A. Y., & Zhang, F. (2017). RNA targeting with CRISPR-Cas13. Nature, 550(7675), 280–284. 10.1038/nature24049

31. Colognori, D., Trinidad, M., & Doudna, J. A. (2023). Precise transcript targeting by CRISPR-Csm complexes. Nature biotechnology, 41(9), 1256–1264. 10.1038/s41587-022-01649-9

32. Cho, J. S., Yang, D., Prabowo, C. P. S., Ghiffary, M. R., Han, T., Choi, K. R., Moon, C. W., Zhou, H., Ryu, J. Y., Kim, H. U., & Lee, S. Y. (2023). Targeted and high-throughput gene knockdown in diverse bacteria using synthetic sRNAs. Nature communications, 14(1), 2359. 10.1038/s41467-023-38119-y

33. Romeo, T., & Babitzke, P. (2018). Global Regulation by CsrA and Its RNA Antagonists. Microbiology spectrum, 6(2), 10.1128/microbiolspec.RWR-0009-2017. https://doi.org/10.1128/microbiolspec.RWR-0009-2017

34. Morin, M., Ropers, D., Letisse, F., Laguerre, S., Portais, J. C., Cocaign-Bousquet, M., & Enjalbert, B. (2016). The post-transcriptional regulatory system CSR controls the balance of metabolic pools in upper glycolysis of Escherichia coli. Molecular microbiology, 100(4), 686–700. 10.1111/mmi.13343

35. Vakulskas, C. A., Potts, A. H., Babitzke, P., Ahmer, B. M., & Romeo, T. (2015). Regulation of bacterial virulence by Csr (Rsm) systems. Microbiology and molecular biology reviews : MMBR, 79(2), 193–224. 10.1128/MMBR.00052-14

36. Sowa, S. W., Gelderman, G., Leistra, A. N., Buvanendiran, A., Lipp, S., Pitaktong, A., Vakulskas, C. A., Romeo, T., Baldea, M., & Contreras, L. M. (2017). Integrative FourD omics approach profiles the target network of the carbon storage regulatory system. Nucleic acids research, 45(4), 1673–1686. 10.1093/nar/gkx048

37. Leistra, A. N., Gelderman, G., Sowa, S. W., Moon-Walker, A., Salis, H. M., & Contreras, L. M. (2018). A Canonical Biophysical Model of the CsrA Global Regulator Suggests Flexible Regulator-Target Interactions. Scientific reports, 8(1), 9892. 10.1038/s41598-018-27474-2

38. Potts, A. H., Vakulskas, C. A., Pannuri, A., Yakhnin, H., Babitzke, P., & Romeo, T. (2017). Global role of the bacterial post-transcriptional regulator CsrA revealed by integrated transcriptomics. Nature communications, 8(1), 1596. 10.1038/s41467-017-01613-1

39. Rojano-Nisimura, A. M., Simmons, T. R., Leistra, A. N., Mihailovic, M. K., Buchser, R., Ekdahl, A. M., Joseph, I., Curtis, N. C., & Contreras, L. M. (2023). CsrA Shows Selective Regulation of sRNA-mRNA Networks. bioRxiv : the preprint server for biology, 2023.03.29.534774. 10.1101/2023.03.29.534774

40. Dubey, A. K., Baker, C. S., Romeo, T., & Babitzke, P. (2005). RNA sequence and secondary structure participate in high-affinity CsrA-RNA interaction. *RNA (New York*, N.Y.), 11(10), 1579–1587. 10.1261/rna.2990205

41. Baker, C. S., Morozov, I., Suzuki, K., Romeo, T., & Babitzke, P. (2002). CsrA regulates glycogen biosynthesis by preventing translation of glgC in Escherichia coli. Molecular microbiology, 44(6), 1599–1610. 10.1046/j.1365-2958.2002.02982.x

42. Renda, A., Poly, S., Lai, Y. J., Pannuri, A., Yakhnin, H., Potts, A. H., Bevilacqua, P. C., Romeo, T., & Babitzke, P. (2020). CsrA-Mediated Translational Activation of *ymdA* Expression in Escherichia coli. mBio, 11(5), e00849–20. 10.1128/mBio.00849-20

43. Yakhnin, A. V., Baker, C. S., Vakulskas, C. A., Yakhnin, H., Berezin, I., Romeo, T., & Babitzke, P. (2013). CsrA activates flhDC expression by protecting flhDC mRNA from RNase E-mediated cleavage. Molecular microbiology, 87(4), 851–866. 10.1111/mmi.12136

44. Weilbacher, T., Suzuki, K., Dubey, A. K., Wang, X., Gudapaty, S., Morozov, I., Baker, C. S., Georgellis, D., Babitzke, P., & Romeo, T. (2003). A novel sRNA component of the carbon storage regulatory system of Escherichia coli. Molecular microbiology, 48(3), 657–670. 10.1046/j.1365-2958.2003.03459.x

45. Leng, Y., Vakulskas, C. A., Zere, T. R., Pickering, B. S., Watnick, P. I., Babitzke, P., & Romeo, T. (2016). Regulation of CsrB/C sRNA decay by EIIA(Glc) of the phosphoenolpyruvate: carbohydrate phosphotransferase system. Molecular microbiology, 99(4), 627–639. 10.1111/mmi.13259

46. Adamson, D. N., & Lim, H. N. (2013). Rapid and robust signaling in the CsrA cascade via RNA-protein interactions and feedback regulation. Proceedings of the National Academy of Sciences of the United States of America, 110(32), 13120–13125. 10.1073/pnas.1308476110

47. Liu, M. Y., Yang, H., & Romeo, T. (1995). The product of the pleiotropic Escherichia coli gene csrA modulates glycogen biosynthesis via effects on mRNA stability. Journal of bacteriology, 177(10), 2663–2672. 10.1128/jb.177.10.2663-2672.1995

48. Moon, T. S., Lou, C., Tamsir, A., Stanton, B. C., & Voigt, C. A. (2012). Genetic programs constructed from layered logic gates in single cells. Nature, 491(7423), 249–253. 10.1038/nature11516

49. Romeo T. (1998). Global regulation by the small RNA-binding protein CsrA and the non-coding RNA molecule CsrB. Molecular microbiology, 29(6), 1321–1330. 10.1046/j.1365-2958.1998.01021.x

50. Leistra, A. N., Amador, P., Buvanendiran, A., Moon-Walker, A., & Contreras, L. M. (2017). Rational Modular RNA Engineering Based on In Vivo Profiling of Structural Accessibility. ACS synthetic biology, 6(12), 2228–2240. 10.1021/acssynbio.7b00185

51. Salis H. M. (2011). The ribosome binding site calculator. Methods in enzymology, 498, 19–42. 10.1016/B978-0-12-385120-8.00002-4

52. Meyer, A. J., Segall-Shapiro, T. H., Glassey, E., Zhang, J., & Voigt, C. A. (2019). Escherichia coli “Marionette” strains with 12 highly optimized small-molecule sensors. Nature chemical biology, 15(2), 196–204. 10.1038/s41589-018-0168-3

53. Partipilo, G., Graham, A. J., Belardi, B., & Keitz, B. K. (2022). Extracellular Electron Transfer Enables Cellular Control of Cu(I)-Catalyzed Alkyne-Azide Cycloaddition. ACS central science, 8(2), 246–257. 10.1021/acscentsci.1c01208

54. Gao, Y., Zhou, Y., Ji, X., Graham, A. J., Dundas, C. M., Mahfoud, I. E. M., Tibbett, B. M., Tan, B., Partipilo, G., Dodabalapur, A., Rivnay, J., & Keitz, B. K. (2023). A Hybrid Transistor with Transcriptionally Controlled Computation and Plasticity. bioRxiv : the preprint server for biology, 2023.08.16.553547. 10.1101/2023.08.16.553547

55. Zhao, F., Chavez, M. S., Naughton, K. L., Niman, C. M., Atkinson, J. T., Gralnick, J. A., El-Naggar, M. Y., & Boedicker, J. Q. (2022). Light-Induced Patterning of Electroactive Bacterial Biofilms. ACS synthetic biology, 11(7), 2327–2338. 10.1021/acssynbio.2c00024

56. Binnenkade, L., Lassak, J., & Thormann, K. M. (2011). Analysis of the BarA/UvrY two-component system in Shewanella oneidensis MR-1. PloS one, 6(9), e23440. 10.1371/journal.pone.0023440

57. Anderson, J. C., Dueber, J. E., Leguia, M., Wu, G. C., Goler, J. A., Arkin, A. P., & Keasling, J. D. (2010). BglBricks: A flexible standard for biological part assembly. Journal of biological engineering, 4(1), 1. 10.1186/1754-1611-4-1

58. Altegoer, F., Rensing, S. A., & Bange, G. (2016). Structural basis for the CsrA-dependent modulation of translation initiation by an ancient regulatory protein. Proceedings of the National Academy of Sciences of the United States of America, 113(36), 10168–10173. 10.1073/pnas.1602425113

59. Praveschotinunt, P., Duraj-Thatte, A. M., Gelfat, I., Bahl, F., Chou, D. B., & Joshi, N. S. (2019). Engineered E. coli Nissle 1917 for the delivery of matrix-tethered therapeutic domains to the gut. Nature communications, 10(1), 5580. 10.1038/s41467-019-13336-6

60. Rottinghaus, A. G., Ferreiro, A., Fishbein, S. R. S., Dantas, G., & Moon, T. S. (2022). Genetically stable CRISPR-based kill switches for engineered microbes. Nature communications, 13(1), 672. 10.1038/s41467-022-28163-5

61. Stork, D. A., Squyres, G. R., Kuru, E., Gromek, K. A., Rittichier, J., Jog, A., Burton, B. M., Church, G. M., Garner, E. C., & Kunjapur, A. M. (2021). Designing efficient genetic code expansion in Bacillus subtilis to gain biological insights. Nature communications, 12(1), 5429. 10.1038/s41467-021-25691-4

62. Mukherjee, S., Yakhnin, H., Kysela, D., Sokoloski, J., Babitzke, P., & Kearns, D. B. (2011). CsrA-FliW interaction governs flagellin homeostasis and a checkpoint on flagellar morphogenesis in Bacillus subtilis. Molecular microbiology, 82(2), 447–461. 10.1111/j.1365-2958.2011.07822.x

63. Sun, Z., Zhou, N., Zhang, W., Xu, Y., & Yao, Y. F. (2022). Dual role of CsrA in regulating the hemolytic activity of *Escherichia coli* O157:H7. Virulence, 13(1), 859–874. 10.1080/21505594.2022.2073023

64. Pourciau, C., Lai, Y. J., Gorelik, M., Babitzke, P., & Romeo, T. (2020). Diverse Mechanisms and Circuitry for Global Regulation by the RNA-Binding Protein CsrA. Frontiers in microbiology, 11, 601352. 10.3389/fmicb.2020.601352

65. McKee, A. E., Rutherford, B. J., Chivian, D. C., Baidoo, E. K., Juminaga, D., Kuo, D., Benke, P. I., Dietrich, J. A., Ma, S. M., Arkin, A. P., Petzold, C. J., Adams, P. D., Keasling, J. D., & Chhabra, S. R. (2012). Manipulation of the carbon storage regulator system for metabolite remodeling and biofuel production in Escherichia coli. Microbial cell factories, 11, 79. 10.1186/1475-2859-11-79

66. Martin, V. J., Pitera, D. J., Withers, S. T., Newman, J. D., & Keasling, J. D. (2003). Engineering a mevalonate pathway in Escherichia coli for production of terpenoids. Nature biotechnology, 21(7), 796–802. 10.1038/nbt833

67. Pitera, D. J., Paddon, C. J., Newman, J. D., & Keasling, J. D. (2007). Balancing a heterologous mevalonate pathway for improved isoprenoid production in Escherichia coli. Metabolic engineering, 9(2), 193–207. 10.1016/j.ymben.2006.11.002

68. Shin, J., South, E. J., & Dunlop, M. J. (2022). Transcriptional Tuning of Mevalonate Pathway Enzymes to Identify the Impact on Limonene Production in *Escherichia coli*. ACS omega, 7(22), 18331–18338. 10.1021/acsomega.2c00483

69. Leavell, M. D., McPhee, D. J., & Paddon, C. J. (2016). Developing fermentative terpenoid production for commercial usage. Current opinion in biotechnology, 37, 114–119. 10.1016/j.copbio.2015.10.007

70. Kant, G., Pandey, A., Shekhar, H., & Srivastava, S. (2023). Enhanced bio-synthesis of isoprene via modifying mevalonate and methylerythritol phosphate pathways for industrial application: A review. Process Biochemistry, 130, 256–271. 10.1016/j.procbio.2023.04.021

71. Ma, S. M., Garcia, D. E., Redding-Johanson, A. M., Friedland, G. D., Chan, R., Batth, T. S., Haliburton, J. R., Chivian, D., Keasling, J. D., Petzold, C. J., Lee, T. S., & Chhabra, S. R. (2011). Optimization of a heterologous mevalonate pathway through the use of variant HMG-CoA reductases. Metabolic engineering, 13(5), 588–597. 10.1016/j.ymben.2011.07.001

72. Wang, J., Niyompanich, S., Tai, Y. S., Wang, J., Bai, W., Mahida, P., Gao, T., & Zhang, K. (2016). Engineering of a Highly Efficient Escherichia coli Strain for Mevalonate Fermentation through Chromosomal Integration. Applied and environmental microbiology, 82(24), 7176–7184. 10.1128/AEM.02178-16

73. Xu, X., Xie, M., Zhao, Q., Xian, M., & Liu, H. (2018). Microbial production of mevalonate by recombinant Escherichia coli using acetic acid as a carbon source. Bioengineered, 9(1), 116–123. 10.1080/21655979.2017.1323592

74. Calos M. P. (1978). DNA sequence for a low-level promoter of the lac repressor gene and an ’up’ promoter mutation. Nature, 274(5673), 762–765. 10.1038/274762a0

75. Cetnar, D. P., & Salis, H. M. (2021). Systematic Quantification of Sequence and Structural Determinants Controlling mRNA stability in Bacterial Operons. ACS synthetic biology, 10(2), 318–332. 10.1021/acssynbio.0c00471

76. Wang, B., Kitney, R. I., Joly, N., & Buck, M. (2011). Engineering modular and orthogonal genetic logic gates for robust digital-like synthetic biology. Nature communications, 2, 508. 10.1038/ncomms1516

77. Groseclose, T. M., Rondon, R. E., Herde, Z. D., Aldrete, C. A., & Wilson, C. J. (2020). Engineered systems of inducible anti-repressors for the next generation of biological programming. Nature communications, 11(1), 4440. 10.1038/s41467-020-18302-1

78. Nielsen, A. A., Der, B. S., Shin, J., Vaidyanathan, P., Paralanov, V., Strychalski, E. A., Ross, D., Densmore, D., & Voigt, C. A. (2016). Genetic circuit design automation. Science (New York, N.Y.), 352(6281), aac7341. 10.1126/science.aac7341

79. Huang, B. D., Groseclose, T. M., & Wilson, C. J. (2022). Transcriptional programming in a Bacteroides consortium. Nature communications, 13(1), 3901. 10.1038/s41467-022-31614-8

80. Cardinale, S., & Arkin, A. P. (2012). Contextualizing context for synthetic biology-- identifying causes of failure of synthetic biological systems. Biotechnology journal, 7(7), 856–866. 10.1002/biot.201200085

81. Kittleson, J. T., Wu, G. C., & Anderson, J. C. (2012). Successes and failures in modular genetic engineering. Current opinion in chemical biology, 16(3-4), 329–336. 10.1016/j.cbpa.2012.06.009

82. Castillo-Hair, S. M., Fujita, M., Igoshin, O. A., & Tabor, J. J. (2019). An Engineered *B. subtilis* Inducible Promoter System with over 10 000-Fold Dynamic Range. ACS synthetic biology, 8(7), 1673–1678. 10.1021/acssynbio.8b00469

83. Kushwaha, M., & Salis, H. M. (2015). A portable expression resource for engineering cross-species genetic circuits and pathways. Nature communications, 6, 7832. 10.1038/ncomms8832

84. Gordon, G. C., Korosh, T. C., Cameron, J. C., Markley, A. L., Begemann, M. B., & Pfleger, B. F. (2016). CRISPR interference as a titratable, trans-acting regulatory tool for metabolic engineering in the cyanobacterium Synechococcus sp. strain PCC 7002. Metabolic engineering, 38, 170–179. 10.1016/j.ymben.2016.07.007

85. Ho, H. I., Fang, J. R., Cheung, J., & Wang, H. H. (2020). Programmable CRISPR-Cas transcriptional activation in bacteria. Molecular systems biology, 16(7), e9427. 10.15252/msb.20199427

86. Zhang, S., & Voigt, C. A. (2018). Engineered dCas9 with reduced toxicity in bacteria: implications for genetic circuit design. Nucleic acids research, 46(20), 11115–11125. 10.1093/nar/gky884

87. Shi, L., & Tu, B. P. (2015). Acetyl-CoA and the regulation of metabolism: mechanisms and consequences. Current opinion in cell biology, 33, 125–131. 10.1016/j.ceb.2015.02.003

88. McGuffie, M. J., & Barrick, J. E. (2021). pLannotate: engineered plasmid annotation. Nucleic acids research, 49(W1), W516–W522. 10.1093/nar/gkab374

