## Supplementary Information for "Rewiring native post-transcriptional carbon regulators to build multi-layered genetic circuits and optimize engineered microbes for bioproduction"

### Table of Contents

**Supplementary Materials and Methods**

**Supplementary Tables**
Table S1 - Ingredients in *Shewanella* Basal Medium
Table S2 - Ingredients in M9Y Medium
Table S3 - Ingredients in 10x MC
Table S4 - Ingredients in MC Medium
Table S5 - Bacterial Strains and Plasmids used in this study.
Table S6 - Oligonucleotides used in this study.
Table S7 - 5' UTRs and CsrB sRNA sequences used in this study.

**Supplementary Figures**

Supplementary Figure S1 - Predicted secondary structures of the wild type and engineered *glgC* 5' UTR sequences to evaluate optimal spacer length.

Supplementary Figure S2 - Growth Curve of induced and uninduced Csr-regulated Buffer Gate containing cultures.

Supplementary Figure S3 - Confirming *in vivo* CsrA repression of the engineered *glgC* 5' UTR-GFP fusion construct.

Supplementary Figure S4 - Evaluating Csr-Controlled Buffer Gate Performance using mutant CsrB sRNA sequences.

Supplementary Figure S5 - Evaluating Csr-Controlled Buffer Gate Performance using variable 5-nucleotide spacers in the engineered *glgC* 5' UTR.

Supplementary Figure S6 - Csr-Controlled Buffer Gate Performance of all additional CsrB sRNA and 5-nt spacer combinations.

Supplementary Figure S7 - REUs of all Csr Buffer Gate combinations of *csrB* sRNAs and 5-nt RBS sequences.

Supplementary Figure S8 - Performance of the Csr-Controlled Buffer Gate in various *csr* deletion strains.

Supplementary Figure S9 - Testing Csr-Controlled Buffer Gate Performance using Anderson Promoter Library sequences.

Supplementary Figure S10 - Performance of the Csr-Controlled Buffer Gate against a LacI-controlled transcriptional buffer gate in *S. oneidensis*.

Supplementary Figure S11 - Comparing the Csr-Controlled Buffer Gate against a Lac-regulated transcriptional Buffer Gate across multiple Csr-containing bacteria.

Supplementary Figure S12 - Growth Curves of Mevalonate Producing Strains.

Supplementary Figure S13 - Mevalonate production from placI-Mev-containing cultures at different induction conditions.

Supplementary Figure S14 - qPCR on synthetic mevalonate operon mRNA in the Csr-regulated and LacI-regulated systems.

Supplementary Figure S15 - Mevalonate production in the *ΔcsrBCD* and *csrA::kan* strains for the Csr-regulated and LacI-regulated systems.

### Supplementary Materials and Methods

#### Materials:

Sodium DL-lactate ($NaC_3H_5O_3$, TCI, 60% in water), HEPES buffer solution ($C_8H_{18}N_2O_4S$, VWR, 1 M in water, pH = 7.3), potassium phosphate dibasic ($K_2HPO_4$, Sigma-Aldrich), isopropyl ß-D-1-thiogalactopyranoside (IPTG, Teknova), potassium phosphate monobasic ($KH_2PO_4$, Sigma-Aldrich), sodium chloride (NaCl, VWR), kanamycin sulfate ($C_{18}H_{38}N_4O_{15}S$, Growcells) ammonium sulfate ($(NH_4)_2SO_4$, Fisher Scientific), magnesium(II) sulfate heptahydrate ($MgSO_4 \cdot 7H_2O$, VWR), trace mineral supplement (ATCC), casamino acids (VWR), sodium phosphate dibasic (MilliPore Sigma), sodium phosphate monobasic (MilliPore Sigma), glucose (Fisher)

#### Methods:

**Transformations *S. oneidensis*:** Overnight culture was grown of the parent strain (5 mL) in LB. The culture was centrifuged 4200 rpm for 8 minutes and decanted. The cell pellet was resuspended in 1 mL of 10% glycerol pre-warmed at 37 °C. The cells were transferred to a 1.7-mL Eppendorf tube and centrifuged at 7900 xg for 2 minutes and decanted. This was repeated with warm 10% glycerol 2 more times. The final resuspension was in 250 µL. The purified plasmid (10-100 ng of DNA, approx. 1 µL) was mixed with 33 µL of cell suspension and allowed to sit for 15 minutes. The cells were then electroporated at 1250 V and quickly resuspended in 200 µL of warm LB. The cells were allowed to recover to 2 hours at 30 °C shaking at 250 rpm before being diluted 100-fold and plated onto antibiotic selection plates.

**Transformations *E. coli* Nissle 1917:** Overnight culture was grown of the parent strain (5 mL) in LB. The culture was centrifuged 4200 rpm for 8 minutes and decanted. The cell pellet was resuspended in 1 mL of 10% glycerol cooled to 4 °C. The cells were transferred to a 1.7-mL Eppendorf tube and centrifuged at 6000 xg for 3 minutes and decanted. This was repeated with cold 10% glycerol 2 more times. The final resuspension was in 250 µL. The purified plasmid (10-100 ng of DNA, approx. 1 µL) was mixed with 33 µL of cell suspension and allowed to sit for 15 minutes. The cells were then electroporated at 2500 V and quickly resuspended in 200 µL of warm LB. The cells were allowed to recover to 2 hours at 30 °C shaking at 250 rpm before being diluted 100-fold and plated onto antibiotic selection plates.

**Transformations *P. putida*:** Overnight culture was grown of the parent strain (5 mL) in LB. The culture was centrifuged 4200 rpm for 10 minutes and decanted. The cell pellet was resuspended in 1 mL of 10% glycerol cooled to 4 °C. The cells were transferred to a 1.7-mL Eppendorf tube and centrifuged at 7900 xg for 2 minutes and decanted. This was repeated with cold 10% glycerol 2 more times. The final resuspension was in 800 µL. The purified plasmid (40-400 ng of DNA, approx. 4 µL) was mixed with 40 µL of cell suspension and allowed to sit for 15 minutes. The cells were then electroporated at 2400 V and quickly resuspended in 200 µL of warm LB. The cells were allowed to recover for 2 hours at 30 °C shaking at 250 rpm before being diluted 100-fold and plated onto antibiotic selection plates.

**Transformation *B. subtilis*:** Freshly streaked colonies of *B. subtilis* were inoculated into 1 mL of 1x MC +3 mM $MgSO_4$ (ingredients in Tables S3 and S4) in a disposable culture tube. Inoculated samples were thoroughly pipetted to break up any remaining biofilm. Samples were then grown at 37 °C for two hours to become competent cells. Once the competent cell cultures reached an $OD_{600}$ of 0.5, approximately two hours, 200 µL of competent cell culture was added to a fresh tube containing 500 ng of purified plasmid. The mixture of competent cells and plasmid was incubated for two hours at 37 °C. Following the two-hour incubation, the entire 200 µL mixture was plated onto an LB + 5 µg/mL Chloramphenicol and grown at 37 °C overnight. Single colonies were picked the next day and grown in 1 mL of LB in at 15 mL culture tube for four hours to be used in plasmid sequence verification via Miniprep (Qiagen) or saved for glycerol stocks.

**Large scale fluorescent assay for *Shewanella oneidensis*:** Overnight cultures were grown in triplicate in 5 mL of LB supplemented with the appropriate antibiotic for maintaining the plasmid. *S. oneidensis* was grown at 30 °C. The overnight cell cultures were diluted 1/100 into 30 mL of LB supplemented with 25 µg/mL kanamycin. The cells were grown, shaking at 200 rpm, in 250 mL flasks at 30 °C and induced. Each time point was taken by diluting the culture 1/4 in PBS and measuring via platereader to determine $OD_{600}$ and the fluorescence (excitation 488 nm, emission 515 nm, gain 1800). Fold turn-on was determined by normalizing the fluorescent/$OD_{600}$ readings by the value of the uninduced sample.

**Small-scale fluorescent assay:** Overnight cultures were grown in triplicate in 5 mL of LB supplemented with the appropriate antibiotic for maintaining the plasmid. *S. oneidensis* was grown at 30 °C. The overnight cell cultures were diluted 20 µL into 175 µL of SBM containing lactate and kanamycin to yield a solution with 20 mM lactate and 20 mg/mL kanamycin in a clear bottom, black walled Greiner 96-well plate. The solution was allowed to grow aerobically at 30 °C and shaking for approximately 2 hours until the average $OD_{600}$ is 0.2. Then 5 µL of IPTG of varying stock concentrations was diluted in to induce the cells. The fluorescence was tracked on a plate reader for 20 hours (excitation 488 nm, emission 515 nm, gain 1200). The same assay was used for *E. coli* Nissle 1917 and *P. Putida* with the following amendments. The cells were grown at 37 °C, the reaction media was M9Y, glucose (20 mM) was used as the carbon source, and varying antibiotics were used for the different strains. The gain for *E. coli* Nissle 1917 was changed to 1800. At 20 hours, the fluorescent readings were normalized by the $OD_{600}$ of the well. Fold turn-on was determined by normalizing the fluorescent/$OD_{600}$ readings by the value of the uninduced sample.

**RNA Extraction:** 5 mL of samples from desired cultures were centrifuged at 4000xg for 7 minutes at 4 °C. After being centrifuged, the supernatant was removed, and cells were flash frozen with liquid nitrogen and stored at -80 °C for future use. When ready for processing, cell pellets were thawed on ice. Next the pellets were resuspended in 200 µL of 2 mg/mL Lysozyme in 1x TBE (Invitrogen). The samples were incubated at room temperature for 10 minutes. Next, 700 µL of 1x DNA/RNA shield (Zymo) was added to each sample. Samples were then centrifuged at 21,000xg for 10 minutes at room temperature. The supernatant was aspirated and transferred into a clean 1.7 mL microcentrifuge tube, and one sample volume of DNA/RNA Lysis Buffer (Zymo) was added to each sample. RNA from each sample was purified by following the instructions provided in the Zymo *Quick*-DNA/RNA Miniprep Plus Kit. RNA was eluted in the final step by using 40 µL of UltraPure Water (Invitrogen). RNA Concentrations were determined using NanoDrop (ThermoFisher).

**RT-qPCR Protocol:** RNA Extraction samples were diluted down to concentrations of 1 ng/µL, 0.1 ng/µL, and 0.01 ng/µL using UltraPure Water (Invitrogen). Next, the RT-qPCR reactions were set-up according to the instructions provided in the Luna Universal One-Step RT-qPCR Kit.

**Table S1.** Ingredients in *Shewanella* Basal Medium. Growth media was supplemented with casamino acids and Wolfe's mineral solution. (1, 2)

| Ingredient | Quantity for 1 L of 1 X SBM |
|---|---|
| $K_2HPO_4$ | 225 mg |
| $KH_2PO_4$ | 225 mg |
| NaCl | 460 mg |
| $(NH_4)_2SO_4$ | 225 mg |
| $MgSO_4 \cdot 7H_2O$ | 117 mg |
| HEPES | 100 mL of 1 M HEPES |
| Casamino acids | 5 mL of 10% casamino acids |
| Wolfe's mineral solution | 5 mL of Wolfe's Mineral Solution |
| $ddH_2O$ | Up to 1 L, adjust to pH = 7.2 |

**Table S2.** Ingredients in M9Y Medium. Growth media was supplemented with casamino acids and Wolfe's mineral solution. (3)

| Ingredient | Quantity for 1 L of 1 X SBM |
|---|---|
| $Na_2HPO_4$ | 31 g |
| $NaH_2PO_4$ | 15 g |
| NaCl | 2.5 g |
| $MgSO_4$ | 240 mg |
| $CaCl_2$ | 10 mg |

**Table S3**. Ingredients in 10x MC. Ingredients should be added in order. This protocol is derived from (4).

| Ingredient | Quantity for 100 mL of 10x MC |
|---|---|
| $K_2HPO_4$ | 8.70 g |
| $KH_2PO_4$ | 8.42 g |
| Glucose | 20 g |
| Casein Hydrolysate | 1 g |
| Potassium Glutamate | 2 g |
| Ferric Ammonium Citrate | 22 g |
| Sodium Citrate | 7.74 g |

**Table S4.** Ingredients in MC Medium

| Ingredient | Quantity for 1 L of 1x MC |
|---|---|
| $ddH_2O$ | 997 mL |
| 10x MC | 100 mL |
| 1M $MgSO_4$ | 3 mL |

**Table S5.** Bacterial Strains and Plasmids used in this study.

| Strain or Plasmid | Description/Genotype | Reference |
|---|---|---|
| *E. coli DH5α* | *fhuA2 lac(del)U169 phoA glnV44 Φ80' lacZ(del)M15 gyrA96 recA1 relA1 endA1 thi-1 hsdR17* | Contreras Lab, U. of Texas at Austin |

| | | |
|---|---|---|
| *E. coli MG1655* | *F- lambda- ilvG- rfb-50 rph-1* | Contreras Lab, U. of Texas at Austin |
| CML379 | *MG1655 ΔcsrA::kan* | (5) |
| *ΔcsrBC* | *MG1655 ΔcsrBΔcsrC* | (5) |
| *ΔBCD* | *MG1655 ΔcsrBΔcsrCΔcsrD* | This study |
| *S. oneidensis* MR-1 | MR-1 (ATCC700550), wild-type strain | American-Type Culture Collection |
| *E. coli* Nissle 1917 | Mutaflor confirmed by 16S RNA sequencing | Mutaflor |
| *P. putida* | Wild-type strain (KD2440) | Andrew D. Ellington, U. of Texas at Austin |
| pTRS034 | Csr-Controlled GFP Buffer Gate: *glgC 5' UTR*-TTGGT-*gfpmut3* ($P_{con}$12), WT CsrB ($P_{LlacO}$; lacI), ColE1 ori, CarbR | This study |
| pTRS040 | Csr-Controlled GFP Buffer Gate: *glgC 5' UTR*-TATTA-*gfpmut3* ($P_{con}$12), WT CsrB ($P_{LlacO}$; lacI), ColE1 ori, CarbR | This study |
| pTRS041 | Csr-Controlled GFP Buffer Gate: *glgC 5' UTR*-TTCAT-*gfpmut3* ($P_{con}$12), WT CsrB ($P_{LlacO}$; lacI), ColE1 ori, CarbR | This study |
| pTRS042 | Transcriptional Buffer Gate: *gfpmut3* ($P_{LlacO}$; *lacI*), ColE1 ori, CarbR | This study |
| pTRS043 | Csr-Controlled EYFP Buffer Gate: *glgC 5' UTR*-TTGGT-*eyfp* ($P_{con}$12), WT CsrB ($P_{LlacO}$; lacI), ColE1 ori, CarbR | This study |
| pTRS044 | Csr-Controlled EYFP Buffer Gate: *glgC 5' UTR*-TTGGT-*mcherry* ($P_{con}$12), WT CsrB ($P_{LlacO}$; lacI), ColE1 ori, CarbR | This study |
| pTRS046 | Csr-Controlled GFP Buffer Gate: *glgC 5' UTR*-TTGGT-*gfpmut3* ($P_{con}$12), WT CsrB ($P_{LlacO}$; lacI), pBBR1 ori, CmR | This study |
| pTRS048 | Csr-Controlled GFP Buffer Gate: *glgC 5' UTR*-TTGGT-*gfpmut3* ($P_{con}$12), L2 CsrB ($P_{LlacO}$; lacI), ColE1 ori, CarbR | This study |
| pTRS049 | Csr-Controlled GFP Buffer Gate: *glgC 5' UTR*-TTGGT-*gfpmut3* ($P_{con}$12), L3 CsrB ($P_{LlacO}$; lacI), ColE1 ori, CarbR | This study |
| pTRS050 | Csr-Controlled GFP Buffer Gate: *glgC 5' UTR*-TTGGT-*gfpmut3* ($P_{con}$12), H4 CsrB ($P_{LlacO}$; lacI), ColE1 ori, CarbR | This study |
| pTRS051 | Csr-Controlled GFP Buffer Gate: *glgC 5' UTR*-TTGGT-*gfpmut3* ($P_{con}$12), H8 CsrB ($P_{LlacO}$; lacI), ColE1 ori, CarbR | This study |
| pTRS052 | Csr-Controlled GFP Buffer Gate: *glgC 5' UTR*-TTGGT-*gfpmut3* ($P_{con}$12), H11 CsrB ($P_{LlacO}$; lacI), ColE1 ori, CarbR | This study |
| pTRS053 | Csr-Controlled GFP Buffer Gate: *glgC 5' UTR* all CsrA binding sites mutated-TTGGT-*gfpmut3* ($P_{con}$12), WT CsrB ($P_{LlacO}$; lacI), ColE1 ori, CarbR | This study |
| pTRS059 | Constitutive *gfpmut3* ($P_{con}$12), ColE1 ori, CarbR | This study |
| pTRS060 | Csr-Controlled GFP Buffer Gate: *glgC 5' UTR*-TTGGT-*gfpmut3* ($P_{con}$12), WT CsrB ($P_{LlacO}$; lacI), ColE1 ori, KanR | This study |
| pTRS061 | Csr-Controlled GFP Buffer Gate: *glgC 5' UTR*- | This study |

| | | |
|---|---|---|
| | TTCAT-*gfpmut3* ($P_{con}$12), H11 CsrB ($P_{LlacO}$; lacI), ColE1 ori, CarbR | |
| pTRS062 | Csr-Controlled GFP Buffer Gate: glgC 5' UTR-TTCAT-gfpmut3 ($P_{con}$12), L2 CsrB ($P_{LlacO}$; lacI), ColE1 ori, CarbR | This study |
| pTRS063 | Csr-Controlled GFP Buffer Gate: *glgC 5' UTR*-TTGGT-*mcherry* ($P_{con}$12), WT CsrB ($P_{LlacO}$; lacI), p15a ori, KanR | This study |
| pTRS064 | Csr-Controlled GFP Buffer Gate: *glgC 5' UTR*-TTGGT-*eyfp* ($P_{con}$12), WT CsrB ($P_{LlacO}$; lacI), p15a ori, KanR | This study |
| pTRS065 | Transcriptional Buffer Gate: *gfpmut3* ($P_{LlacO}$; *lacI*), ColE1 ori, KanR | This study |
| pTRS066 | Csr-Controlled GFP Buffer Gate: *glgC 5' UTR*-TTACA-*gfpmut3* ($P_{con}$12), WT CsrB ($P_{LlacO}$; lacI), ColE1 ori, KanR | This study |
| pTRS067 | Csr-Controlled GFP Buffer Gate: *glgC 5' UTR*-TTACA-*gfpmut3* ($P_{con}$12), H11 CsrB ($P_{LlacO}$; lacI), ColE1 ori, CarbR | This study |
| pTRS068 | Csr-Controlled GFP Buffer Gate: *glgC 5' UTR*-TTGGT-*gfpmut3* ($P_{con}$12), ColE1 ori, CarbR | This study |
| pTRS069 | Csr-Controlled EYFP Buffer Gate: *glgC 5' UTR*-TTGGT-*eyfp* ($P_{con}$12), ColE1 ori, CarbR | This study |
| pTRS070 | Csr-Controlled mCherry Buffer Gate: *glgC 5' UTR*-TTGGT-*mcherry* ($P_{con}$12), ColE1 ori, CarbR | This study |
| pTRS071 | Csr-Controlled GFP Buffer Gate: *glgC 5' UTR*-TTTAC-*gfpmut3* ($P_{con}$12), WT CsrB ($P_{LlacO}$; lacI), ColE1 ori, CarbR | This study |
| pTRS072 | Csr-Controlled Operon Buffer Gate: *glgC 5' UTR*-TTGGT-*gfpmut3*- *glgC 5' UTR*-TTGGT-*mcherry*-*glgC 5' UTR*-TTGGT-*eyfp* ($P_{con}$12), WT CsrB ($P_{LlacO}$; lacI), ColE1 ori, CarbR | This study |
| pTRS073 | Csr-Controlled GFP Buffer Gate: *glgC 5' UTR*-TTTAC-*gfpmut3* ($P_{con}$12), L2 CsrB ($P_{LlacO}$; lacI), ColE1 ori, CarbR | This study |
| pTRS074 | Csr-Controlled GFP Buffer Gate: *glgC 5' UTR*-TTTAC-*gfpmut3* ($P_{con}$12), H11 CsrB ($P_{LlacO}$; lacI), ColE1 ori, CarbR | This study |
| pTRS075 | Csr-Controlled GFP Buffer Gate: *glgC 5' UTR*-TTGGT-*gfpmut3* ($P_{J23119}$), WT CsrB ($P_{LlacO}$; lacI), ColE1 ori, CarbR | This study |
| pTRS076 | Csr-Controlled GFP Buffer Gate: *glgC 5' UTR*-TTGGT-*gfpmut3* ($P_{J23100}$), WT CsrB ($P_{LlacO}$; lacI), ColE1 ori, CarbR | This study |
| pTRS077 | Csr-Controlled GFP Buffer Gate: *glgC 5' UTR*-TTGGT-*gfpmut3* ($P_{J23107}$), WT CsrB ($P_{LlacO}$; lacI), ColE1 ori, CarbR | This study |
| pTRS078 | Csr-Controlled GFP Buffer Gate: *glgC 5' UTR*-TTGGT-*gfpmut3* ($P_{J23116}$), WT CsrB ($P_{LlacO}$; lacI), ColE1 ori, CarbR | This study |
| pTRS079 | Csr-Controlled GFP Buffer Gate: *glgC 5' UTR*-TTGGT-*gfpmut3* ($P_{J23106}$), WT CsrB ($P_{LlacO}$; lacI), ColE1 ori, CarbR | This study |
| pTRS080 | Csr-Controlled GFP Buffer Gate: *glgC 5' UTR*-TTGGT-*gfpmut3* ($P_{J23110}$), WT CsrB ($P_{LlacO}$; lacI), | This study |

| | | |
|---|---|---|
| | ColE1 ori, CarbR | |
| pTRS081 | Csr-Controlled GFP Buffer Gate: *glgC 5' UTR*-TTGGT-*gfpmut3* (P$_{J23105}$)*,* WT CsrB (P$_{LlacO}$; lacI), ColE1 ori, CarbR | This study |
| pTRS082 | Csr-Controlled GFP Buffer Gate: *glgC 5' UTR*-TTGGT-*gfpmut3* (P$_{J23114}$)*,* WT CsrB (P$_{LlacO}$; lacI), ColE1 ori, CarbR | This study |
| pTRS089 | Csr-Controlled Operon Buffer Gate: *glgC 5' UTR*-TTACA-*gfpmut3*- *glgC 5' UTR*-TTCAT-*mcherry*-*glgC 5' UTR*-TTGGT-*eyfp* (P$_{con}$12)*,* WT CsrB (P$_{LlacO}$; lacI), ColE1 ori, CarbR | This study |
| pTRS090 | Csr-Controlled GFP NOT Gate: y*mdA 5' UTR*-TTGGT-*gfpmut3* (P$_{con}$12)*,* WT CsrB (P$_{LlacO}$; lacI), ColE1 ori, CarbR | This study |
| pTRS093 | Csr-Controlled Mevalonate Operon: *glgC 5' UTR*-TTGGT-*atoB*- *glgC 5' UTR*-TTGGT-*hmgS*- *glgC 5' UTR*-TTGGT-*thmgR* (P$_{con}$12)*,* WT CsrB (P$_{LlacO}$; lacI), ColE1 ori, CarbR | This study |
| pTRS094 | Csr-Controlled GFP NOT Gate: y*mdA 5' UTR*-TTGGT-*gfpmut3* (P$_{con}$12)*,* ColE1 ori, CarbR | This study |
| pTRS095 | OR/NOR Gate: WT CsrB (P$_{LTetO}$: *tetR*), WT CsrB (P$_{LlacO}$: *lacI*), p15a ori, KanR | This study |
| pTRS096 | AND/NAND Gate: WT CsrB (P$_{LTetO}$: *tetR,* P$_{araBAD}$; *araC*), p15a ori, KanR | This study |
| pTRS097 | Transcriptionally controlled synthetic mevalonate operon: pMevO (P$_{LlacO}$: *lacI*), ColE1 ori, CarbR | This study |
| pTRS-Bsub-Switch | Csr-Controlled Buffer Gate with *B. subtilis* compatibility *gfp* (P$_{J23101}$), WT *csrB* (P$_{grac}$), *lacI*, ColE1 ori, rep repA ori, CarbR, CmR | This study |
| pgRNA-csrD | P$_{J23119}$ csrD CRISPR gRNA, ColE1 ori, KanR | This study |
| pMP11 | CRISRP-Cas9 + λ-red plasmid | (6) |
| pHL1756-*glgC-gfp* | *glgC5' UTR* + 100 nt *glgC* CDS + *gfp* CDS (P$_{con}$12), *tetR* (lacI$^{q}$), ColE1 ori, CarbR | (5) |
| pHL600-PLtetO-CsrB | WT c*srB* sRNA (P$_{LTetO}$), p15a, KanR | (7) |
| pCG004 | *gfp* (P$_{grac}$) *lacI,* ColE1 ori, rep repA ori, CarbR, CmR | (4) |
| pRB14 | *csrD* (P$_{VanCC}$) VanR, p15a ori, KanR | This study |
| pAJM773 | *yfp* (P$_{VanCC}$) VanR, p15a ori, KanR | (8) |

**Table S6.** Oligonucleotides used in this study.

| Primer Name | Sequence | Purpose |
|---|---|---|
| rTRS046 | ATTGACATTGTGAGCGGATAACAAGATACTgagcacGTCGACAGGGAGTCAGA | FW Gibson primer to amplify CsrB WT and H8 mutant from pHL600-pTetO-CsrB plasmids and add PLlacO promoter upstream |
| rTRS047 | AATAAAAAAAGGGAGCACTGTAT | RV primer to amplify CsrB WT and H8 from pHL600 |
| rTRS048 | CAGTGCTCCCTTTTTTTATTGCTTTAATCGTACAGGGTAGTAC | FW Gibson primer to amplify pHL1756 backbone with homology to CsrB |
| rTRS049 | ATCCGCTCACAATGTCAATGTTATCCGCTCACATTTATTGTGGAATCCATTATAACCGC | RV Gibson primer to amplify pHL1756 backbone and insert PLlacO promoter upstream of CsrB |
| rTRS050 | tgtggataaccgtattaccg | Sequencing primer for inserting CsrB into pHL1756 |

| | | |
|---|---|---|
| rTRS051 | ATTGACATTGTGAGCGGATAACA ATATAATGgagcacGTCGACAGGG AGTCAGA | FW Gibson primer to amplify WT/H8 CsrB from pHL600 and add trc promoter upstream |
| rTRS052 | ATGAGTAAAGGAGAAGAACTTT | FW Primer to amplify GFP and backbone of pTRS-UTR+GFP plasmids (originally pHL1756) |
| rTRS053 | TTCTCCTTTACTCATGACTAACTC CTTTTTTATCATCTC | RV Gibson primer to insert glgC 5' UTR only intp pHL1756 |
| rTRS056 | TTCTCCTTTACTCATATGAAGACT AACTCCTTTTTTATCATCTC | RV Gibson primer to insert glgC 5' UTR + TTCAT into pHL1756 |
| rTRS057 | TTCTCCTTTACTCATGGGGAGAC TAACTCCTTTTTTATCATCTC | RV Gibson primer to insert glgC 5' UTR + TCCCC into pHL1756 |
| rTRS058 | aaactctcaaggatcttacc | Sequencing primer for 5' UTRs and spacers of pHL1756 |
| rTRS076 | actcgtgcacccaactga | FW Primer to amplify 5' UTR and GFP sequences from pTRS plasmids |
| rTRS077 | tcagttgggtgcacgagt | RV Primer to amplify backbone of pTRS plasmids |
| rTRS078 | GCACCGTCGTTGTTGACA | FW Primer to amplify pTRS-glgC14K-GFP-PLlacO-CsrB WT BB after rrnB Term2 |
| rTRS079 | AAATACATTCAAATATGTATCCG CT | RV Primer to amplify pTRS-glgC14K-GFP-PLlacO-CsrB WT BB after rrnB Term2 |
| rTRS080 | CATTCAAATATGTATCCGCTgtgaa accagtaacgttatacgat | FW Gibsom primer to amplify lacI gene, with homology to rTRS079 |
| rTRS081 | ctggaaagcgggcagtgaGCACCGTCG TTGTTGACA | RV Gibsom primer to amplify lacI gene, with homology to rTRS078 |
| rTRS084 | GATCCTGAGCGGATACATATTTG | RV primer to amplify pTRS-glgC14K-GFP-PLlacO-CsrB WT BB after rrnBT1T2 terminator (homology with rTRS082) |
| rTRS085 | TTCCACAATAAATGTGAGCGGA | FW primer to amplify pTRS-glgC14K-GFP-PLlacO-CsrB WT before PLlacO promoter (homology with rTRS083) |
| rTRS086 | ACATGGTCCTTCTTGAGTTT | Seq Primer to screen for inserts downstream of rrnBT1T2 Terminator |
| rTRS087 | AAACTCAAGAAGGACCATGT | RV Primer to amplify GFP and backbone of pTRS plasmids (rev. comp. of rTRS086) |
| rTRS088 | CGTCAGATGACGTGCCTTTTTTC TTGTGAGCAGTtaagaaaccattattatc atgaca | FW Gibson primer to insert the aspA Terminator downstream of CsrB in the pTRS plasmids |
| rTRS089 | AAAAAAGGCACGTCATCTGACGT GCCTTTTTTATTTGTACTACCCTG TACGATTAAAG | RV Gibson primer to insert the aspA Terminator downstream of CsrB in the pTRS plasmids |
| rTRS090 | TCCTCACTATCGGAGTTAACACA AGGatgCAGCCACTTGATACTAAC GTGAAAAAATAT | 60-mer oligo to knockout csrD from the genome of E. coli |
| rTRS091 | TTCAACCTGGCTGTGCCCCCGTT TTAGAGCTAGAAATAGCAAGTTA AAATAAG | FW Gibson primer to insert the csrD gRNA sequence in to the pgRNA-kan plasmid |
| rTRS092 | GGGGGCACAGCCAGGTTGAAAC TAGTATTATACCTAGGACTGAGC TAGCTG | RV Gibson primer to insert the csrD gRNA sequence in to the pgRNA-kan plasmid |
| rTRS093 | ATCTTTATAGTTCAGGCTCG | FW cPCR primer to test for csrD genomic KO |
| rTRS094 | ATAAATGAGGGTATTCGCAG | RV cPCR primer to test for csrD genomic KO |
| rTRS117 | GGAGTTAGTCttacaATGAGTAAA GGAG | FW SDM Primer to replace 14K RBS in pSW12 with 10K RBS |
| rTRS118 | GGAGTTAGTCttaccATGAGTAAAG G | FW SDM Primer to replace 14K RBS in pSW12 with 8K RBS |

| | | |
|---|---|---|
| rTRS119 | GGAGTTAGTCtttacATGAGTAAAGG | FW SDM Primer to replace 14K RBS in pSW12 with 3K RBS |
| rTRS120 | TTTTTTATCATCTCTGGAACAC | RV SDM Primer to replace 14K RBS in pSW12 with Different RBS (works with rTRS116-119) |
| rTRS121 | TAACCGAAAGTAGTGACAAG | RV Primer to sequence GFP+RBS+lacI in pTRS-GPP plasmids |
| rTRS138 | cccgttgaatatggctcatcattggaaaacgttctt | FW Gibson primer to amplify pSW12 backbone around Carb gene |
| rTRS139 | atgctcgatgagtttttctgataactgtcagaccaagtttact | RV Gibson primer to amplify pSW12 backbone around Carb gene |
| rTRS140 | gttgtttacatttttatgcttgg | FW Primer to amplify glgC14-GFP-lacI-CsrB sequence from pSW12 |
| rTRS141 | cggtaatacggttatccacattcaaatatgtatccgctcat | FW Gibson Primer to amplify pBTRCK backbone |
| rTRS142 | agcataaaaatgtaaacaacatcgctcaatactgaccatt | RV Gibson primer to amplify pBTRCK backbone |
| rTRS143 | TGGACGAGCTGTACAAGTAAgctagcggctgttttgg | FW Gibson primer to amplify pSW12 backbone to insert EYFP |
| rTRS144 | TCCTCGCCCTTGCTCACCATACCAAGACTAACTCCTTTTTTATCA | RV Gibson primer to amplify pSW12 backbone to insert EYFP |
| rTRS145 | TGGATGAACTGTACAAATAAgctagcggctgttttgg | FW Gibson primer to amplify pSW12 backbone to insert mCherry |
| rTRS146 | TCTTCGCCTTTTGAAACCATACCAAGACTAACTCCTTTTTTATCA | RV Gibson primer to amplify pSW12 backbone to insert mCherry |
| rTRS147 | cggtaatacggttatccacaaattcggatatattccgctt | FW Gibson primer to amplify p15a backbone w/homology to SW12-glgC14-mCherry-lacI-CsrB sequence |
| rTRS148 | aaaaatgtaaacaactcgtgatacgcctattttta | RV Gibson primer to amplify p15a backbone w/homology to SW12-glgC14-mCherry-lacI-CsrB sequence |
| rTRS149 | gttgtttacatttttatgcttgg | FW primer to amplify SW12-glgC14-mCherry-lacI-CsrB sequence |
| rTRS150 | CCCGATTTACGTTACGCGCCGTGCTCAGTATCTTGTTATCCG | RV Gibson primer to amplify lacI and lacI promoter region from pSW12 plasmid |
| rTRS151 | CTGACGTCTCGAGCACCGTCCGTTGACACCATCGAATGG | FW Gibson primer to amplify lacI and lacI promoter region from pSW12 plasmid |
| rTRS152 | tgtggataaccgtattaccgAAGAGTTTGTAGAAACGCAAA | RV Gibson primer to amplify rrnB12 terminator and GFP from pSW12 plasmid |
| rTRS153 | GTAACGTAAATCGGGTTTAAATTATGAGTAAAGGAGAAGAACTTT | FW Gibson primer to amplify 14K RBS + GFP-rrnB12 terminator from pSW12 plasmid |
| rTRS154 | cggtaatacggttatccacagaa | FW primer to amplify ColE1 ori and Carb resistance from pSW12 plasmid |
| rTRS155 | ggtgctcgagacgtcag | RV primer to amplify ColE1 ori and Carb resistance from pSW12 plasmid |
| rTRS156 | aaataatactgttgatgggt | Sequencing primer for lacI and lacI promoter on pTRS-PLlacO14-GFP plasmid |
| rTRS157 | gtcaaaccagatcaattcgc | RV Primer to amplify lacI and GFP on pSW12 |
| rTRS158 | gcgaattgatctggtttgac | FW Primer to amplify lacI and GFP on pSW12 |
| rTRS171 | GTCGACAGGGAGTCAGAC | FW Primer to amplify CsrB sRNAs sequences |
| rTRS172 | GTCTGACTCCCTGTCGAC | RV Primer to amplify pSW12 backbone w/homology to rTRS171 |
| rTRS173 | TACAGTGCTCCCTTTTTTTATT | FW Primer to amplify pSW12 backbone w/homology to rTRS047 |
| rTRS174 | cccgttgaatatggctcatcattggaaaacgt | FW primer to amplify pSW12 with homology to kan cassette |

| | | |
|---|---|---|
| rTRS175 | GATAAAAAAGcAGTTAGTCTTGG TATG | FW SDM Primer to change 4th GGA to GCA in glgC 5' UTR (Ta 58) |
| rTRS176 | ATCTCTGGAACACACACAATC | RV SDM Primer to change 4th GGA to GCA in glgC 5' UTR (Ta 58) |
| rTRS177 | CTCTGGCAGGcACCTGCACAC | FW SDM Primer to change 1st GGA to GCA in glgC 5' UTR (Ta 65) |
| rTRS178 | TCGACCTGTGTGGAATCC | RV SDM Primer to change 1st GGA to GCA in glgC 5' UTR (Ta 65) |
| rTRS179 | GTGTTCCAGAcATGATAAAAAAG GAGTTAG | FW SDM Primer to change 3rd GGA to GCA in glgC 5' UTR (Ta 62) |
| rTRS180 | ACACAATCCGTGTGCAGG | RV SDM Primer to change 3rd GGA to GCA in glgC 5' UTR (Ta 62) |
| rTRS181 | CCCGATTTACGTTACGCGCCAgtc gacctgtgtggaa | RV Gibson Primer to add 14K RBS to Pcon12 promoter, homology with rTRS153 |
| rTRS182 | cccgttgaatatggctcatactcttcctttttcaata tt | FW Gibson primer to insert kan cassette into pSW12 plasmids, fixed to remove AmpR fragment |
| rTRS183 | GATAAAAAAGcAGTTAGTCTTGG TATGAG | FW SDM Primer to change 4th GGA to GCA in mutant glgC 5' UTR (Ta 60) |
| rTRS184 | ATGTCTGGAACACACACAATG | RV SDM Primer to change 4th GGA to GCA in mutant glgC 5' UTR (Ta 60) |
| rTRS185 | ctagcggTCTGGCAGGGACCTGC | FW Primer to amplify glgC14-mCherry and keeps NheI binding site |
| rTRS186 | ctagcTTATTTGTACAGTTCATCCA | RV Primer to amplify glgC14-mCherry and keeps NheI binding site |
| rTRS187 | ctagcggctgttttggc | FW Primer to amplify pSW12-glgC14-GFP backbone and keep NheI binding site before rrnB12 terminator |
| rTRS188 | ctagcTTATTTGTAGAGATCATCCA | RV Primer to amplify pSW12-glgC14-GFP backbone and keep NheI binding site after GFP |
| rTRS189 | ctgttttggcggatgaga | FW Primer to amplify entire pSW12-glgC14-GFP backbone |
| rTRS190 | CCTGCCAGAGGTACCTTATTTGT AGAGATCATCCATGC | RV Gibson primer to amplify entire pSW12-glgC14-GFP backbone, add Kpn1 digest site, with homology to rTRS192 |
| rTRS191 | tctcatccgccaaaacagcc | RV Primer to amplify glgC 5' UTR-mCherry CDS |
| rTRS192 | GGTACCTCTGGCAGGGACCTGC A | FW Gibson primer to amplify glgC 5' UTR-mCherry CDS, add Kpn1 digest site |
| rTRS193 | AACCATACATGAATTGCG | RV Sequencing primer in mCherry CDS |
| rTRS194 | gctagcTCTGGCAGGGACCTGCA | FW Primer to amplify EYFP CDS with NheI cut site |
| rTRS195 | GCTTAGCTTACTTGTACAGCTCG TCCATGC | RV Primer to amplify EYFP CDS with Blp1 cut site |
| rTRS196 | GCTGTACAAGTAAGCTAAGCctgtt ttggcggatgag | FW Gibson primer to amplify pSWO backbone with Blp1 cut site |
| rTRS197 | CCCTGCCAGAgctagcTTATTTGTA CAGTTC | RV Gibson primer to amplify pSWO backbone at mCherry CDS with NheI cut site |
| rTRS198 | ATCTTGAAGTTCACCTTG | RV Seq primer in EYFP CDS |
| rTRS199 | cggtaatacggttatccaca | FW Primer to amplify all switch backbones directly upstream of ori (homology to rTRS050) |
| rTRS200 | atgaaaaattgtgtcatcgtc | FW primer to amplify atoB CDS from pMevT |
| rTRS201 | ttaattcaaccgttcaatcac | RV primer to amplify atoB CDS from pMevT |
| rTRS202 | acgatgacacaatttttcatACCAAGACTA ACTCCTTTTTTATC | RV Gibson primer to amplify pSW12 backbone with homology to atoB gene |
| rTRS203 | tgattgaacggttgaattaagctagcggctgtttt gg | FW Gibson primer to amplify pSW12 backbone with homology to atoB gene |

| | | |
|---|---|---|
| rTRS204 | atgaaactctcaactaaact | FW Primer to amplify ERG13 CDS from pMevT |
| rTRS205 | ttattttttaacatcgtaagatc | RV Primer to amplify ERG13 CDS from pMevT |
| rTRS206 | cttacgatgttaaaaaataagctagcggctgttttgg | FW Gibson primer to amplify pSW12 backbone with homology to ERG13 gene |
| rTRS207 | agtttagttgagagtttcatACCAAGACTAACTCCTTTTTTATC | RV Gibson primer to amplify pSW12 backbone with homology to ERG13 gene |
| rTRS208 | actatggttttaaccaataaaac | FW Primer to amplify HMG1 CDS from pMevT |
| rTRS209 | ttaggatttaatgcaggtg | RV Primer to amplify HMG1 CDS from pMevT |
| rTRS210 | cacctgcattaaatcctaagctagcggctgttttgg | FW Gibson primer to amplify pSW12 backbone with homology to HMG1 gene |
| rTRS211 | ttattggttaaaaccatagtACCAAGACTAACTCCTTTTTTATC | RV Gibson primer to amplify pSW12 backbone with homology to HMG1 gene |
| rTRS212 | ctgacgtctcgagcacccaacgcaattaatgtgagttag | FW Gibson primer to remove lacI from pTRS-PLlacO-GFP plasmid (homology with rTRS155) |
| rTRS213 | cggtgaagggcaatcagct | FW Primer to amplify lacI-CsrB backbone region from switch plasmid |
| rTRS198 | ATCTTGAAGTTCACCTTG | RV Seq primer in EYFP CDS |
| rTRS199 | cggtaatacggttatccaca | FW Primer to amplify all switch backbones directly upstream of ori (homology to rTRS050) |
| rTRS200 | atgaaaaattgtgtcatcgtc | FW primer to amplify atoB CDS from pMevT |
| rTRS201 | ttaattcaaccgttcaatcac | RV primer to amplify atoB CDS from pMevT |
| rTRS202 | acgatgacacaatttttcatACCAAGACTAACTCCTTTTTTATC | RV Gibson primer to amplify pSW12 backbone with homology to atoB gene |
| rTRS203 | tgattgaacggttgaattaagctagcggctgttttgg | FW Gibson primer to amplify pSW12 backbone with homology to atoB gene |
| rTRS204 | atgaaactctcaactaaact | FW Primer to amplify ERG13 CDS from pMevT |
| rTRS205 | ttattttttaacatcgtaagatc | RV Primer to amplify ERG13 CDS from pMevT |
| rTRS206 | cttacgatgttaaaaaataagctagcggctgttttgg | FW Gibson primer to amplify pSW12 backbone with homology to ERG13 gene |
| rTRS207 | agtttagttgagagtttcatACCAAGACTAACTCCTTTTTTATC | RV Gibson primer to amplify pSW12 backbone with homology to ERG13 gene |
| rTRS208 | actatggttttaaccaataaaac | FW Primer to amplify HMG1 CDS from pMevT |
| rTRS209 | ttaggatttaatgcaggtg | RV Primer to amplify HMG1 CDS from pMevT |
| rTRS210 | cacctgcattaaatcctaagctagcggctgttttgg | FW Gibson primer to amplify pSW12 backbone with homology to HMG1 gene |
| rTRS211 | ttattggttaaaaccatagtACCAAGACTAACTCCTTTTTTATC | RV Gibson primer to amplify pSW12 backbone with homology to HMG1 gene |
| rTRS212 | ctgacgtctcgagcacccaacgcaattaatgtgagttag | FW Gibson primer to remove lacI from pTRS-PLlacO-GFP plasmid (homology with rTRS155) |
| rTRS213 | cggtgaagggcaatcagct | FW Primer to amplify lacI-CsrB backbone region from switch plasmid |
| rTRS214 | agctgattgcccttcaccg | RV Primer to amplify lacI-GFP backbone region from switch plasmid |
| rTRS215 | gctagctcagtcctaggtataatgctagcTCTGGCAGGGACCTGCA | FW Gibson Primer to replace Pcon12 with J23119 constitutive promoter |
| rTRS216 | taggactgagctagctgtcaagacgtcaggtggcacttt | RV Gibson Primer to replace Pcon12 with J23119 constitutive promoter |
| rTRS217 | gctagctcagtcctaggtacagtgctagcTCTGGCAGGGACCTGCA | FW Gibson Primer to replace Pcon12 with J23100 constitutive promoter |
| rTRS218 | taggactgagctagccgtcaagacgtcaggtggcacttt | RV Gibson Primer to replace Pcon12 with J23100 constitutive promoter |
| rTRS219 | gctagctcagtcctaggtattgtgctagcTCTGGCAGGGACCTGCA | FW Gibson Primer to replace Pcon12 with J23104 constitutive promoter |

| | | |
|---|---|---|
| rTRS220 | taggactgagctagctgtcaagacgtcaggtg gcacttt | RV Gibson Primer to replace Pcon12 with J23104 constitutive promoter |
| rTRS221 | gctagctcagtcctaggtatagtgctagcTCT GGCAGGGACCTGCA | FW Gibson Primer to replace Pcon12 with J23111 (use rTRS222) or J23106 (use rTRS223) constitutive promoter |
| rTRS222 | taggactgagctagccgtcaagacgtcaggtg gcacttt | RV Gibson Primer to replace Pcon12 with J23111 constitutive promoter |
| rTRS223 | taggactgagctagccgtaaagacgtcaggtg gcacttt | RV Gibson Primer to replace Pcon12 with J23106 constitutive promoter or J23110 constitutive promoter or J23105 |
| rTRS224 | gctagctcagtcctaggtacaatgctagcTCT GGCAGGGACCTGCA | FW Gibson Primer to replace Pcon12 with J23110 constitutive promoter |
| rTRS225 | gctagctcagtcctaggtactatgctagcTCT GGCAGGGACCTGCA | FW Gibson Primer to replace Pcon12 with J23105 constitutive promoter |
| rTRS226 | gctagctcagtcctaggtacaatgctagcTCT GGCAGGGACCTGCA | FW Gibson Primer to replace Pcon12 with J23114 constitutive promoter |
| rTRS227 | taggactgagctagccataaagacgtcaggtg gcacttt | RV Gibson Primer to replace Pcon12 with J23114 constitutive promoter |
| rTRS228 | AAAGGCGCGTCCTATTAAGAGA GTTTTTTTATGGTGAGCAAGGGC GA | FW Gibson Primer to insert 600 RBS + EYFP in pTRS-Pcon12 plasmids |
| rTRS229 | TCTTAATAGGACGCGCCTTTAgtc gacctgtgtggaatc | RV Gibson Primer to amplify bb of pTRS-Pcon12 plasmid and insert 600 RBS |
| rTRS230 | AATAGGGTAAATTAAGGACTATA GGATGGTTTCAAAAGGCGAAG | FW Gibson Primer to insert 11k RBS + mCherry in pTRS-Pcon12 plasmids |
| rTRS231 | AGTCCTTAATTTACCCTATTAgtcg acctgtgtggaatc | RV Gibson Primer to amplify bb of pTRS-Pcon12 plasmid and insert 11k RBS |
| rTRS232 | ggactctggggttcgtcgagcaccgtcgttgtt | FW Gibson Primer to amplify Pcon12 + glgC 5' UTR 14K RBS + GFPmut3, lacI promoter-lacI, PLlacO-CsrB WT |
| rTRS233 | ttcatccgcttattatcacttgtggataaccgtatta ccgcc | RV Gibson Primer to amplify Pcon12 + glgC 5' UTR 14K RBS + GFPmut3, lacI promoter-lacI, PLlacO-CsrB WT |
| rTRS234 | agtgataataagcggatgaatgg | FW Primer to amplify pBBR1 backbone |
| rTRS235 | cgaaccccagagtcccg | RV Primer to amplify pBBR1 backbone |
| rTRS236 | aaatatgtatccgctcagga | FW Primer to amplify directly upstream of lacI promoter, remove partial Amp Promoter |
| rTRS237 | tcctgagcggatacatatttgagtttgtagaaac gcaaaaag | RV Gibson Primer to amplify into rrnB12 terminator and GFP, remove partial Amp Promoter |
| rTRS238 | gcattaaatcctaatagcggctgttttggcgg | FW Gibson Primer to amplify pTRS034 backbone with homology to HMG1 |
| rTRS239 | gggattacTCTGGCAGGGACCTGCA | FW Primer to amplify glgC14+HMG1 |
| rTRS240 | ccgctattaggatttaatgcaggtgacgga | RV Primer to amplify glgC14+HMG1 |
| rTRS241 | cagctaaTCTGGCAGGGACCTGCA | FW Primer to amplify glgC14+ERG13 |
| rTRS242 | CCTGCCAGAgtaatcccgctagcttatttttt aacatcgt | RV Gibson primer to amplify glgC14+ERG13 with homology to HMG1 insert |
| rTRS243 | CCTGCCAGAttagctgtttaattcaaccgtt caatcacca | RV Gibson primer to amplify glgC14+atoB with homology to ERG13 insert |
| rTRS244 | aaaatatgagtttagcccc | FW sequencing primer for atoB in pTRS operon plasmid |
| rTRS245 | ttatcgtagatggcaatatc | RV sequencing primer for ERG13 and atoB1 in pTRS operon plasmid |
| rTRS246 | tttaaccgataagaacatt | FW sequencing primer for ERG13 pTRS operon plasmid |

| | | |
|---|---|---|
| rTRS247 | attaggtgatactacgaga | FW Sequencing primer for HMG1 in pTRS operon plasmid |
| rTRS248 | ttttatttgatgcctggc | RV Sequencing primer from rrnB12 terminator in pTRS034 and other switch plasmids |
| rTRS249 | ttcgatggtgtcaacgtcgctcaatactgaccatt t | RV Gibson primer to insert lacI-PLlacO14-GFP into pBTRCK backbone |
| rTRS250 | acgatgacacaatttttcatAATTTAAACC CGATTTACGTTACG | RV Gibson primer to amplify pTRS-PLlacO14 backbone with homology to atoB |
| rTRS251 | ctaggtatagtgctagcTCTGGCAGGGA CCTGCAC | FW Gibson Primer to replace J23100 promoter with J23106 promoter via SDM in switch plasmid (Ta 67) |
| rTRS252 | gactgagctagccgtaaaGACGTCAGG TGGCACTTTTC | RV Gibson Primer to replace J23100 promoter with J23106 promoter via SDM in switch plasmid (Ta 67) |
| rTRS253 | ctaggtacaatgctagcTCTGGCAGGGA CCTGCAC | FW Gibson Primer to replace J23100 promoter with J23114 promoter via SDM in switch plasmid (Ta 67) |
| rTRS254 | gactgagctagccataaaGACGTCAGG TGGCACTTTTC | RV Gibson Primer to replace J23100 promoter with J23114 promoter via SDM in switch plasmid (Ta 67) |
| rTRS255 | atagcagaaagtcaaaagcc | RV Primer to amplify pBTRCK bb |
| rTRS256 | gcaaaacctttcgcggtat | FW Primer to amplify part of lacIq promoter and lacI gene |
| rTRS257 | ctaggtactatgctagcTCTGGCAGGGA CCTGCAC | FW Gibson Primer to replace J23100 promoter with J23105 promoter via SDM in switch plasmid (Ta 67) |
| rTRS258 | gactgagctagccgtaaaGACGTCAGG TGGCACTTTTC | RV Gibson Primer to replace J23100 promoter with J23105 promoter via SDM in switch plasmid (Ta 67) |
| rTRS259 | ctaggtattatgctagcTCTGGCAGGGA CCTGCAC | FW Gibson Primer to replace J23100 promoter with J23107 promoter via SDM in switch plasmid (Ta 67) |
| rTRS260 | ggctgagctagccgtaaaGACGTCAGG TGGCACTTTTC | RV Gibson Primer to replace J23100 promoter with J23107 promoter via SDM in switch plasmid (Ta 67) |
| rTRS261 | ctagggactatgctagcTCTGGCAGGGA CCTGCAC | FW Gibson Primer to replace J23100 promoter with J23116 promoter via SDM in switch plasmid (Ta 67) |
| rTRS262 | gactgagctagctgtcaaGACGTCAGGT GGCACTTTTC | RV Gibson Primer to replace J23100 promoter with J23116 promoter via SDM in switch plasmid (Ta 67) |
| rTRS263 | ctaggtacaatgctagcTCTGGCAGGGA CCTGCAC | FW Gibson Primer to replace J23100 promoter with J23110 promoter via SDM in switch plasmid (Ta 67) |
| rTRS264 | gactgagctagccgtaaaGACGTCAGG TGGCACTTTTC | RV Gibson Primer to replace J23100 promoter with J23110 promoter via SDM in switch plasmid (Ta 67) |
| rTRS265 | ATGGTTTTAACCAATAAAACAG | FW Primer to remove threonine codon from HMG1 gene in pSW12-MevT plasmid (Ta 58) |
| rTRS266 | ACCAAGACTAACTCCTTTTTATC | FW Primer to remove threonine codon from HMG1 gene in pSW12-MevT plasmid (Ta 58) |
| rTRS267 | ataccaatggcaactacaga | FW Primer to amplify pSW12-MevT to remove threonine codon |
| rTRS268 | gtttgttggaacaatcaaagat | RV Primer to amplify pSW12-MevT to remove threonine codon |

| | | |
|---|---|---|
| rTRS269 | AGTTCTTCTCCTTTACTCATgtaaaGACTAACTCCTTTTTTATCATCTCTG | RV Gibson Primer to replace 14K RBS with 3K RBS for GFP |
| rTRS270 | TCTTCGCCTTTTGAAACCATTGTAAGACTAACTCCTTTTTTATCATCTCTG | RV Gibson Primer to replace 14K RBS with 10K RBS for mCherry |
| rTRS271 | ATGGTTTCAAAAGGCGAAGA | FW Primer to amplify mCherry with homology to rTRS270 |
| rTRS272 | ACCCTCAAGGATGACTAATCATTGAGGAAATAGAATAATTGGTATGAGTAAAGGAGAAG | FW Gibson Primer to insert ymdA 5' UTR into pSW12 plasmid |
| rTRS273 | GATTAGTCATCCTTGAGGGTTGTGTTATCCATACTTTCAagtcgacctgtgtggaat | RV Gibson Primer to insert ymdA 5' UTR into pSW12 plasmid |
| rTRS278 | CTTGGAGACGGACAGGTTTATG | FW qPCR Primer for MevT Switch Operon |
| rTRS279 | CACGGGTAATTCCGTACTCTTT | RV qPCR Primer for MevT Switch Operon |
| rTRS286 | ctcagtcctaggtattatgctagcTCTGGCAGGGACCTGC | FW Gibson Primer to insert glgC-GFP sequence with J23101 Promoter into pGB004 |
| rTRS287 | aaacatcttatgcttacattAAGAGTTTGTAGAAACGCAAAAAGG | RV Gibson Primer to insert glgC-GFP sequence with J23101 Promoter into pGB004 |
| rTRS288 | aatgtaagcataagatgtttttc | FW Gibson Primer to amplify AmpR-ColE1-cat-lacI bb of pGB004 (homology to rTRS287) |
| rTRS289 | ccttcctcctttaattggg | RV Gibson Primer to amplify AmpR-ColE1-cat-lacI bb of pGB004 (homology to rTRS290) |
| rTRS290 | tcccaattaaaggaggaaggGTCGACAGGGAGTCAGAC | FW Gibson Primer to amplify CsrB and AspA Terminator into pGB004 (homology to rTRS289) |
| rTRS291 | tttcagttgcagacaaagattgtggataaccgtattaccg | RV Gibson Primer to amplify CsrB and AspA Terminator into pGB004 (homology to rTRS292) |
| rTRS292 | atctttgtctgcaactgaaaa | FW Primer to amplify RepA bb of pGB004 (homology to rTRS291) |
| rTRS293 | taatacctaggactgagctagctgtaaagactcagatgttaaagtgtttg | RW Gibson Primer to amplify RepA bb of pGB004 and insert J23101 promoter (homology to rTRS286) |
| rTRS294 | taatactagagccaggcat | FW Primer to amplify pGB004 starting after original GFP |
| rTRS295 | cccaattaaaggaggaaggGGCGCGTAACGTAAATCG | FW Gibson Primer to amplify 14K RBS +GFP from pTRS-PLlacO14 |
| rTRS296 | atgcctggctctagtattaTTATTTGTAGAGATCATCCATGCC | RV Gibson Primer to amplify 14K RBS +GFP from pTRS-PLlacO14 |
| rTRS315 | CAGTGCTCCCTTTTTTTATTctgttttggcggatgagaga | FW Gibson primer to amplify pTRS063 backbone except GFP seq |
| rTRS316 | gccattcgatggtgtcaacgtcgtgatacgcctatttttatagg | RV Gibson primer to amplify pTRS063 backbone except GFP seq |
| rTRS317 | cggtaatacggttatccaca | FW Primer to amplify pTRS063 backbone of ori and KanR |
| rTRS318 | tctctcatccgccaaaacagTTAAGACCCACTTTCACATTTAA | RV Gibson primer to amplify pTRS095 at TetR |
| oRB150 | ATGAGATTAACGACGAAAT | CsrD Forward from genome |
| oRB151 | TTAAACCGAGTATCTTTGT | CsrD Reverse from genome |
| oRB189 | ATTTCGTCGTTAATCTCATctagtatttcccctctttctcta | pAJM773 Backbone Reverse to add CsrD |
| oRB190 | ACAAAGATACTCGGTTTAActcggtaccaaattccaga | pAJM773 Backbone Forward to add CsrD |
| secA_F_payne_qpcr | cgtaccggtgaaggaaaaac | Fw qPCR Primer for secA housekeeping gene |

| | | |
|---|---|---|
| secA_R_payne_qpcr | gtagtcgttgacggtaactacg | Rv qPCR Primer for secA housekeeping gene |
| mCherry gBlock | ATGGTTTCAAAAGGCGAAGAAGACAACATGGCGATTATCAAGGAATTTATGCGTTTCAAGGTCCACATGGAAGGCAGCGTCAATGGTCACGAATTTGAAATTGAAGGCGAAGGTGAAGGCCGTCCGTATGAAGGCACCCAGACGGCAAAACTGAAGGTCACCAAAGGCGGTCCGCTGCCGTTTGCTTGGGATATTCTGTCACCGCAATTCATGTATGGTTCGAAAGCGTACGTTAAGCATCCGGCCGATATCCCGGACTATCTGAAACTGTCCTTTCCGGAAGGCTTCAAATGGGAACGTGTTATGAACTTCGAAGATGGCGGTGTGGTTACCGTCACGCAGGATAGCTCTCTGCAAGACGGTGAATTTATTTATAAAGTGAAGCTGCGCGGCACCAATTTCCCGAGCGATGGTCCGGTTATGCAGAAAAAGACGATGGGCTGGGAAGCGAGTTCCGAACGTATGTACCCGGAAGACGGTGCCCTGAAAGGCGAAATCAAGCAGCGCCTGAAACTGAAGGATGGCGGTCACTATGACGCAGAAGTGAAAACCACGTACAAGGCTAAAAAGCCGGTCCAACTGCCGGGTGCATACAACGTGAACATCAAGCTGGATATCACCAGCCATAACGAAGACTATACGATCGTTGAACAGTACGAACGTGCAGAAGGCCGCCACTCTACCGGCGGTATGGATGAACTGTACAAATAA | mCherry gBlock E. coli codon optomized |
| EYFP gBlock | ATGGTGAGCAAGGGCGAGGAGCTGTTCACCGGGGTGGTGCCCATCCTGGTCGAGCTGGACGGCGACGTAAACGGCCACAAGTTCAGCGTGTCCGGCGAGGGCGAGGGCGATGCCACCTACGGCAAGCTGACCCTGAAGTTCATCTGCACCACCGGCAAGCTGCCCGTGCCCTGGCCCACCCTCGTGACCACCTTCGGCTACGGCCTGCAATGCTTCGCCCGCTACCCCGACCACATGAAGCTGCACGACTTCTTCAAGTCCGCCATGCCCGAAGGCTACGTCCAGGAGCGCACCATCTTCTTCAAGGACGACGGCAACTACAAGACCCGCGCCGAGGTGAAGTTCGAGGGCGACACCCTGGTGAACCGCATCGAGCTGAAGGGCATCGACTTCAAGGAGGACGGCAACATCCTGGGGCACAAGCTGGAGTACAACTACAACAGCCACAACGTCTATATCATGGCCGACAAGCAGAAGAACGGCATCA | EYFP gBlock |

| | | |
|---|---|---|
| | AGGTGAACTTCAAGATCCGCCA<br>CAACATCGAGGACGGCAGCGTG<br>CAGCTCGCCGACCACTACCAGC<br>AGAACACCCCCATCGGCGACGG<br>CCCCGTGCTGCTGCCCGACAAC<br>CACTACCTGAGCTACCAGTCCG<br>CCCTGAGCAAAGACCCCAACGA<br>GAAGCGCGATCACATGGTCCTG<br>CTGGAGTTCGTGACCGCCGCCG<br>GGATCACTCTCGGCATGGACGA<br>GCTGTACAAGTAA | |
| Tet_CsrB_<br>gBlk | cgttgacaccatcgaatggcgcaaaacctttcg<br>cggtatggcatgatagcgcccggaagagagtc<br>aattcaggATGTCTAGATTAGATAAA<br>AGTAAAGTGATTAACAGCGCATT<br>AGAGCTGCTTAATGAGGTCGGA<br>ATCGAAGGTTTAACAACCCGTAA<br>ACTCGCCCAGAAGCTAGGTGTA<br>GAGCAGCCTACATTGTATTGGCA<br>TGTAAAAAATAAGCGGGCTTTGC<br>TCGACGCCTTAGCCATTGAGATG<br>TTAGATAGGCACCATACTCACTT<br>TTGCCCTTTAGAAGGGGAAAGCT<br>GGCAAGATTTTTTACGTAATAAC<br>GCTAAAAGTTTTAGATGTGCTTT<br>ACTAAGTCATCGCGATGGAGCA<br>AAAGTACATTTAGGTACACGGCC<br>TACAGAAAAACAGTATGAAACTC<br>TCGAAAATCAATTAGCCTTTTTAT<br>GCCAACAAGGTTTTTCACTAGAG<br>AATGCATTATATGCACTCAGCGC<br>TGTGGGGCATTTTACTTTAGGTT<br>GCGTATTGGAAGATCAAGAGCAT<br>CAAGTCGCTAAAGAAGAAAGGG<br>AAACACCTACTACTGATAGTATG<br>CCGCCATTATTACGACAAGCTAT<br>CGAATTATTTGATCACCAAGGTG<br>CAGAGCCAGCCTTCTTATTCGGC<br>CTTGAATTGATCATATGCGGATT<br>AGAAAAACAACTTAAATGTGAAA<br>GTGGGTCTTAAgctagcggTCCCTA<br>TCAGTGATAGAGATTGACATCCC<br>TATCAGTGATAGAGATACTGAGC<br>ACGTCGACAGGGAGTCAGACAA<br>CGAAGTGAACATCAGGATGATG<br>ACACTTCTGCAGGACACACCAG<br>GATGGTGTTTCAGGGAAAGGCT<br>TCTGGATGAAGCGAAGAGGATG<br>ACGCAGGACGCGTTAAAGGACA<br>CCTCCAGGATGGAGAATGAGAA<br>CCGGTCAGGATGATTCGGTGGG<br>TCAGGAAGGCCAGGGACACTTC<br>AGGATGAAGTATCACATCGGGG<br>TGGTGTGAGCAGGAAGCAATAG<br>TTCAGGATGAACGATTGGCCGC<br>AAGGCCAGAGGAAAAGTTGTCA | gBlock to insert tet regulated CsrB expression for multi-input gates |

| | | |
|---|---|---|
| | AGGATGAGCAGGGAGCAACAAA<br>AGTAGCTGGAATGCTGCGAAAC<br>GAACCGGGAGCGCTGTGAATAC<br>AGTGCTCCCTTTTTTTATT | |
| araC_pBAD_gBlk | ctgttttggcggatgagagaagattttcagcctg<br>atacagattaaatcagaacgcagaagcggtct<br>gataaaacagaatttgcctggcggcagtagcg<br>cggtggtcccacctgaccccatgccgaactca<br>gaagtgaaacgccgtagcgccgatggtagtgt<br>ggggtctccccatgcgagagtagggaactgcc<br>aggcatcaaataaaacgaaaggctcagtcga<br>aagactgggcctttcgttttatctgttgtttgtcggt<br>gaacgctctcctgagtaggacaaatccgccgg<br>gagcggatttgaacgttgcgaagcaacggccc<br>ggagggtggCGGGCAGGACGCCCG<br>CCATAAACTGCCAGGCATCAAAT<br>TAAGCAGAAGGCCATCCTGACG<br>GATGGCCTTTTTGCGTTTCTACA<br>AACTCTTTTGTTTATTTTTCTAAA<br>TACATTCAAATATGTATCCGCTC<br>AGGATCgcgcaacgcaattaatgtgagtta<br>gcgcgaattgatctggtttgacagcttatcatTT<br>CCACAAGTCAAAAACCTCCGACC<br>GGAGGCTTTTGACTTGAGGGGG<br>ATCCGGCACATTTCCCCGAAAAG<br>TGCCACCTGCATCGATTTATTAT<br>GACAACTTGACGGCTACATCATT<br>CACTTTTTCTTCACAACCGGCAC<br>GGAACTCGCTCGGGCTGGCCCC<br>GGTGCATTTTTTAAATACCCGCG<br>AGAAGTAGAGTTGATCGTCAAAA<br>CCAACATTGCGACCGACGGTGG<br>CGATAGGCATCCGGGTGGTGCT<br>CAAAAGCAGCTTCGCCTGGCTG<br>ATACGTTGGTCCTCGCGCCAGC<br>TTAAGACGCTAATCCCTAACTGC<br>TGGCGGAAAAGATGTGACAGAC<br>GCGACGGCGACAAGCAAACATG<br>CTGTGCGACGCTGGCGATATCA<br>AAATTGCTGTCTGCCAGGTGATC<br>GCTGATGTACTGACAAGCCTCG<br>CGTACCCGATTATCCATCGGTG<br>GATGGAGCGACTCGTTAATCGC<br>TTCCATGCGCCGCAGTAACAATT<br>GCTCAAGCAGATTTATCGCCAGC<br>AGCTCCGAATAGCGCCCTTCCC<br>CTTGCCCGGCGTTAATGATTTGC<br>CCAAACAGGTCGCTGAAATGCG<br>GCTGGTGCGCTTCATCCGGGCG<br>AAAGAACCCCGTATTGGCAAATA<br>TTGACGGCCAGTTAAGCCATTCA<br>TGCCAGTAGGCGCGCGGACGAA<br>AGTAAACCCACTGGTGATACCAT<br>TCGCGAGCCTCCGGATGACGAC<br>CGTAGTGATGAATCTCTCCTGGC<br>GGGAACAGCAAAATATCACCCG | gBlock to place csrB under araBAD promoter regulation |

| | |
|---|---|
| | GTCGGCAAACAAATTCTCGTCCC<br>TGATTTTTCACCACCCCCTGACC<br>GCGAATGGTGAGATTGAGAATAT<br>AACCTTTCATTCCCAGCGGTCGG<br>TCGATAAAAAAATCGAGATAACC<br>GTTGGCCTCAATCGGCGTTAAAC<br>CCGCCACCAGATGGGCATTAAA<br>CGAGTATCCCGGCAGCAGGGGA<br>TCATTTTGCGCTTCAGCCATACT<br>TTTCATACTCCCGCCATTCAGAG<br>AAGAAACCAATTGTCCATATTGC<br>ATCAGACATTGCCGTCACTGCGT<br>CTTTTACTGGCTCTTCTCGCTAA<br>CCAAACCGGTAACCCCGCTTATT<br>AAAAGCATTCTGTAACAAAGCGG<br>GACCAAAGCCATGACAAAAACG<br>CGTAACAAAAGTGTCTATAATCA<br>CGGCAGAAAAGTCCACATTGATT<br>ATTTGCACGGCGTCACACTTTGC<br>TATGCCATAGCATTTTTATCCATA<br>AGATTAGCGGATCCTACCTGAC<br>GCTTTTTATCGCAACTCTCTACT<br>GTTTCTCCAT |

**Table S7.** Engineered 5’ UTRs and CsrB sRNA sequences used in this study. 5-nt spacers are in red, and mutations to the *csrB* sRNA are in blue.

| Part | Sequence | Reference |
|---|---|---|
| Buffer Gate 5’ UTR + 14K RBS | TCTGGCAGGGACCTGCACACGGATTGTGTGTGTTCCAGAGA<br>TGATAAAAAAGGAGTTAGTCTTGGT | This study |
| Buffer Gate 5’ UTR + 10K RBS | TCTGGCAGGGACCTGCACACGGATTGTGTGTGTTCCAGAGA<br>TGATAAAAAAGGAGTTAGTCTTACA | This study |
| Buffer Gate 5’ UTR + 5K RBS | TCTGGCAGGGACCTGCACACGGATTGTGTGTGTTCCAGAGA<br>TGATAAAAAAGGAGTTAGTCTTCAT | This study |
| Buffer Gate 5’ UTR + 3K RBS | TCTGGCAGGGACCTGCACACGGATTGTGTGTGTTCCAGAGA<br>TGATAAAAAAGGAGTTAGTCTTTAC | This study |
| NOT Gate 5’ UTR + 14K RBS | TGAAAGTATGGATAACACAACCCTCAAGGATGACTAATCATT<br>GAGGAAATAGAATAATTGGT | This study |
| WT *csrB* sRNA | GTCGACAGGGAGTCAGACAACGAAGTGAACATCAGGATGAT<br>GACACTTCTGCAGGACACACCAGGATGGTGTTTCAGGGAAA<br>GGCTTCTGGATGAAGCGAAGAGGATGACGCAGGACGCGTT<br>AAAGGACACCTCCAGGATGGAGAATGAGAACCGGTCAGGAT<br>GATTCGGTGGGTCAGGAAGGCCAGGGACACTTCAGGATGA<br>AGTATCACATCGGGGTGGTGTGAGCAGGAAGCAATAGTTCA<br>GGATGAACGATTGGCCGCAAGGCCAGAGGAAAAGTTGTCAA<br>GGATGAGCAGGGAGCAACAAAAGTAGCTGGAATGCTGCGA<br>AACGAACCGGGAGCGCTGTGAATACAGTGCTCCCTTTTTTT<br>ATT | https://ecocyc.org/ |
| L2 *csrB* sRNA | GTCGACAGGGAGTCAGACAACGAAGTGAACATCAGGATGAT<br>GACACTTCTGCAGGACACACCAGGATGGTGTTTCAGGGAAA<br>GGCATCGGGGTGGTGCGAAGAGGATGACGCAGGACGCGTT<br>AAAGGACACCTCCAGGATGGAGAATGAGAACCGGTCAGGAT<br>GATTCGGTGGGTCAGGAAGGCCAGGGACACTTCAGGATGA<br>AGTATCACATCGGGGTGGTGTGAGCAGGAAGCAATAGTTCA | (9) |

| | | |
|---|---|---|
| | GGATGAACGATTGGCCGCAAGGCCAGAGGAAAAGTTGTCAA<br>GGATGAGCAGGGAGCAACAAAAGTAGCTGGAATGCTGCGA<br>AACGAACCGGGAGCGCTGTGAATACAGTGCTCCCTTTTTTT<br>ATT | |
| H11 *csrB* sRNA | GTCGACAGGGAGTCAGACAACGAAGTGAACATCAGGATGAT<br>GACACTTCTGCAGGACACACCAGGATGGTGTTTCAGGGAAA<br>GGCTTCTGGATGAAGCGAAGAGGATGACGCAGGGAGCGTT<br>AAAGGACACCTCCAGGATGGAGAATGAGAACCGGTCAGGAT<br>GATTCGGTGGGTCAGGAAGGCCAGGGACACTTCAGGATGA<br>AGTATCACATCAGGGAGGTGTGAGCAGGGAGCAATAGTTCA<br>GGATGAACGATTGGCCAGGGAGGCCAGAGGAAAAGTTGTC<br>AAGGATGAGCAGGGAGCAACAAAAGTAGCTGGAATGCTGC<br>GAAACGAACCGGGAGCGCTGTGAATACAGTGCTCCCTTTTT<br>TTATT | (9) |
| L3 *csrB* sRNA | GTCGACAGGGAGTCAGACAACGAAGTGAACATCAGGATGAT<br>GACACTTCTGCAGGACACACCAGGATGGTGTTTCAGGGAAA<br>GGCTTCTGGATGAAGCGAAGAGGATGACGCAGGACGCGTT<br>AAAGGACACCTCCAGGATGGAGAATGAGAACCGGTCAGGAT<br>GATTCGGTGGGTCAGGAAGGCCAGGGACACTTCAGGATGA<br>AGTATCACATCGGGGTGGTGTGAGCAGGAAGCAATAGTTCA<br>GGATGAACGATTGGCCGCAAGGCCAGAGGAAAAGTTGTCAA<br>GGATGAGCGGGGTGCAACAAAAGTAGCTGGAATGCTGCGA<br>AACGAACCGGGAGCGCTGTGAATACAGTGCTCCCTTTTTTT<br>ATT | (9) |
| H4 *csrB* sRNA | GTCGACAGGGAGTCAGACAACGAAGTGAACATCAGGATGAT<br>GACACTTCTGCAGGACACACCAGGATGGTGTTTCAGGGAAA<br>GGCTTCTGGATGAAGCGAAGAGGATGACGCAGGACGCGTT<br>AAAGGACACCTCCAGGATGGAGAATGAGAACCGGTCAGGAT<br>GATTCGGTGGGTCAGGAAGGCCAGGGACACTTCAGGATGA<br>AGTATCACATCGGGGTGGTGTGAGCAGGAAGCAATAGTTCA<br>GGATGAACGATTGGCCAGGGAGGCCAGAGGAAAAGTTGTC<br>AAGGATGAGCAGGGAGCAACAAAAGTAGCTGGAATGCTGC<br>GAAACGAACCGGGAGCGCTGTGAATACAGTGCTCCCTTTTT<br>TTATT | (9) |
| H8 *csrB* sRNA | GTCGACAGGGAGTCAGACAACGAAGTGAACATCAGGATGAT<br>GACACTTCTGCAGGACACACCAGGATGGTGTTTCAGGGAAA<br>GGCTTCTGGATGAAGCGAAGAGGATGACGCAGGACGCGTT<br>AAAGGACACCTCCAGGATGGAGAATGAGAACCGGTCAGGAT<br>GATTCGGTGGGTCAGGAAGGCCAGGGACACTTCAGGATGA<br>AGTATCACATCAGGGAGGTGTGAGCAGGAAGCAATAGTTCA<br>GGATGAACGATTGGCCAGGGAGGCCAGAGGAAAAGTTGTC<br>AAGGATGAGCAGGGAGCAACAAAAGTAGCTGGAATGCTGC<br>GAAACGAACCGGGAGCGCTGTGAATACAGTGCTCCCTTTTT<br>TTATT | (9) |

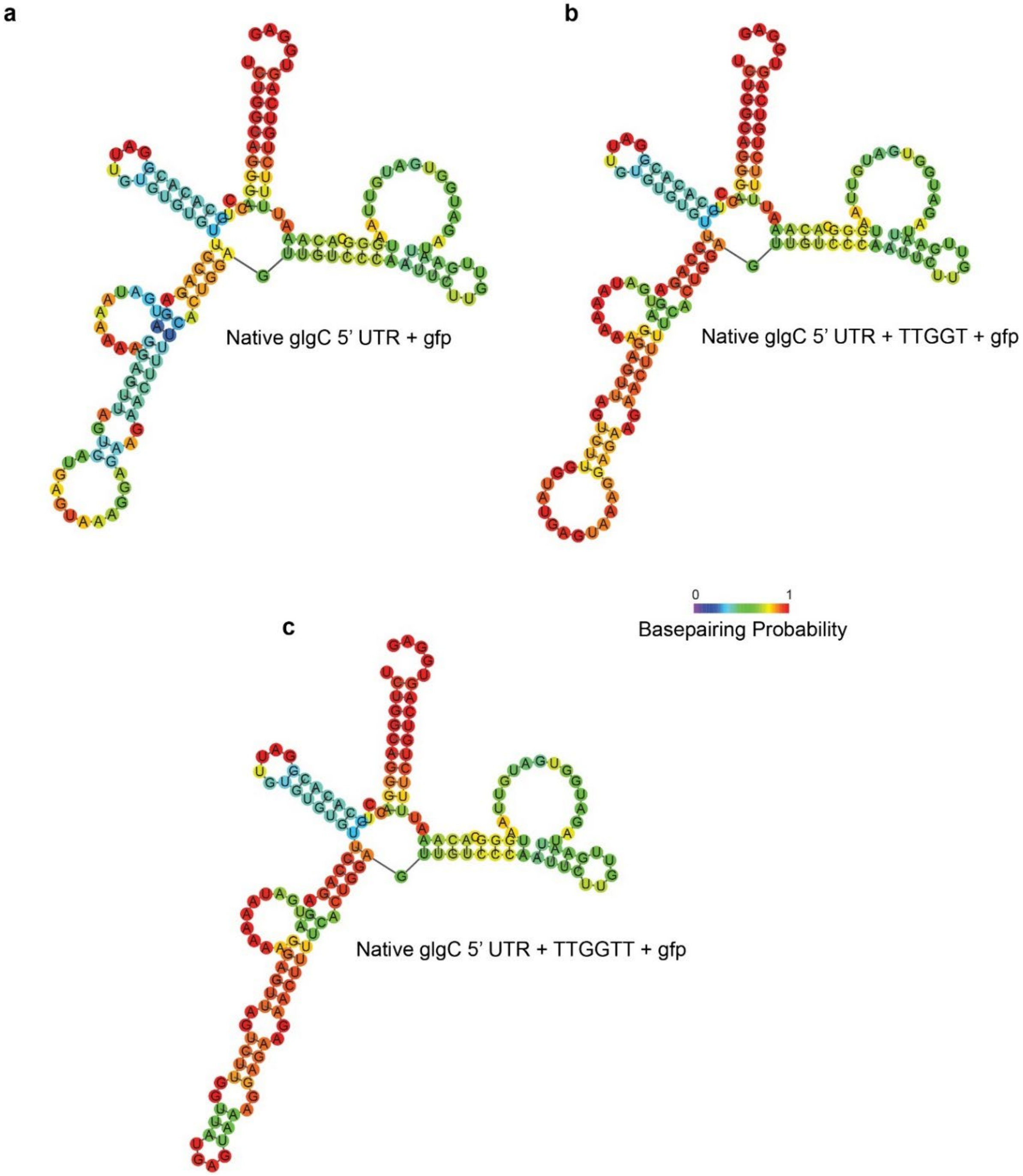

**Supplementary Figure S1. Predicted secondary structures of the wild type and engineered *glgC* 5' UTR sequences to evaluate optimal spacer length.** a) Predicted secondary structure of the wild type *glgC* 5' UTR fused to the first 100 nucleotides (nts) of the *gfp* CDS. b) Predicted secondary structure of the wild type *glgC* 5' UTR with TTGGT appended to the 3' UTR of the sequence fused to the first 100 nucleotides (nts) of the *gfp* CDS. c) Predicted secondary structure of the wild type *glgC* 5' UTR with TTGGTT appended to the 3' UTR of the sequence fused to the first 100 nucleotides (nts) of the *gfp* CDS.

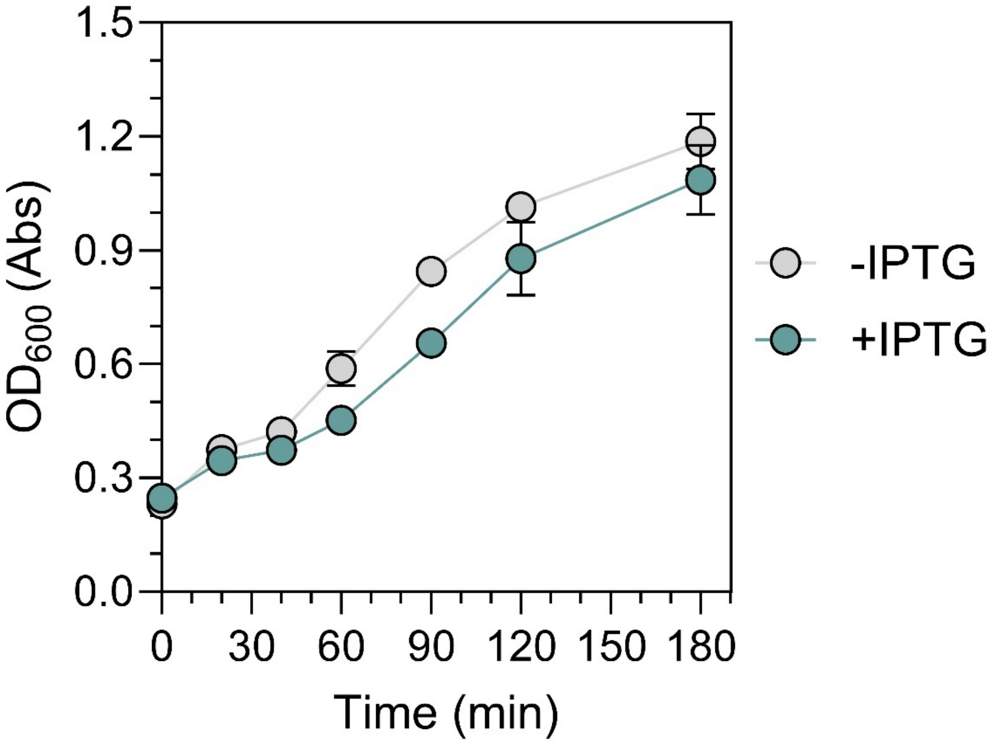

**Supplementary Figure S2. Growth Curve of induced and uninduced Csr-regulated Buffer Gate containing cultures.** $OD_{600}$ of induced (blue line) and uninduced (grey line) cultures after induction of the Csr Buffer Gate with 500 µM IPTG.

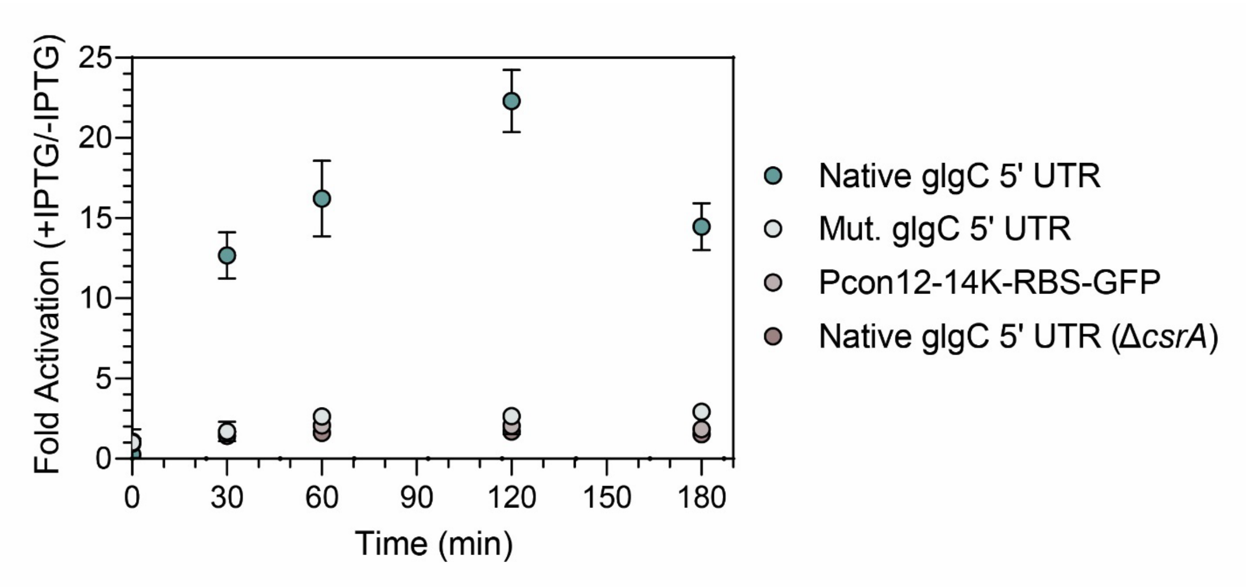

**Supplementary Figure S3. Confirming *in vivo* CsrA repression of the engineered *glgC* 5' UTR-GFP fusion construct.** Time-course of the ratio between the GFP fluorescence of the induced and uninduced Csr-Controlled Buffer Gate cultures (Fold Activation) for the engineered *glgC* 5' UTR fusion sequence (dark blue circles), an engineered *glgC* 5' UTR fusion sequence with all CsrA binding sites mutated (light blue circles), constitutively expressed GFP transcript (light red circles), and the engineered *glgC* 5' UTR fusion sequence expressed in a CsrA deletion strain (dark red circles).

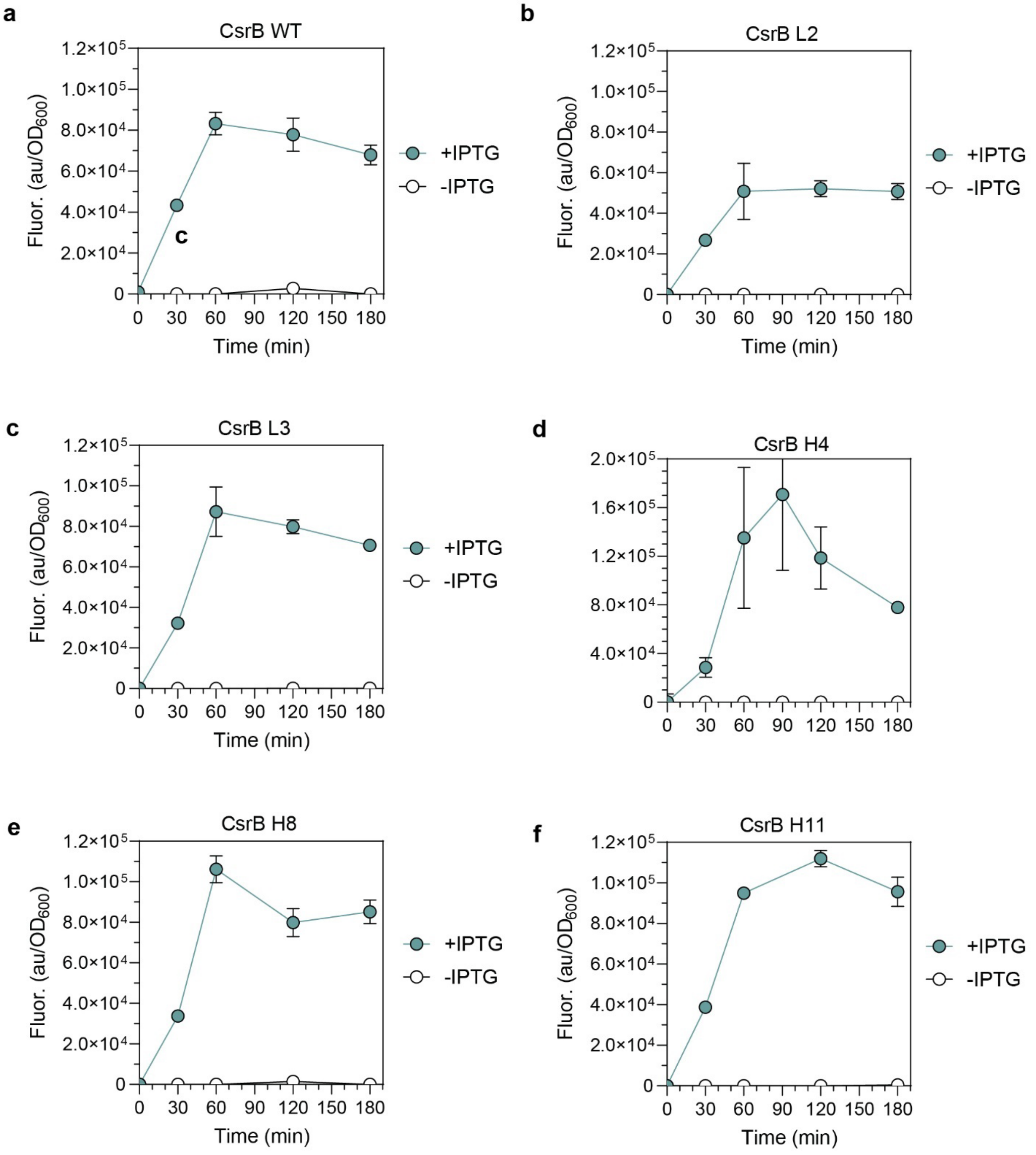

**Supplementary Figure S4. Evaluating Csr-Controlled Buffer Gate Performance using mutant CsrB sRNA sequences.** Time-course experiments measuring $OD_{600}$-normalized GFP fluorescence for a) WT CsrB sRNA Sequence and multiple CsrB sRNA mutant sequences derived from Leistra et al. 2017 *ACS Syn. Bio.* specifically, b) CsrB L2, c) CsrB L3, d) CsrD H4, e) CsrB H8, and f) CsrB H11.

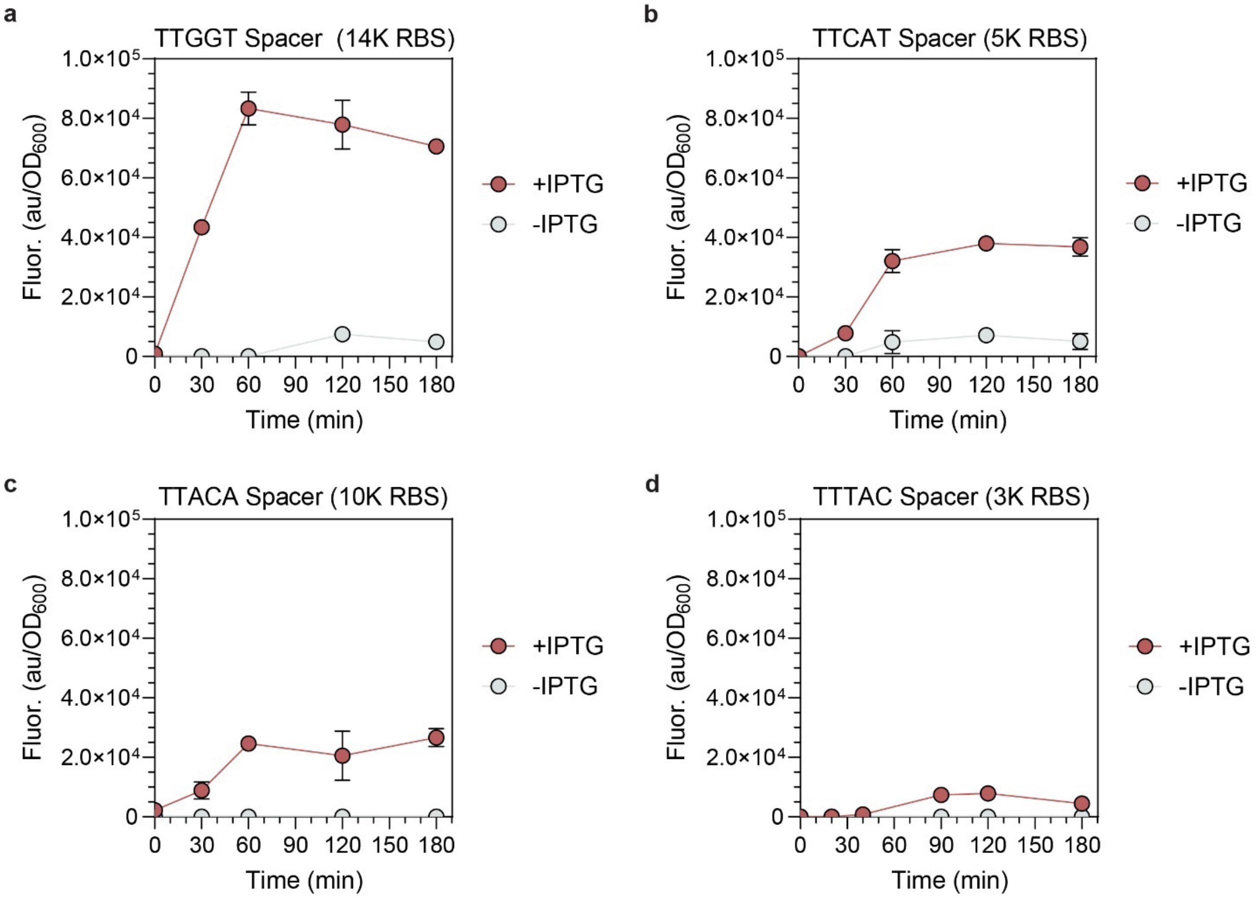

**Supplementary Figure S5. Evaluating Csr-Controlled Buffer Gate Performance using variable 5-nucleotide spacers in the engineered *glgC* 5' UTR.** Time-course experiments of $OD_{600}$-normalized GFP fluorescence for a) TTGGT 5-nt spacer, b) TTCAT 5-nt spacer, c) TTACA 5-nt spacer, d) TTAC 5-nt spacer.

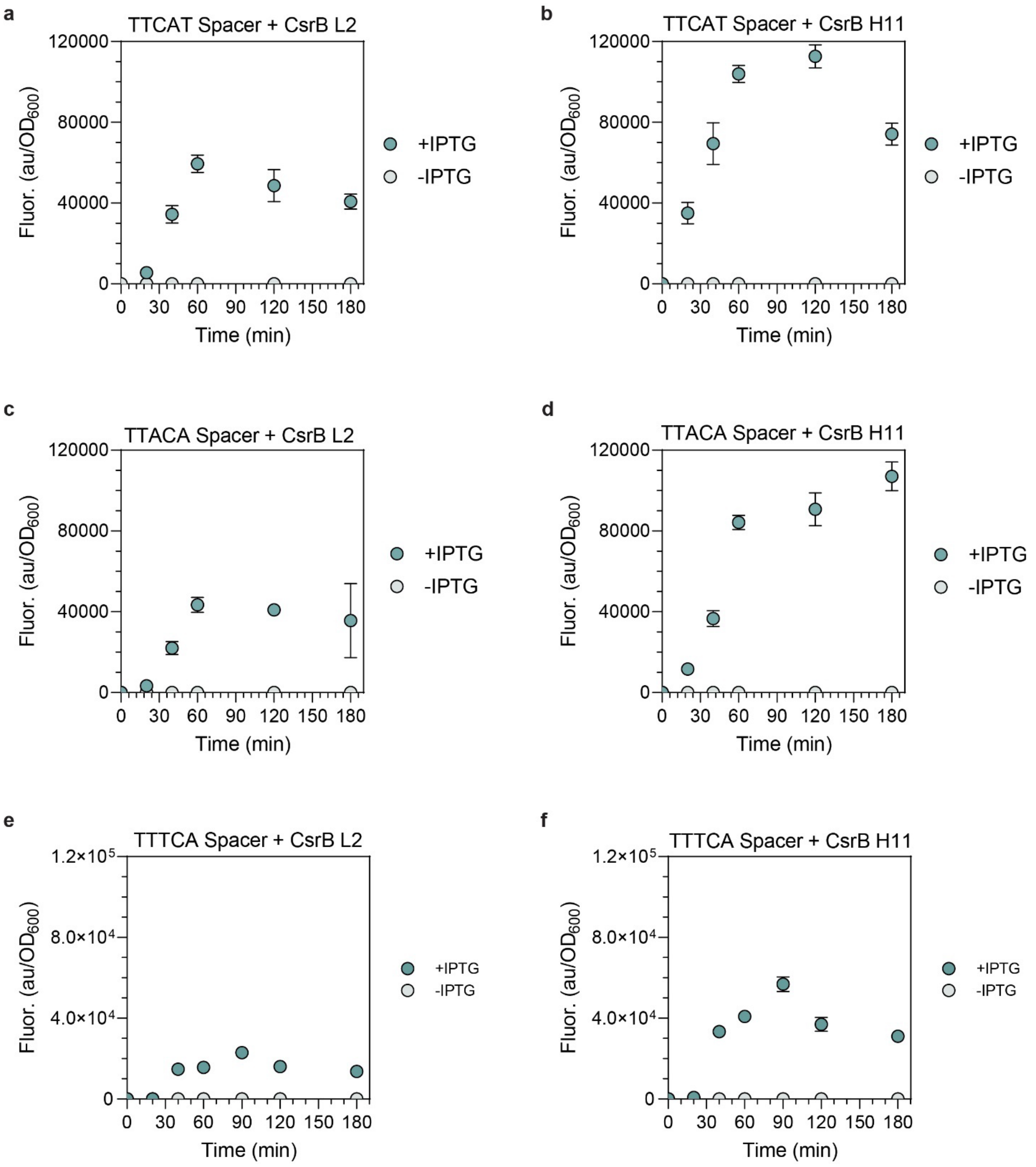

**Supplementary Figure S6. Csr-Controlled Buffer Gate Performance of all additional CsrB sRNA and 5-nt spacer combinations.** Time-course experiments of $OD_{600}$-normalized GFP fluorescence for a) TTCAT 5-nt spacer and CsrB L2, b) TTCAT 5-nt spacer and CsrB H11, c) TTACA 5-nt spacer and CsrB L2, d) TTACA 5-nt spacer and CsrB H11, e) TTTCA 5-nt spacer and CsrB L2, f) TTTCA 5-nt spacer and CsrB H11.

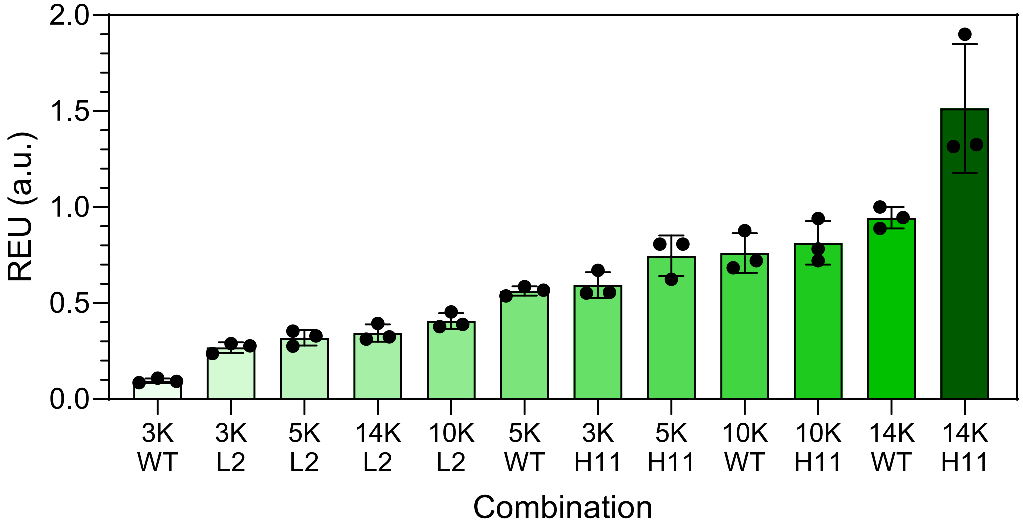

**Supplementary Figure S7. REUs of all Csr Buffer Gate combinations of *csrB* sRNAs and 5-nt RBS sequences.** REU of each Csr Buffer Gate two hours post-*csrB* induction.

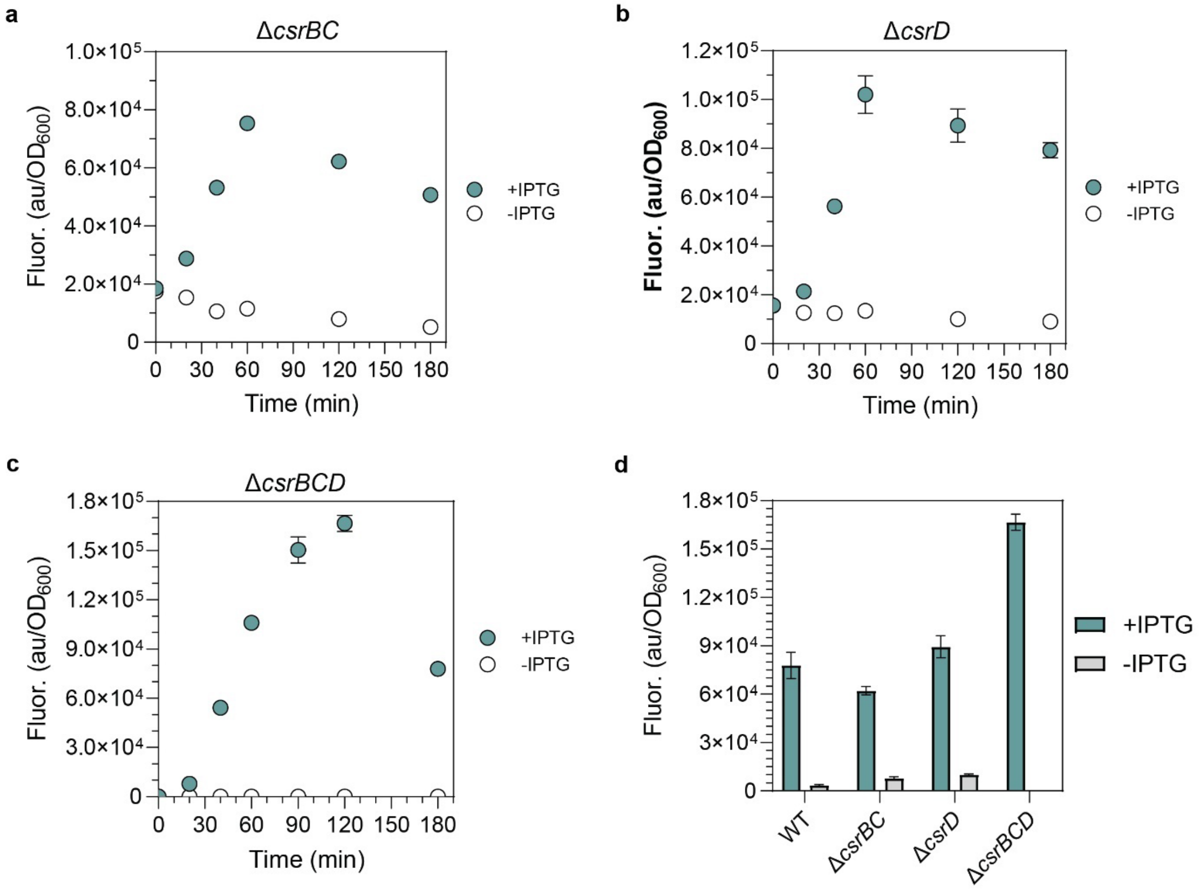

**Supplementary Figure S8. Performance of the Csr-Controlled Buffer Gate in various *csr* deletion strains.** Time-course experiments of $OD_{600}$-normalized GFP fluorescence using the original Csr-Controlled Buffer Gate construct (TTGGT 5-nt Spacer, WT CsrB) in the a) *ΔcsrB ΔcsrC* (*ΔcsrBC*) *E. coli* strain, b) *ΔcsrD E. coli* strain, and c) *ΔcsrBΔcsrCΔcsrD (ΔcsrBCD*) *E. coli* strain. d) $OD_{600}$-normalized GFP fluorescence of the original Csr-Controlled Buffer Gate in each strain 2 hours post-induction.

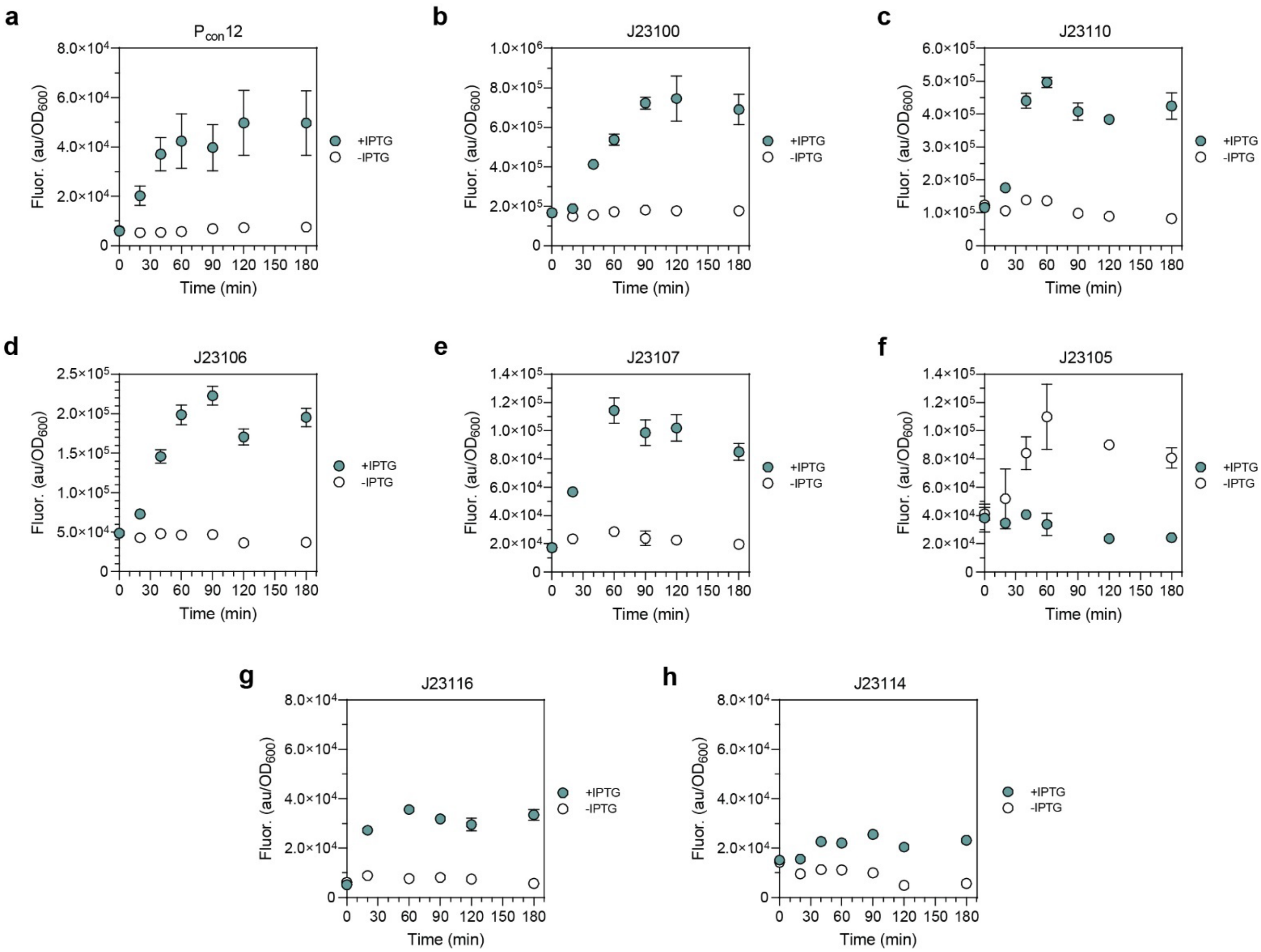

**Supplementary Figure S9. Testing Csr-Controlled Buffer Gate Performance using Anderson Promoter Library sequences.** Time-course experiments of $OD_{600}$-normalized GFP fluorescence using the original Csr-Controlled Buffer Gate construct (TTGGT 5-nt Spacer, WT CsrB) using the a) original Pcon12 promoter from Adamson et al. 2013 *PNAS*, b) Anderson Promoter J23100, c) Anderson Promoter J23110, d) Anderson Promoter J23106, e) Anderson Promoter J23107, f) Anderson Promoter J23105, g) Anderson Promoter J23116, and h) Anderson Promoter J23114.

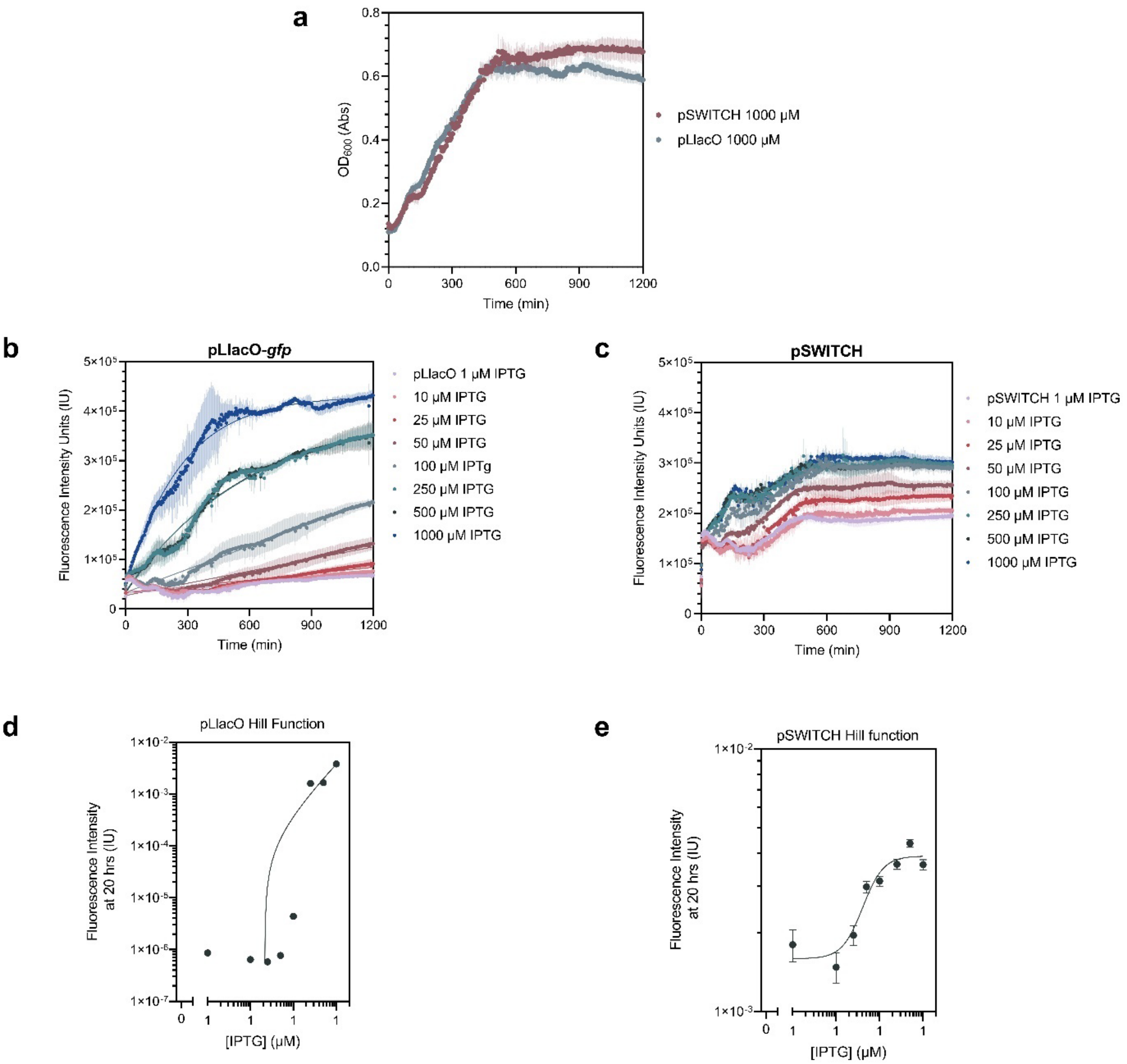

**Supplementary Figure S10. Performance of the Csr-Controlled Buffer Gate against a LacI-controlled transcriptional buffer gate in *S. oneidensis*.** a) Growth curve measured by absorbance at 600 nm of *S. oneidensis* containing either the Csr-Controlled Buffer Gate (pSWITCH) or the transcriptionally controlled Buffer Gate (placI-gfp). b) Fluorescence time-course of LacI-controlled transcriptional buffer gate regulating GFP (pLlacO-gfp) at various IPTG induction concentrations. c) Fluorescence time-course of Csr-controlled Buffer Gate (pSWITCH). d) Hill Function derived from fluorescence time-course of pLlacO-gfp construct. e) Hill Function derived from fluorescence time-course of pSWITCH construct.

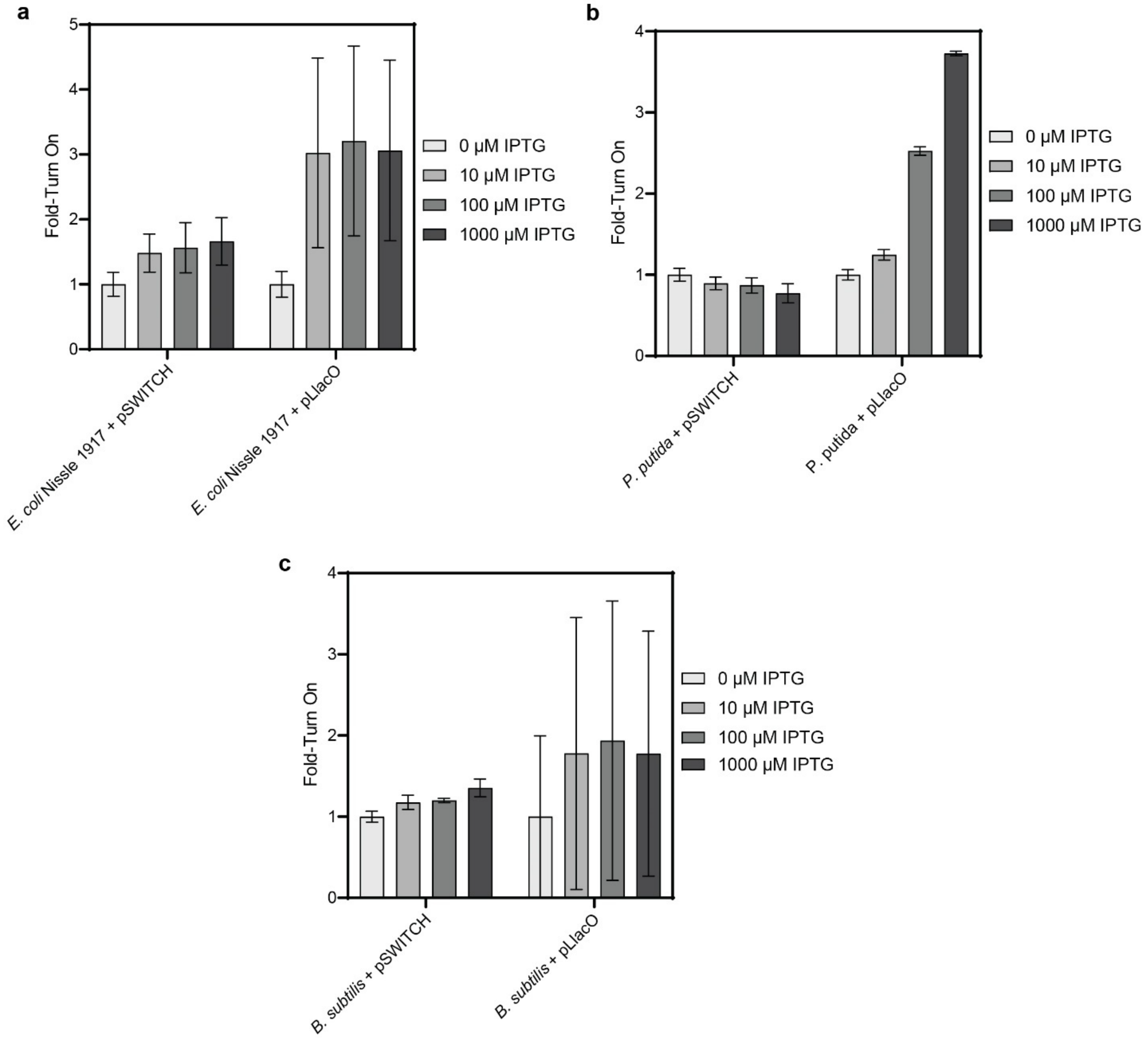

**Supplementary Figure S11. Comparing the Csr-Controlled Buffer Gate against a Lac-regulated transcriptional Buffer Gate across multiple Csr-containing bacteria.** a) Fold Turn-On of the Csr-controlled Buffer Gate (pSWITCH) and the Lac-regulated transcriptional Buffer Gate (pLlacO) at increasing IPTG induction concentrations for a) *Escherichia Coli Nissle 1917*, b) *Pseudomonas putida,* and c) *Bacillus subtilis*.

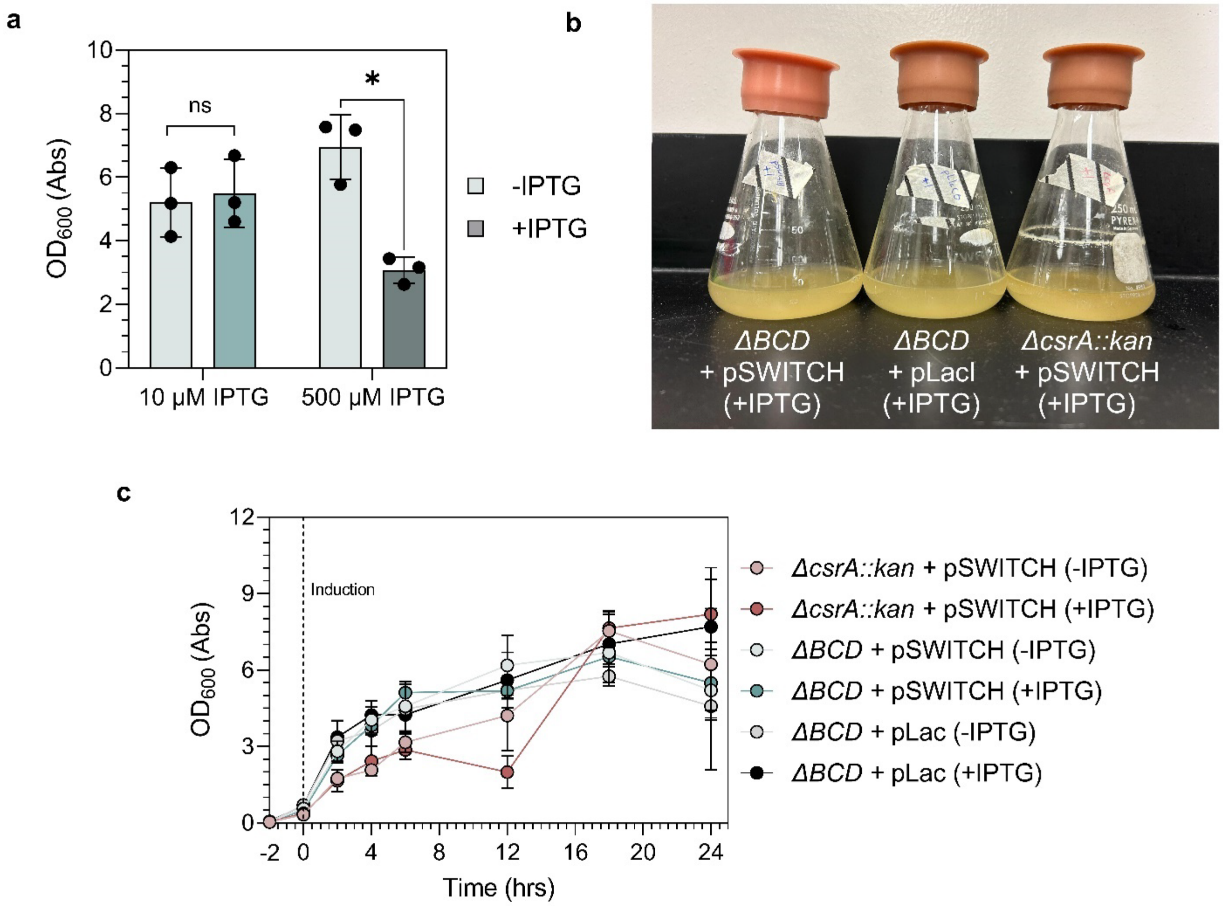

**Supplementary Figure S12. Growth Curves of Mevalonate Producing Strains** a) Final $OD_{600}$ of pCsr-Mev-containing strains induced with 10 µM IPTG and 500 µM IPTG. b) Imagine of flasks containing either the pCsr-Mev (pSWITCH) or plac-Mev (placI) regulated constructs in the Mevalonate producing strain (Δ*BCD*) or the CsrA deletion strain (*csrA::kan*). c) Growth curves of mevalonate producing cultures the pCsr-Mev construct in Δ*BCD* and *csrA::kan* strain, as well as the plac-Mev construct in the Δ*BCD* strain.

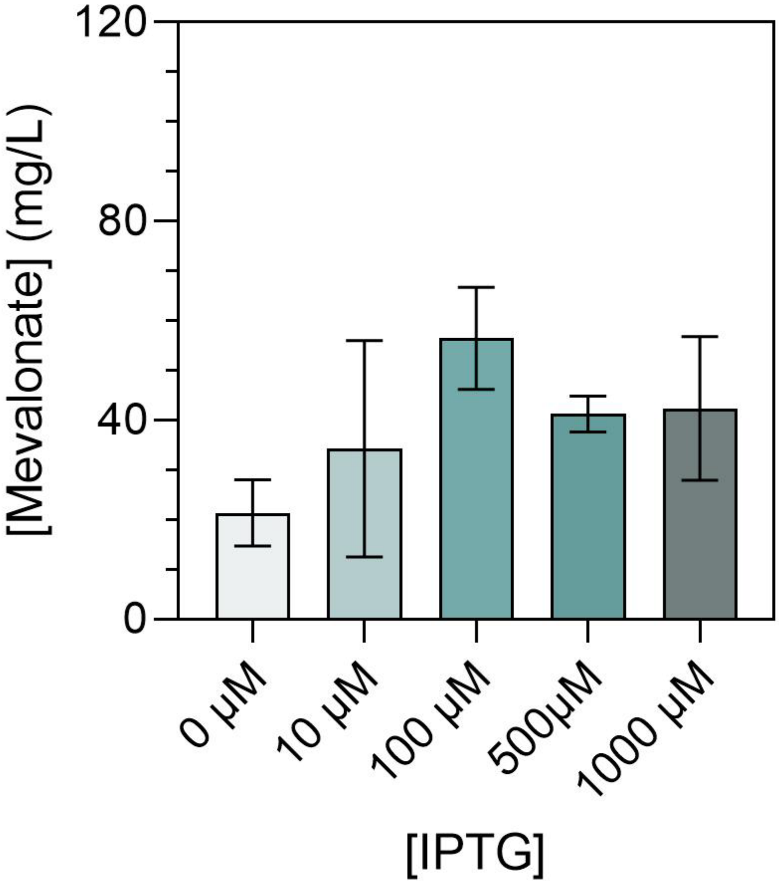

**Supplementary Figure S13. Mevalonate production from placI-Mev-containing cultures at different induction conditions.** Final mevalonate concentration from each batch of cultures induced with a different concentration of IPTG.

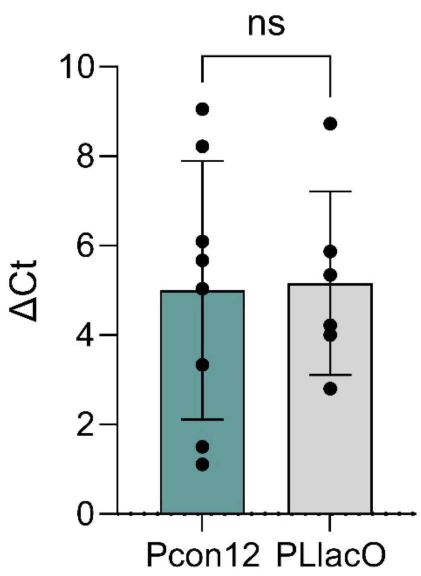

**Supplementary Figure S14. qPCR on synthetic mevalonate operon mRNA in the Csr-regulated and LacI-regulated systems.** ΔCt for the Csr-regulated system ($P_{con12}$ promoter) and the LacI-regulated system ($P_{LlacO}$). Abundance was quantified by comparing ΔCt between the mevalonate operon and the *secA* housekeeping gene.

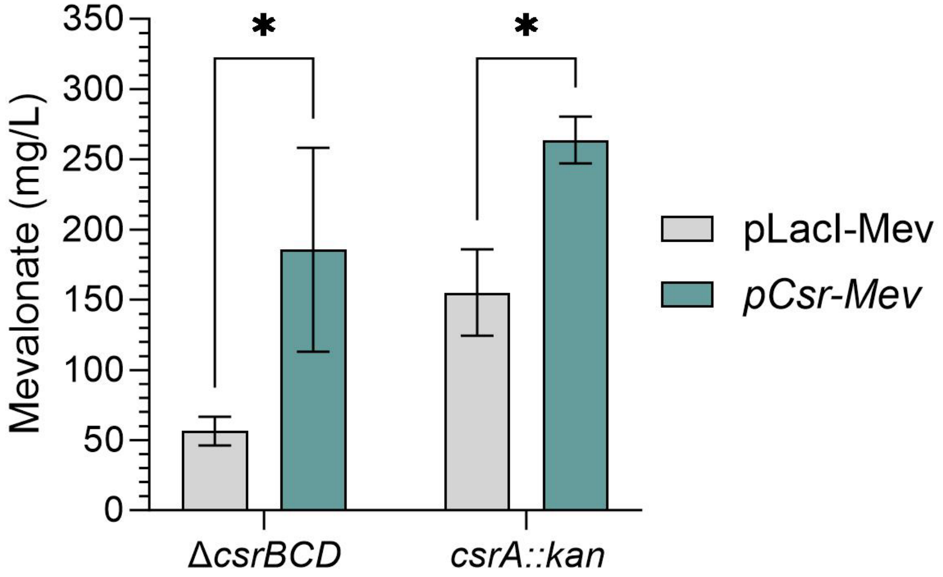

**Supplementary Figure S15. Mevalonate production in the *ΔcsrBCD* and *csrA::kan* strains for the Csr-regulated and LacI-regulated systems.** Mevalonate concentration 24 hours post-induction in the Δ*csrBCD* and *csrA::kan* strains of *E. coli.*

**Supplementary References**

1. Partipilo et al. 2022 *ACS Cent. Sci.*
2. Dundas et al. 2020 *ACS Synth. Biol.*
3. Coelho et al. 2013 *Nat. Chem. Biol.*
4. Stork et al. 2021 *Nat. Commun.*
5. Sowa et al. 2017 *NAR*
6. Hernandez-Lozada et al. 2018 *ACS Synth. Biol.*
7. Rojano-Nisimura and Simmons et al. 2023 *bioRxiv (in review at Frontiers in Molecular Biology)*
8. Meyer et al. 2019 *Nat. Commun.*
9. Leistra et al. 2017 *ACS Synth. Biol.*
